# Programmable De Novo Design of Mesoporous Protein Crystal Frameworks

**DOI:** 10.64898/2026.08.25.747085

**Authors:** Zhe Li, Shunzhi Wang, Will Sheffler, Yang Hsia, Byeongdu Lee, Greg L. Hura, Muammer Y. Yaman, Bo Liu, Ryan D. Kibler, Neville P. Bethel, David Chmielewski, Danny D. Sahtoe, Wei Yang, Hao Shen, Hanlun Jiang, Una Nattermann, Yunbing Shui, Haoheng Liu, Hannah Nguyen, Alex Kang, Justin Decarreau, Andrew J. Borst, Asim K. Bera, Banumathi Sankaran, David S. Ginger, David Baker

## Abstract

Three-dimensional protein crystals are ordered, porous macroscopic materials with potential applications in catalysis, biosensing, and biomedicine. However, most protein crystals are obtained by empirical screening, providing limited control over the lattice architecture, pore geometry or component composition that determine material function. Here, we present a modular strategy for the programmable design of highly porous, framework-like protein crystals using predefined protein-protein interactions. This strategy yielded over 30 distinct protein crystals, including single-component and multicomponent *P*2_1_3 and *I*2_1_3 lattices that grow to over 100 µm in size. Small-angle X-ray scattering and electron microscopy showed close agreement between experimental lattices and computational models. RFdiffusion-guided design generated isomorphous variants with matched lattice parameters, enabling coherent protein crystal alloys, epitaxial core–shell growth and reversible shell assembly. The designed crystals exhibit tunable mesoporous architectures, with limiting apertures of 2–18 nm, and support genetically encoded incorporation of fluorescent protein guests. These results establish a general route to programmable lattice engineering of protein crystals and position them as genetically encoded, compositionally tunable mesoporous materials.

## Main

Self-assembled protein materials hold great promise for biomedical applications due to their biocompatibility, molecular recognition capabilities, and structural programmability^1–4^. Among these, three-dimensional (3D) protein crystals form a distinct class, characterized by ordered networks of interconnected solvent channels— making them suitable for diverse applications such as nanomaterial templating, biocatalysis, biosensing and drug delivery^5–8^. While protein crystals are routinely obtained through crystallographic screening for structural biology, such crystals are typically discovered empirically and provide limited control over lattice architecture, pore geometry and component composition that determine material function^9^. Previous efforts to direct three-dimensional protein self-assembly have used a range of interaction types, including electrostatic^10^, metal-mediated^11^, DNA-hybridization^12^, and aromatic-aromatic interactions^13,14^. However, few methods have effectively leveraged designed side-chain-mediated protein–protein interactions without artificial modifications—analogous to the native contacts that stabilize conventional protein crystals—to drive macroscopic crystal self-assembly with programmed lattice architecture^15,16^. Our previous work established the feasibility of accurate protein crystal design through hierarchical assembly of protein polyhedral nanocages, but its reliance on preassembled quaternary scaffolds limited the modularity of lattice and component design^16^. An outstanding current challenge is to move beyond scaffold-dependent assembly toward modular crystal-design systems that decouple lattice geometry, interaction strength, sequence identity and composition while preserving predictable macroscopic order.

Many crystallographic space groups can be decomposed into sets of interactions within and between cyclic point group assemblies. We reasoned that recent advances in de novo protein design^17,18^ could enable the construction of 3D protein crystals by rigidly fusing cyclic symmetry-generating protein modules to interaction-directing modules that place them in orientations compatible with periodic lattice formation. We further reasoned that tuning the relative strengths and reversibility of these intra- and intercomponent interactions could enable efficient hierarchical assembly of the crystal lattice. We therefore set out to explore this approach as a modular route for generating diverse crystal geometries and enabling predictive design of protein crystal architectures.

## Results

We chose to focus on the *P*2_1_3 and *I*2_1_3 crystal space groups as design targets. Our selection was guided by three criteria: (1) the building blocks should be readily accessible through de novo protein design with high success rates; (2) the assembly should involve a minimal number of distinct protein–protein interfaces, reducing the complexity of computational design and experimental validation; and (3) the assembly pathway should proceed through a minimal number of hierarchical steps, avoiding multi-step intermediates and facilitating efficient lattice formation. For the first criterion, the *P*2_1_3 and *I*2_1_3 space groups can be geometrically deconstructed into low-symmetry cyclic oligomers^19^, which are compatible with well-established and reliable design workflows^20–25^. For the second and third criteria, each target lattice requires only two to three distinct protein-protein interfaces^19,26^ and proceeds through just two hierarchical steps: from monomer to oligomer, and from oligomer to 3D crystal. Importantly, the two space groups also provide complementary tests of the same design framework, with *P*2_1_3 enabling multicomponent assembly through heterotypic contacts between two C3 oligomeric building blocks and *I*2_1_3 enabling single-component assembly through homotypic contacts between copies of one C3 building block. Together, these characteristics make both space groups well suited for generalizable, modular design of self-assembled protein crystals.

### Modular computational design strategy

To guide our design strategy, we adopted a graph-theoretical model in which protein– protein interfaces are represented as nodes and intramolecular linkers as edges. In this scheme, the *P*2_1_3 lattice comprises two distinct C3 oligomer nodes connected via a heterodimeric interface node (Fig. 1a), whereas the *I*2_1_3 lattice consists of a single C3 oligomer node alternating with a homodimeric interface node (Extended Data Fig. 1a). In both cases, successful lattice propagation requires the C3 axes of adjacent oligomer nodes to skew at 70.5° (in *I*2_1_3, this is also defined by the C3 axes of the oligomer nodes and the C2 axes of the crystal contact nodes skew at 54.7°) (Fig. 1b, Extended Data Fig. 1b).

**Fig. 1.**
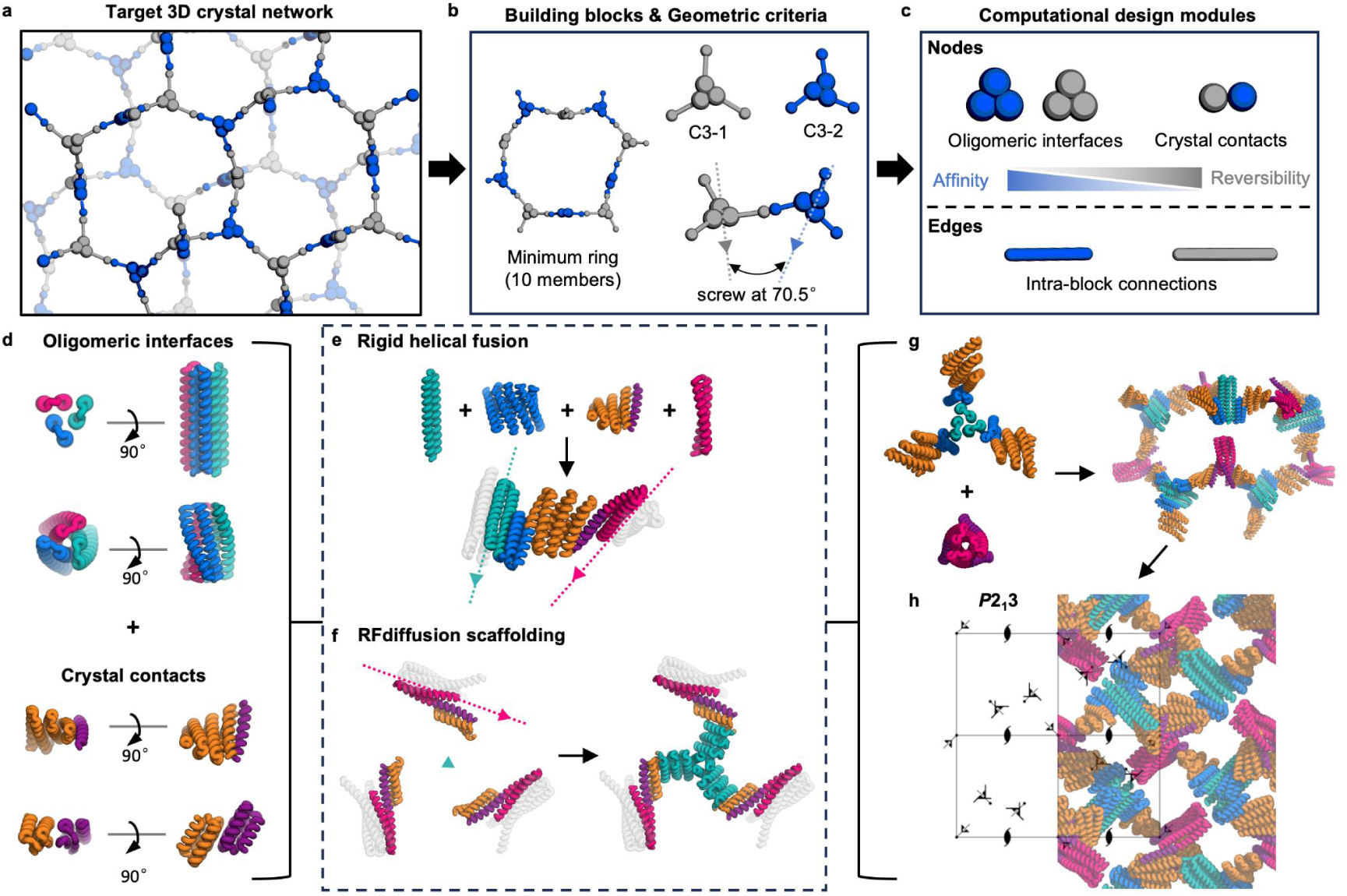
Modular computational design strategy for *P*2_1_3 space group. **a**, Schematic of the target three-dimensional crystal network, with spheres and rods as protein components, analogous to nodes and edges in a graph. **b**, Building blocks and geometric criteria for the network: two distinct protein oligomers C3-1 and C3-2 interact with their cyclic axis skewed at 70.5°, forming a minimum 10-member ring assembly (left). **c**, Computationally designed modules building up the crystal lattice: nodes represent protein–protein interfaces, including high-affinity, low-reversibility oligomeric interfaces and low-affinity, high-reversibility crystal contacts, whereas edges correspond to covalent intra-block connections. **d**, Models of the oligomeric interface modules and the crystal contact modules. **e**, Architecture-guided rigid helical fusion for building block design. **f**, RFdiffusion-based backbone generation for connecting prepositioned modules. **g**, Models of designed protein building blocks and the minimum ring assembly indicated in panel b. **h**, Model of assembled 3D protein crystal superimposed with the symmetry elements of the *P*2_1_3 unit cell. In **d-h**, modules are color-coded; dash lines and triangles indicate the three-fold cyclic axis of the corresponding modules.

We program lattice formation through three coordinated design elements: modular homo-oligomer and interface building blocks, their precise spatial positioning via designed rigid linkers, and the tuning of interfacial interaction strength to guide assembly. The C3 symmetry axes are established by pre-validated cyclic oligomers. The spatial organization and orientations of these building blocks are enforced by rigid linkers which connect to prevalidated interface modules designed to maintain accurate, reproducible propagation of the framework in three dimensions. Tuned interfacial interaction strengths promote self-assembly through a two-step hierarchical process: first, strong homotrimeric interfaces form stable C3 oligomeric building blocks; second, weaker, reversible crystal contacts—heterodimeric in *P*2_1_3 and homodimeric in *I*2_1_3—mediate oligomer–oligomer interactions to build the 3D lattice. (Fig. 1c, Extended Data Fig. 1c). These crystal contact nodes are intentionally tuned for moderate binding affinity, achieving a critical balance: strong enough to drive assembly yet weak enough to permit error correction and prevent kinetic trapping, thereby ensuring high-fidelity lattice formation.

Based on these principles, we established a robust and generalizable design workflow for self-assembling 3D protein crystals, as summarized below (Fig.1d-h, Extended Data Fig. 1d-g). Starting from homo-oligomer modules^20,22,27^ and crystal contact modules (Fig.1d, Extended Data Fig. 1d), we applied two complementary approaches to generate rigid intramolecular linkers between them, thereby producing protein building blocks with two predefined interfaces capable of lattice assembly. In the first approach (Fig.1e, Extended Data Fig. 1e), the WORMS software suite^28^ was repurposed for the *P*2_1_3 and *I*2_1_3 space groups to perform architecture-guided rigid helical fusion^28–33^, with geometric checks for lattice propagation and the generation of target crystal unit cell parameters. Previously characterized and experimentally validated de novo helical repeat proteins^34,35^ were used as linkers to ensure structural compatibility. In the second approach (Fig.1f), we used RFdiffusion-based backbone generation^23^ to directly create entirely new protein backbones between pre-positioned homo-oligomers and crystal contact modules. Unlike rigid fusion, this method allowed adaptable and fine-grained control over lattice geometry, producing a broader and more diverse set of designs. The designed connections from both methods were subsequently processed by Rosetta^36^ or ProteinMPNN^37^ for sequence design, and the resulting full-length protein monomers were evaluated and filtered using AlphaFold2^38^ (pLDDT > 90, RMSD < 1.5 Å). Selected designs were experimentally validated for 3D crystal self-assembly (Fig.1g,h, Extended Data Fig. 1f,g).

For experimental validation, synthetic genes encoding the designed proteins were cloned and expressed in *E. coli*. Proteins were purified using Ni-NTA affinity chromatography followed by size exclusion chromatography (SEC). Soluble proteins were examined for 3D self-assembly either by direct mixing or hanging-drop vapor diffusion (See Methods). Obtained protein crystals were further characterized by transmission electron microscopy (TEM), small-angle X-ray scattering (SAXS) and X-ray crystallography for structural analysis. Experimentally tested protein sequences including all validated designs were summarized in Supplementary Table S1.

### Design and validation of *P*2_1_3 crystal contact modules

We first tested whether satisfying the geometric constraints of the target lattice was sufficient to drive macroscopic crystal assembly. Our initial design campaign focused on *P*2_1_3 crystal architectures and utilized previously designed homo-oligomeric modules connected via de novo helical repeat proteins, producing single-chain building blocks with strong oligomeric interfaces at both ends. Pilot experiments revealed that crystal designs with overly strong or irreversible crystal contact modules failed to assemble into ordered lattices. Of the three *P*2_1_3 designs that we experimentally tested, two yielded soluble proteins, but both formed disordered aggregates hundreds of nanometers in size, as shown by SEC and nsTEM (Extended Data Fig. 2b, 2c). We hypothesize that effectively irreversible oligomeric interfaces caused the system to fall into kinetic traps during the initial stages of assembly, leading to the accumulation of structural errors that impede long-range lattice propagation.

To overcome this limitation, we developed a “split” design strategy to utilize weaker protein–protein interactions as the *P*2_1_3 crystal contacts, with the goal of enhancing cooperativity and reversibility (Extended Data Fig. 2a, Supplementary Fig. S1a). Single-chain constructs were split into two chains at loop regions, with the loop residues around the cut point removed. This produced two protein trimers that interact across the newly formed interface, generating the 3D lattice (Supplementary Fig. S1c). In our initial experiment, six candidate design pairs were selected based on favorable side-chain packing at the designed interface. The two protein components of each design were expressed and purified separately and then tested for their co-assembly into crystals. SEC and SDS-PAGE analysis showed that for one pair—p213-w7-6A and p213-w7-6B—both components could be independently purified and were stable in solution as single monomeric SEC peaks without higher-order aggregation (Extended Data Fig. 2b, Supplementary Fig. S1d). Strikingly, this pair successfully assembled into rhombic dodecahedral shaped crystals of 20–50 µm in size after mixing and 2-3 days of incubation in 150 mM NaCl, 25 mM Tris-HCl, pH 8.0 (Extended Data Fig. 2d). We designated this design as “*P*2_1_3-1-0”, in which the first number after the space group identifies the crystal contact design variant, and the second number denotes the specific design within that series. SAXS profiles of the crystals revealed crystal assembly unit cell parameters (*P*2_1_3 space group, 180 Å) in good agreement with our computational design model (169 Å) (Fig. 2b). Cryogenic transmission electron microscopy (CryoEM) and negative-stained transmission electron microscopy (nsEM) analyses also confirmed nanoscale ordering consistent with the designed crystal architectures (Extended Data Fig. 5a, 6a, 6b).

**Fig. 2.**
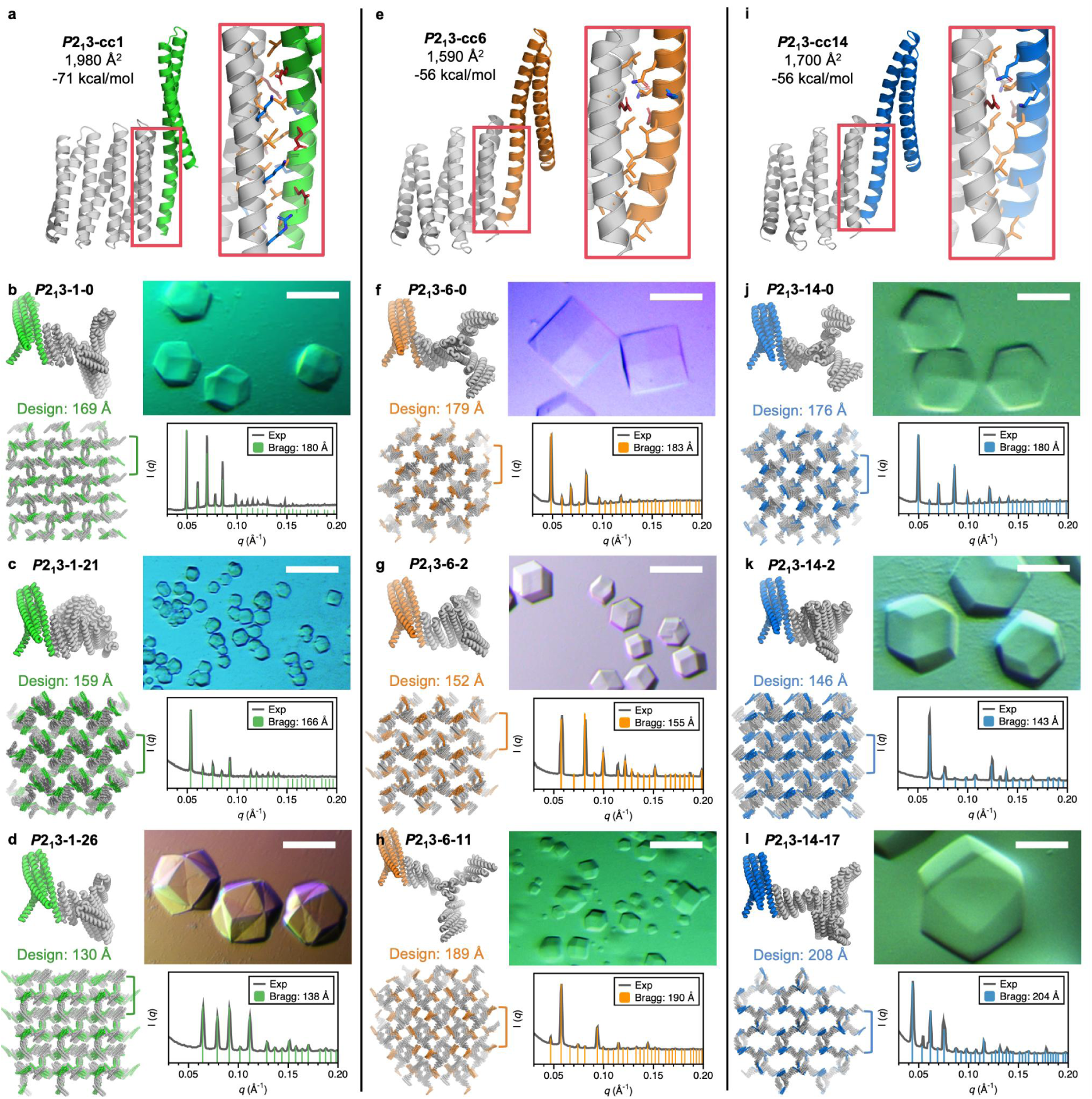
Computational design and experimental characterization of *P*2_1_3 crystals. Columns from left to right show representative crystal contacts and their corresponding derived crystal designs. **a**,**e**,**i**, Computational design models of crystal contact modules *P*2_1_3-cc1, *P*2_1_3-cc6, and *P*2_1_3-cc14. Components colored green, orange, and blue were conserved across all design variants, while the gray component was varied through protein fusion strategies. Insets on the right show side-chain interaction details at the designed protein-protein interface. **b-d**, **f-h**, **j-l**, Computational design models of the crystals assembled via *P*2_1_3-cc1, *P*2_1_3-cc6, and *P*2_1_3-cc14, respectively, alongside their experimental validation by optical microscopy and SAXS. For each crystal assembly: top left, two C3 components interacting through the designed crystal contacts; bottom left, symmetry-expanded protein crystal lattice viewed along the [100] direction with the designed unit cell parameter indicated above; top right, optical microscopy images of representative crystals. Scale bar, 100 µm; bottom right, experimental SAXS profile (Exp) fitted with theoretical Bragg diffraction peaks (Bragg), with corresponding experimental unit cell parameters shown in the legend.

We compared the *P*2_1_3-1-0 design to the other five designs that did not form crystals to identify properties of crystal contact modules appropriate for our self-assembly strategy. Among the tested designs, Rosetta metric calculations identified the *P*2_1_3-1-0 crystal interface as the weakest, with the smallest solvent-accessible surface area (SASA) and the least negative predicted interface ddG (Supplementary Fig. S1b). Closer examination of the protein–protein interface revealed distinct differences in secondary-structure interactions and side-chain packing (Supplementary Fig. S2a). The successful module displayed a unique “1+2” helix interface format, in which one helix on one side of the contact engaged with two helices on the opposing side. Compared with the more common “2+2” format, this arrangement generated a smaller crystal-contact interface. In contrast, the larger split interfaces in many “2+2” designs likely exposed more hydrophobic surface area, promoting nonspecific aggregation and poor solubility.

Building on the “1+2” helix interface format, we next designed and experimentally characterized a library of heterodimeric crystal contact modules for *P*2_1_3 crystal designs (Supplementary Fig. S2b). Starting from a set of C3 helical bundles rigidly fused to de novo helical repeat proteins, we split the proteins after the second helix and filtered the resulting constructs to retain only those adopting the “1+2” interface geometry. The C3 bundle side of the split was prioritized for experimental validation because it is shorter and involves fewer variants, enabling efficient gene synthesis and screening to identify viable assembly pairs. Of the 18 bundle designs tested, 6 displayed excellent solubility and monodisperse SEC profiles, identifying them as promising crystal contact modules for subsequent WORMS designs and crystal assembly experiments. By contrast, designs with poor solubility were excluded, since both components must be soluble to serve as effective building blocks for crystal formation. The crystal structure of one designed component, p213-14B, was determined at 2.70 Å resolution and closely matched the computational design, with an all-atom root-mean-square deviation (RMSD) of 0.35 Å. (Supplementary Fig. S3).

### Design and characterization of 2-comp *P*2_1_3 crystals

With the soluble and monodisperse crystal contact modules, *P*2_1_3 crystals were computationally designed and experimentally characterized. Protein helical bundle modules and repeat protein modules were first fused using the Rosetta helical fusion protocol to generate a C3 module library, which was then connected to the crystal contact modules using a modified WORMS software suite optimized for 3D crystal design. After Rosetta/ProteinMPNN sequence design, AlphaFold2 structural validation, and visual inspection for steric clashes between propagating components, 97 crystal designs incorporating 6 crystal contact modules were experimentally evaluated for 3D crystal self-assembly.

For most of the designed crystal components we obtained a soluble protein fraction suitable for crystallization screening, although expression levels and monodispersity varied. These components exhibited predominantly monodisperse SEC profiles, and their SAXS profiles were consistent with the corresponding design models (Extended Data Fig. 3). Crystal formation was assessed by optical microscopy, nsTEM/cryoEM, and SAXS profiles (Fig. 2, Extended Data Fig. 4-6, Supplementary Fig. S4-6). Three crystal contact modules—*P*2_1_3-cc1, *P*2_1_3-cc6, and *P*2_1_3-cc14—showed the highest success rates. Here, “cc” denotes crystal contact, and subsequent crystals are indexed accordingly (for example, *P*2_1_3-1-0 is the first crystal derived from *P*2_1_3-cc1). These interfaces feature interdigitated hydrophobic cores stabilized by peripheral polar interactions. Compared with previously designed de novo nanocage interfaces, the designed crystal-contact interfaces exhibited comparable predicted interface ddG values, similar shape-complementarity (SC) scores, and slightly larger solvent-accessible surface area (SASA) and contact molecular surface (CMS) values (Supplementary Table S2, Supplementary Fig. S7). Most assemblies formed rapidly, yielding macroscopic crystals over 50 µm in size after overnight incubation at room temperature (Supplementary Table S3). All resulting crystals exhibited quasi-rhombic dodecahedral morphologies characteristic of the cubic crystal class, consistent with the designed *P*2_1_3 lattice packing (Supplementary Fig. S8), although individual designs showed uneven facet development and surface distortions. Single-crystal X-ray crystallography yielded diffraction at low resolution (>7 Å), insufficient for atomic-level structural determination, likely due to unusual high solvent content, structural flexibility^39^ and suboptimal crystallinity, but SAXS analysis of the crystals revealed distinct diffraction peaks at low scattering vectors (q < 0.2 Å^-1^). Fitting these peaks to the Bragg reflections of *P*2_1_3 lattices showed a near-perfect match between experimental and theoretical peak positions, with fitted unit-cell parameters within 11 Å of the design models (Fig. 2, Extended Data Fig. 4, Extended Data Table 1). CryoEM and nsEM characterization also exhibited highly ordered nanoscale patterning consistent with the computational designs (Extended Data Fig. 5a, 5b, 6a-e, Supplementary Fig. S5, S9 and S10).

### Design and characterization of 1-comp *I*2_1_3 crystals

For both our validated *P*2_1_3 crystals and previously reported hierarchically docked protein crystals, the architectures required two distinct protein components to co-assemble in solution—unlike most protein crystals obtained through structural biology crystallization screening, which typically assemble from a single protein species. Achieving single-component protein crystal self-assembly would represent an important step toward biomedical translation, as it could simplify genetic encoding and delivery, reduce production and formulation complexity, and streamline downstream development^8,40^. To this end, we next focused on the design of *I*2_1_3 protein crystals, which possess higher symmetry than *P*2_1_3, requiring a homodimeric crystal contact between identical protein units rather than a heterodimeric interface. Beyond the interaction-strength constraints identified for P213 designs, the homodimeric interface in I213 crystals must remain weak and reversible to be compatible with our workflow, where assembly occurs after protein expression and purification from E. coli. This prevents premature crystallization while maintaining protein solubility before controlled assembly in vitro.

We set out to design and validate homodimeric protein–protein interfaces as modular components for *I*2_1_3 protein crystal architectures, using two complementary approaches based on helical and β-sheet edge contact designs (Fig. 3a, 3c). In the first approach (Extended Data Fig. 2e, 2f), C2 helical bundle modules, which form strong, irreversible homodimeric interfaces, were fused with de novo repeat protein modules. These fused C2 modules were subsequently connected to C3 oligomeric modules to construct the *I*2_1_3 crystal framework. To fine-tune interaction strength, the all-helical C2 interfaces were weakened by deleting selected helical heptads and reconnecting the termini through loops generated using Rosetta fragment insertion^36^, producing smaller and more reversible interfaces suitable for crystal assembly. In the second approach (Extended Data Fig. 2g, 2h), we aimed to design crystal contact modules featuring β-strand edges, which have previously been shown to confer greater reversibility and specificity. To this end, we extracted a homodimeric interface from a screened crystal structure of a de novo α/β protein^41^, and subjected it to an optimization cycle combining ProteinMPNN^37^ sequence design and AlphaFold2^38^ structure prediction to diversify and evolve both the interface geometry and interaction pattern. Across both approaches, we designed crystal contacts spanning a range of interaction strengths to identify the optimal energetic window for 3D crystal assembly.

**Fig. 3.**
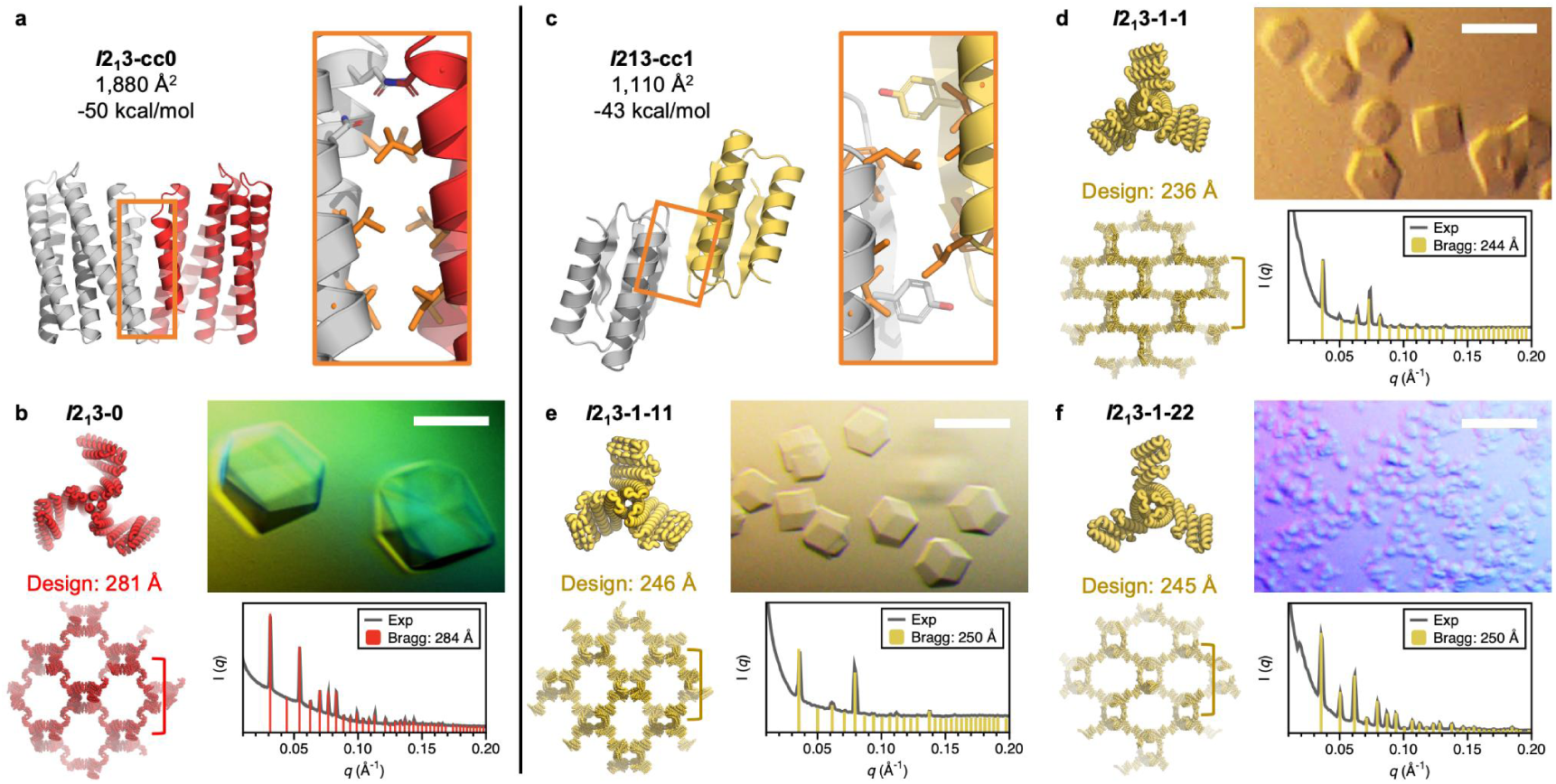
Computational design and experimental characterization of *I*2_1_3 crystals. Columns from left to right show representative crystal contacts and the corresponding derived crystal designs. **a**,**c**, Computational design models of crystal contact modules *I*2_1_3-cc0 and *I*2_1_3-cc1. Homodimeric components are shown in red or yellow against gray to highlight the interface interactions. Insets on the right show side-chain interaction details at the designed protein-protein interface. *I*2_1_3-cc0 forms an all-helical interface, whereas *I*2_1_3-cc1 features an interface with β-strand interactions located on the distal face of the model. **b**, **d-f**, Computational design models of the crystals assembled via *I*2_1_3-cc0 and *I*2_1_3-cc1, respectively, alongside their experimental validation by optical microscopy and SAXS. For each crystal assembly: top left, the C3 components constituting the crystal lattice; bottom left, symmetry-expanded protein crystal lattice viewed along the [100] direction with the designed unit cell parameter indicated above; top right, optical microscopy images of representative crystals. Scale bar, 100 µm; bottom right, experimental SAXS profile (Exp) fitted with theoretical Bragg diffraction peaks (Bragg), with corresponding experimental unit cell parameters shown in the legend.

We experimentally tested 10 designs from the first (“trim”) approach and 54 designs from the second (“extract and evolve”) approach (12, 6, and 36 designs across three successive rounds), and successfully verified four single-component protein crystal designs (Fig. 3b, 3e-g). Among the ten “trim” designs, seven produced soluble proteins as confirmed by SEC (Supplementary Fig. S11). One design, *I*2_1_3-w3-1 (also referred to as *I*2_1_3-0), showed an aggregation peak comparable in intensity to the trimer peak and yielded microcrystals in a hanging-drop crystallization setup against 0.5 M NaCl. After optimization with 10% glycerol as an additive, crystals exceeding 100 µm with rhombic dodecahedral morphology were obtained within one week. The observed rhombic dodecahedral morphology is also consistent with the designed *I*2_1_3 lattice packing (Supplementary Fig. S8). SAXS analysis of these crystals revealed Bragg peaks consistent with the theoretical *I*2_1_3 symmetry, with a fitted lattice parameter of 284 Å, closely matching the designed value of 281 Å. For the second group featuring β-strand edges, the initial 12 designs with fully contacted edges yielded no soluble proteins (Supplementary Fig. S12a, 12d). Of six designs incorporating shifted, partially contacted β-strand edges (Supplementary Fig. S12b), one crystallized rapidly under similar 0.5 M NaCl conditions after overnight incubation. Similar to the trend of screening *P*2_1_3-cc1 crystal contact module, Rosetta metric calculations identified this working crystal interface (*I*2_1_3-cc1) as the weakest among all designs (Supplementary Fig. S12c). Based on this crystal contact module, 36 additional variants were tested experimentally, from which two more designs successfully crystallized. All three designs formed rhombic dodecahedral crystals tens of micrometers in size, with one design (*I*2_1_3-1-22) even crystallizing immediately after protein elution during purification. SAXS characterization confirmed that the lattice parameters and space group symmetries were consistent with the design models (Fig. 3, Extended Data Table 1). NsEM further revealed ordered nanoscale patterns consistent with the computational designs. (Extended Data Fig. 6g-l).

### RFdiffusion generative design and characterization of *P*2_1_3 crystals

In the above design strategy based on architecture-guided rigid helical fusion, two main limitations constrain both design diversity and overall success rate. First, the design outcome is inherently dependent on a limited library of pre-validated assembly modules, constraining compositional flexibility. Second, the sampling of design candidates is further limited by the finite number of feasible fusion geometries between these modules. Together, these limitations hinder efficient exploration of the crystal architecture design space, particularly for lattices with customized parameters, and thus restrict control over crystal pore size, topology, and lattice geometry.

To overcome these constraints, we leveraged the deep learning–based generative model RFdiffusion^23^ to expand the design space. We sought to use RFdiffusion to generate isomorphous variants which maintain identical lattice parameters, and expanded variants, featuring enlarged lattice parameters. We began with previously validated *P*2_1_3 protein crystal designs (Fig. 4a). For crystals featuring the “1+2” helix crystal contact modules, the contact region was simplified to three key helices, which effectively captured the essential protein–protein interactions between assembling components. This abstraction allowed the helices to serve as the minimal functional unit for RFdiffusion-guided backbone generation and lattice remodeling. For the target protein component to be redesigned, the contact helices were maintained at the periphery of the oligomer, while RFdiffusion symmetric motif scaffolding was applied to generate new oligomeric backbones (Fig. 4b, 4c, surrounding interacting protein components are included to illustrate how this engineering influences the neighboring assembly environment.) A series of chain lengths were sampled, followed by sequence design and structure prediction (see Methods). Designs were selected for experimental validation based on prediction metrics (pLDDT > 90, RMSD < 1.5 Å), and visual inspection of overall structural geometry, packing quality, and steric compatibility.

**Fig. 4.**
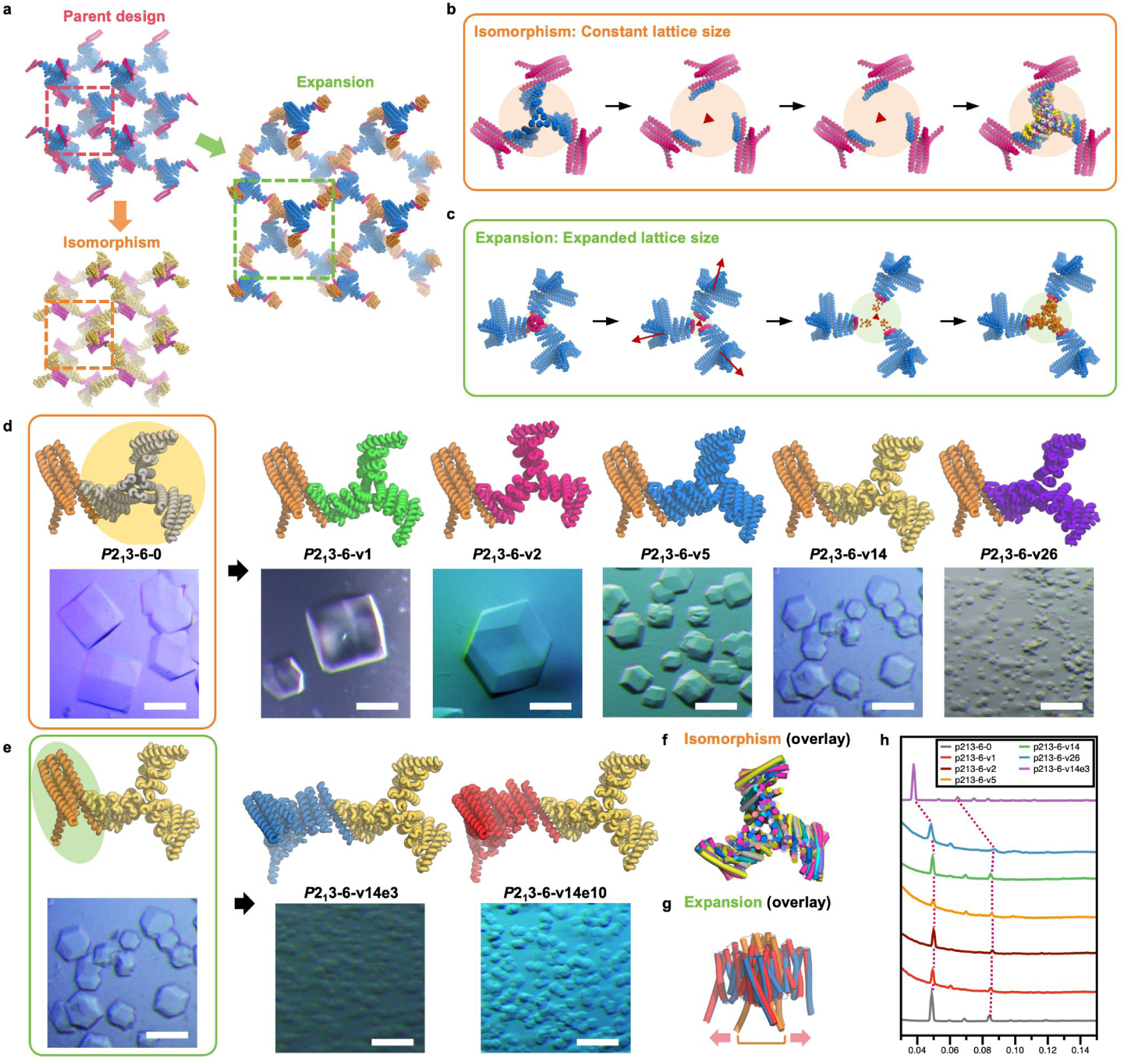
RFdiffusion generative design and experimental characterization of isomorphous and expanded *P*2_1_3 crystals. **a**, Schematic of two RFdiffusion-guided strategies for engineering an experimentally validated protein crystal. Starting from a parent crystal lattice design, either isomorphous variants with unchanged lattice parameters or expanded variants with larger lattice parameters can be generated. Unit cells are indicated by dashed boxes. **b**,**c**, Computational workflows for isomorphous redesign with constant lattice parameters (**b**) and lattice expansion with increased lattice parameters (**c**). In both workflows, protein components surrounding a single C3 symmetry axis were extracted, and RFdiffusion was used to regenerate the central C3 backbone while preserving the terminal helices that form the crystal-contact modules. This process either maintained the original component dimensions (orange highlight) or generated enlarged variants (green highlight). **d,** Isomorphous crystal lattices derived from *P*2_1_3-6-0 (first panel). Five iso-lattices with identical lattice parameters are shown with their design model and corresponding optical microscopy images. Scale bars, 100 µm. **e,** Expanded crystal lattices generated from *P*2_1_3-6-v14 (first panel). Two variants with enlarged lattice parameters are shown with their design models and the optical microscopy images. Scale bars, 50 µm. **f-g**, Overlays of RFdiffusion-generated crystal components with the original component for isomorphous (**f**) and expanded (**g**) lattices, highlighting diversified protein topology. Arrows indicate the expansion. **h**, SAXS profiles demonstrating the consistent lattice parameters across both isomorphous and expanded crystal designs.

Starting from crystal *P*2_1_3-6-0 as the parent design, we experimentally tested 28 isomorphous component A designs and 12 expanded component B designs, all generated through RFdiffusion scaffolding. All designed components were highly soluble, with 10 isomorphous and 2 expanded variants forming crystals in hanging-drop setups (Fig. 4d, 4e, Supplementary Fig. S4). Of these, six designs were further confirmed by SAXS analysis, with the expected lattice parameters consistent with either the original or expanded crystal architectures (Fig. 4h). CryoEM imaging further confirmed that the nanoscale ordering within the crystals closely matched the corresponding design models (Extended Data Fig. 5c, 5d, Supplementary Fig. S5). All crystals exhibited rhombic dodecahedral morphologies as the parent crystal, although their crystallization kinetics and optimal NaCl concentrations varied across designs. The largest crystals were typically obtained at NaCl concentrations between 1.0 and 2.5 M (Supplementary Table S3), possibly reflecting design-specific differences in geometric deviation from the target lattice, component flexibility or other interactions beyond the direct crystal-contact interfaces. These factors are further examined in the following section on assembly fidelity. Although the crystal-forming components share moderate sequence identity (34.8–48.4%) and the trimers align over most of their structures (mean aligned fraction, 0.887), they retain substantial geometric variation (RMSD, 2.66–6.06 Å; TM-score, 0.504–0.891) (Fig. 4f, 4g, Supplementary Fig. S13). We further confirmed that independently redesigned lattice components remained recombinable during crystal self-assembly, extending the modularity of bond-centric protein assembly design^42^ to ordered three-dimensional crystals and multiplicatively expanding the accessible crystal-architecture space. (Extended Data Fig. 7).

### Assembly fidelity and kinetic control in designed protein crystals

Although experimental characterization showed that many of our designed protein crystals assembled into the target lattices —supporting the robustness of our modular design strategy—the relationships between specific design parameters, resulting crystal architectures, and assembly kinetics remain incompletely understood. Clarifying these relationships is essential both for distilling generalizable design principles and for enabling predictable, application-oriented design of protein crystal materials. To address this gap, we systematically examined the dependence of assembly fidelity on backbone geometry deviation during lattice propagation, the details of inter subunit sidechain-sidechain interactions, and the kinetics of subunit association, with a focus on how these parameters influence crystal formation and growth behavior.

In our designed lattices, the closing ring number—the minimum number of building blocks required to form a closed loop—is unusually high for protein assembly frameworks based on symmetry combinations^19^. In both *P*2_1_3 and *I*2_1_3 crystals, ten copies of C3 oligomers are required to assemble with high positional and angular precision to close a ring, causing small geometric deviations to be amplified during long-range propagation. To evaluate the tolerance of the lattice to such backbone deviations, we introduced controlled perturbations by adding or removing individual helical repeats from one of the assembly components, exploiting the robust folding properties of repeat proteins. AlphaFold2 predictions confirmed that these variants folded as designed while introducing predictable angular deviations between cyclic axes. We experimentally tested extended and truncated variants derived from four validated *P*2_1_3-cc6 and *P*2_1_3-cc14 crystal designs (Supplementary Fig. S14). All extended variants failed to form well-defined crystals, instead producing stringy precipitates in the crystallization drops. This behavior is consistent with perturbed lattice assembly, although the underlying cause may involve both angle deviation and increased backbone flexibility introduced by the extension. Most trimmed variants also failed to crystallize, with only one design retaining self-assembly after removal of a single repeat unit. Even for this design, the crystal assembly showed substantially reduced nucleation, suggesting that the modification may have introduced additional assembly strain. Together, these results indicate that accurate backbone geometry is essential for lattice propagation, while the designed system retains only a limited but detectable tolerance to structural perturbation.

After establishing the importance of accurate backbone propagation, we examined how sequence variation on a fixed backbone influences crystal assembly. Using the *P*2_1_3-1-0 design, we used ProteinMPNN^37^ to redesign the sequences on the same assembly backbone, except for the two C3 homooligomeric interfaces, which were kept unchanged. From 100 sequence design runs, 17 variants validated by AlphaFold2 predictions and Rosetta metrics were tested experimentally (Supplementary Fig. S15). Although these designs differed mainly in surface charge distribution and showed improved solubility, only two retained crystal assembly under identical conditions (Extended Data Fig. 8a). Further analysis of individual interface point mutations revealed pronounced sensitivity: even small changes in hydrophobic side-chain size dramatically altered crystallization behavior (Extended Data Fig. 8a). Certain hydrophobic-to-charged substitutions at the crystal contact interface completely abolished crystal assembly, whereas specific combinations of compensatory interface mutations restored assembly and produced well-formed crystals up to 100 µm. This high sensitivity to minimal sequence changes parallels our observations in previous protein crystal design efforts^16^ and highlights the critical role of precise interface sequence tuning in directing robust 3D protein crystal assembly.

We explored the modulation of the kinetics of crystal assembly by shielding the crystal contact interface, thereby modulating the association–dissociation equilibrium between crystal components (Extended Data Fig. 8b). For the “split” interface in *P*2_1_3 crystals, shielding was introduced by extending the two-helix side of the interface with a helix derived from the original one-helix binding partner, using different trimming boundaries. Experimentally, longer extensions produced stronger shielding and progressively slowed crystallization kinetics, with extended shielding markedly reducing assembly rates and the longest shielding element eliminating crystal formation. These results demonstrate that backbone-level engineering of crystal contact modules can be used to control assembly kinetics, enabling programmable modulation of crystal self-assembly.

### Mesoporous architectures of designed protein crystals

Our designed 3D protein crystals represent a new class of porous crystalline biomaterials, and we next sought to systematically characterize their properties. We compiled and analyzed all experimental SAXS profile validated crystal designs, together with key structural parameters including molecular weight, designed and fitted lattice parameters, pore apertures, solvent content, and crystal density (Extended Data Table 1, Supplementary Fig. S16). The fitted lattice parameters closely matched the design models, deviating by less than 6%, underscoring the robustness of our computational strategy (Supplementary Fig. S16a). Two pore metrics were calculated from the design models using MAP_CHANNELS^43^: the limiting aperture, representing the smallest permeable pore dimension (identical along all crystallographic axes due to cubic symmetry), and the maximum aperture, representing the largest internal cavity within the framework. The designed crystals exhibit limiting pore apertures spanning approximately 2–18 nm, with maximum apertures extending up to 5 nm larger. For example, design *P*2_1_3-12-7 features a ∼5 nm limiting aperture and a ∼10 nm maximum cavity, highlighting the potential for permanent guest confinement within the lattice (Supplementary Fig. S17). All designs showed high solvent content (74–95%) and low material density (0.06–0.36 g/cm^3^), consistent with highly open architectures. *I*2_1_3 crystals displayed larger pore apertures and higher solvent fractions, likely reflecting the more open topology introduced by the additional C2 symmetry between building blocks. Overall, the designed crystals combine programmable nanoscale porosity, high solvent content, and low density, establishing a versatile and lightweight biomaterial platform for molecular encapsulation and functional integration.

### Multicomponent protein crystals with programmable composition and spatial organization

To evaluate the potential of our highly porous and programmable 3D protein crystals as scaffolds for functionalization, we introduced fluorescent proteins as model guest cargos (Fig. 5a, Supplementary Fig. S18). The fluorescent proteins were covalently fused via flexible GS linkers to the N-terminus of the A component in RFdiffusion-derived variants of *P*2_1_3-6-0. These engineered crystals possess similar maximum pore apertures (∼8.5 nm), sufficient to accommodate GFP-like beta-barrel fluorescent proteins (Supplementary Fig. S18a). In this design, each cage-like pore positions three fused fluorescent proteins around the A-component trimer, enabling defined multivalent display within the lattice rather than nonspecific pore loading (Supplementary Fig. S18b). Experimentally, all fusion constructs retained crystal assembly with morphologies indistinguishable from the parent designs, and bright-field optical microscopy revealed the characteristic coloration of the fused proteins within the crystalline framework (Fig. 5b, Supplementary Fig. S19).

**Fig. 5.**
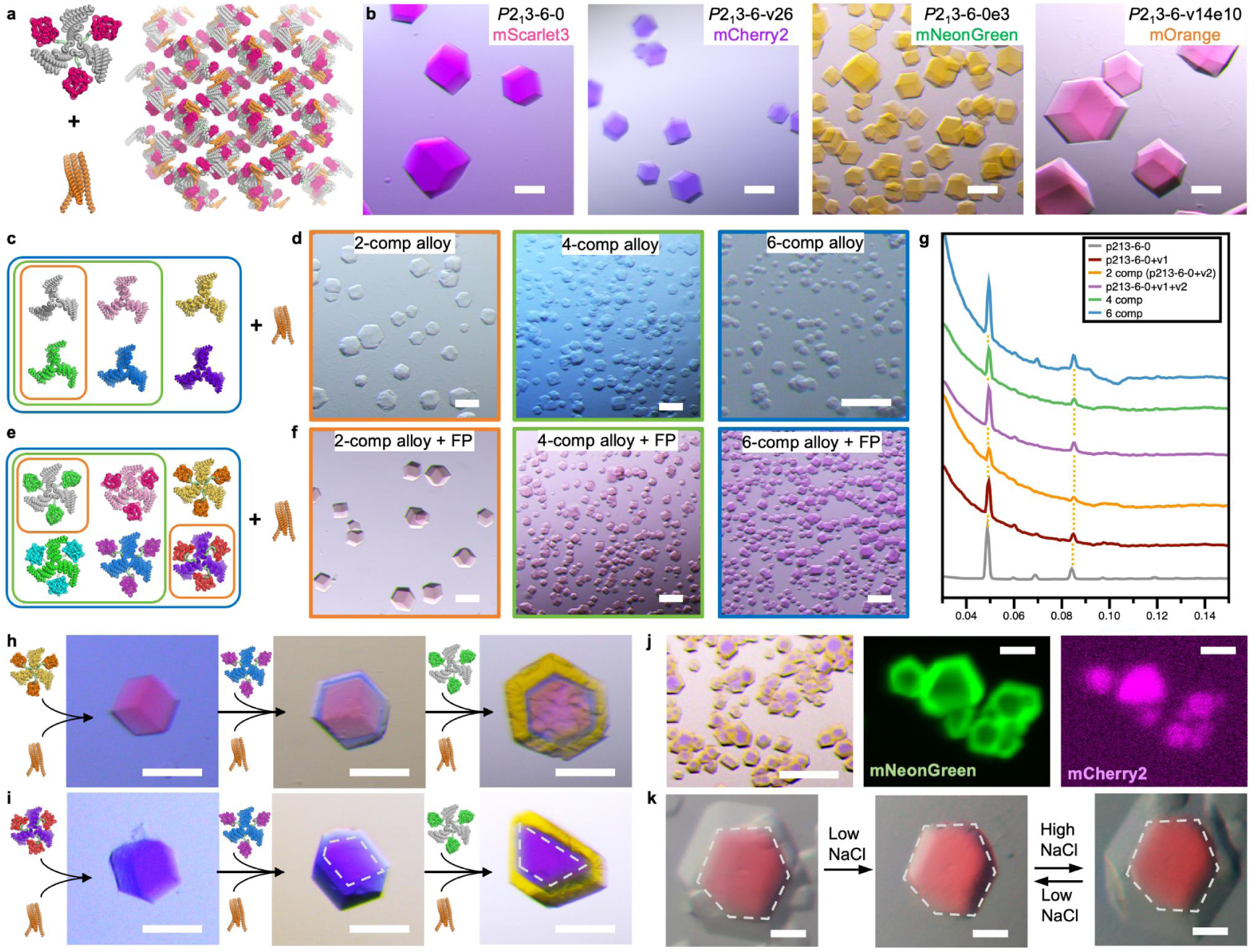
Designed protein crystals with programmable composition, spatial organization and guest immobilization. **a,** Schematic illustrating crystal assembly from A components genetically fused to guest proteins and B components forming the lattice framework. **b,** Optical microscopy images of protein crystals incorporating different GFP-like fluorescent proteins as fusion cargos. Left to right: *P*2_1_3-6-0-mScarlet2, *P*2_1_3-6-v26-mCherry2, *P*2_1_3-6-0e3-mNeonGreen, and *P*2_1_3-6-v14e10-mOrange. **c,** Assembly scheme of compositionally programmed multicomponent protein crystal alloys. Colored boxes indicate component combinations. **d**, Optical microscopy images of assembled crystals from combinations in (c). **e,** Assembly scheme of fluorescent-protein-containing crystal alloys for compositional programming of guest display. **f**, Optical microscopy images of assembled crystals from combinations in (e). **g,** SAXS profile of assembled crystal alloys; dashed lines indicate identical lattice parameters. **h-i**, Sequential assembly of spatially programmed three-layer core-shell crystals. Inner to outer layers: *P*2_1_3-6-v14-mOrange, *P*2_1_3-6-v5-mGrape3, *P*2_1_3-6-0-mNeonGreen (h); *P*2_1_3-6-v23-mCherry2, *P*2_1_3-6-v5-mGrape3, *P*2_1_3-6-0-mNeonGreen (i). **j**, Seed-mediated assembly of two-layer crystals with spatially resolved fluorescent guests: *P*2_1_3-6-v26-mCherry2 core with *P*2_1_3-6-0-mNeonGreen shell. Left to right: optical microscope image, hyperspectra fluorescence image of crystals showing emission at 556 nm and 608 nm. **k**, Reversible assembly and disassembly of specific crystal shells by changing NaCl concentration. Crystal is *P*2_1_3-6-v14-mOrange core with *P*2_1_3-6-v1 shell. Scale bars: left two images in j, 20 µm; all others, 50 µm.

At the self-assembly level, we further advanced our system toward increased architectural complexity by generating protein crystals with programmable composition and spatial organization. Up to six lattice components bearing distinct fluorescent protein fusions—mCitrine, mScarlet3, mNeonGreen, mOrange, mCherry2 and mGrape3—co-assembled into a single crystalline lattice despite differences in structure and sequence (Fig. 5c-k). We refer to these materials as protein crystal “alloys”, by analogy to inorganic alloys in which different atomic species with compatible sizes co-occupy equivalent sites within a single crystalline lattice. In our system, compositionally distinct but isomorphous protein components are integrated into one coherent protein crystal, rather than simply forming separate epitaxial layers of a single scaffold^44,45^. As proof of concept, we explored two complementary modes of programming: compositional programming through random mixed crystal alloys and spatial programming through hierarchical core–shell crystals.

For random mixed crystals, we co-assembled 2, 4, and 6 lattice components with and without fluorescent protein incorporation (Fig. 5c-g). The resulting fluorescent-protein-containing crystals exhibited homogeneous coloration under bright-field microscopy (Fig. 5f), and fluorescence microscopy and hyperspectral imaging confirmed uniform fluorophore distribution throughout the lattice and incorporation of all designed components by resolving the characteristic emission wavelengths of the constituent fluorescent protein fusions (Supplementary Figs. S21–S23). SAXS measurements further showed preserved crystallinity and identical lattice parameters across all mixed assemblies, indicating coherent co-assembly without observable lattice distortion (Fig. 5g). We next used RFdiffusion-generated isomorphous designs with identical lattice parameters to create hierarchical core–shell crystals through a macroseeding-like process. Because different isomorphous variants retained epitaxial compatibility but exhibited distinct NaCl-dependent stability windows (Supplementary Table S3), pre-formed core crystals could be transferred into shell-component solutions for selective epitaxial overgrowth while minimizing core dissolution. Fluorescent protein cargos were incorporated into different layers to visualize spatial organization, and multiple core–shell combinations formed well-defined layered assemblies with sharp boundaries and uniform fluorophore distribution within each layer (Fig. 5h-j). The shell layers could also be reversibly disassembled and reassembled for more than 4 cycles by sequentially decreasing and increasing ionic strength (Fig. 5k, Extended Data Fig. 9, Supplementary Fig. S20). Together, these alloyed and layered protein crystals demonstrate programmable compositional and spatial integration within a single crystalline framework, enabling precise three-dimensional control over structure and function in protein-based materials and providing opportunities to build on recent advances in designed protein crystals for mesoporous material synthesis^45^ and intracellular recording^46^.

## Conclusions

Our approach establishes a modular and generalizable framework for the de novo design of three-dimensional protein crystals, advancing from scaffold-dependent assembly to programmable lattice engineering through symmetry-guided building-block fusion and hierarchical interaction design. By integrating cyclic symmetry-generating building blocks with interaction-directing modules of controlled geometry, strength and reversibility, we encode lattice architecture, pore structure, component identity and functional composition directly in amino acid sequence. This strategy enables high-fidelity hierarchical assembly and, when combined with RFdiffusion-guided generation, expands the accessible design space to isomorphous, expanded and multicomponent crystal architectures. The resulting lattices exhibit programmable mesoporosity, with limiting apertures of 2–18 nm, high solvent content and close agreement with computational models, and support spatially defined guest incorporation, coherent protein crystal alloys and reversible core–shell organization within a single crystalline framework. By transforming protein crystallization from empirical discovery into predictive and programmable lattice design encoded by amino acid sequence, this work establishes designed protein crystals as genetically encoded, compositionally tunable mesoporous materials. These architectured protein frameworks offer new opportunities for molecular confinement, multivalent biocatalysis, targeted therapeutic delivery, responsive biosensing, and synthetic cellular scaffolds, bridging precise molecular design with functional biomaterials for biomedical and nanotechnology applications.

## Author contributions

Conceptualization: U.N., Y.H, Z.L. and D.B.; Methodology: Z.L., S.W., W.S. and D.B.; Investigation: Z.L., B.L., Y.S., H.L., U.N.; Building blocks: Y.H., R.D.K., N.P.B., D.D.S., W.Y.; Design protocols: Z.L., W.S., Y.H., R.D.K., D.D.S., H.J.; SAXS: Z.L., S.W., B.L. and G.L.H.; CryoEM: Z.L., S.W., N.P. B., D.C., A.J.B.; Fluorescence imaging: Z.L., M.Y.Y., D.S.G., H. S., J.D.; X-ray Crystallography: Z.L., H.N., A.K., A.K.B. and B.S.; Visualization: Z.L., S.W., W.Y. and D.B.; Funding acquisition: Z.L., D.B.; Supervision: D.B.; Writing: Z.L., S.W. and D.B.

## Acknowledgement

We thank D. Oberthür, J. Sprenger, E. Scheer, B. Klopprogge, A. Kiene, V. Kremling and H. Chapman at Center for Free-Electron Laser Science, Hamburg, Germany, for helpful discussions, crystal-screening investigations and diffraction characterization on free-electron lasers. We thank D.D. Sanctis and J. Orlans at the European Synchrotron Radiation Facility for the help with crystal diffraction characterization. We also thank J. H. Pereira and P. Zwart at Lawrence Berkeley National Laboratory, for helpful discussions and crystal-screening investigations. We thank S. Dickinson at University of Washington for help with cryoEM sample preparation and screening. We thank Y. Kipnis at University of Washington for helpful discussions and investigations of enzyme immobilization. This work was supported with funds provided by National Natural Science Foundation of China (Grant No. 32471386), Shenzhen Medical Research Fund (Grant No. A2403065), Shenzhen Science and Technology Program (Grant No. 20250603133314001), Guangdong Provincial Key Laboratory of Advanced Biomaterials (2022B1212010003), and Special Funds for the Cultivation of Guangdong College Students’ Scientific and Technological Innovation (“Climbing Program” Special Funds, Project No. pdjh2026c21306). This work was supported with funds provided by Howard Hughes Medical Institute. We acknowledge the support of the Center for Computational Science and Engineering at SUSTech. We acknowledge the assistance of SUSTech Core Research Facilities. This work was supported by We thank staff at the Advanced Photon Source (APS) Northeastern Collaborative Access Team beamlines, APS beamline 24-ID-C and the APS 12-ID-E beamline, which are funded by the National Institute of General Medical Sciences from the National Institutes of Health (P30 GM124165). The Eiger 16M detector on 24-ID-E is funded by a NIH-ORIP HEI grant (S10OD021527). We also thank staff at the APS 12-ID-B beamline for SAXS characterization. This research used resources of the Advanced Photon Source, a U.S. Department of Energy (DOE) Office of Science User Facility operated for the DOE Office of Science by Argonne National Laboratory under Contract No. DE-AC02-06CH11357. We also want to thank the Advanced Light Source (ALS) SIBYLS Beamline 12.3.1 at Lawrence Berkeley National Laboratory. This research used the Advanced Light Source resources, a DOE Office of Science User Facility under contract no. DE-AC02-05CH11231. The ALS-ENABLE beamlines are supported by the National Institutes of Health, National Institute of General Medical Sciences, grant P30 GM124169-01. The Berkeley Center for Structural Biology is supported in part by the National Institutes of Health (NIH), National Institute of General Medical Sciences, and the Howard Hughes Medical Institute. We thank the staff members of BL19U2 beamline (https://cstr.cn/31129.02.NFPS.BL19U2) at the National Facility for Protein Science in Shanghai (https://cstr.cn/31129.02.NFPS), for providing technical support and assistance in data collection and analysis.

## Methods

### Protein building block library preparation

Our initial protein building-block library comprised cyclic oligomers with C2 or C3 symmetry generated using a revised HelixFuse protocol^28,32^, which incorporates ProteinMPNN^37^ for sequence design and AlphaFold2^38^ for design validation. To construct the backbones, twofold- or threefold-symmetric helical bundles were combinatorially fused to a set of de novo repeat proteins by superimposing and overlapping residues within their terminal helices. ProteinMPNN was then used to redesign residues located within 6 Å of each newly formed fusion interface. Designs were filtered using Rosetta-based metrics, including interface shape-complementarity score greater than 0.55, and AlphaFold2 predictions, requiring a single-chain pLDDT score greater than 90 and a backbone RMSD of less than 1.5 Å relative to the design model. At a later stage, protein crystal-contact modules were incorporated into the library. For *P*2_1_3 crystals, selected C3 building blocks from the original library were reclassified as separable crystal contact modules if their constituent “split” interfaces had been experimentally validated in the target lattice. For *I*2_1_3 crystals, crystal contact modules by both “trim” and “extract” approaches were introduced.

### WORMS software suite for *I*2_1_3 and *P*2_1_3 space group symmetry

WORMS was previously developed for architecture-guided design of protein assemblies by rigid fusion of modular protein building blocks^28^. In the original workflow, each building block is treated as a rigid body with defined entry and exit splice frames, and candidate assemblies are generated by enumerating compatible combinations of building blocks and helical splice positions. The rigid-body transforms associated with individual splices are combined by matrix multiplication to evaluate whether each candidate satisfies a user-specified target architecture. Candidate models are filtered during sampling using geometric and structural criteria, including splice RMSD, steric clashes, architectural tolerance, residue-contact requirements around the fusion junction and redundancy, before full-atom models are generated for sequence design.

To adapt WORMS for the design of periodically repeating protein crystals, we implemented additional crystal-symmetry criteria for the cubic space groups *I*2_1_3 and *P*2_1_3; the code is available at https://github.com/willsheffler/worms/tree/unit_tests. In contrast to the closed cyclic, dihedral and point-group architectures targeted in the original WORMS workflow, crystal architectures require the correct relative orientation and translational placement of crystallographic symmetry axes within a unit cell. We therefore extended the WORMS geometric criteria to constrain candidate fusions according to space-group-specific relationships between symmetry axes. For *P*2_1_3 designs, compatible building-block combinations were sampled by matching two C3 building-block axes to the corresponding crystallographic threefold axes, which should skew at 70.5°. For *I*2_1_3 designs, C3 and C2 building-block axes were constrained to match the target axes skew at 54.7°.

For each candidate satisfying the crystal-symmetry criterion, the rigid-body arrangement was aligned to the target crystallographic frame, the lattice constant was inferred from the spacing between the matched symmetry axes, and the corresponding space-group label and unit-cell parameters were assigned to the output model. Candidate crystal designs were further filtered using the standard WORMS splice-geometry, contact, clash and score filters, together with user-defined limits on unit-cell dimensions.

### Crystal backbone generation by WORMS

For WORMS-based backbone generation by helical fusion, the *P*2_1_3 architecture was specified as a fusion between two C3 building blocks, whereas the *I*2_1_3 architecture was specified as a fusion between one C3 and one C2 building block. For the initial designs, in which no predefined crystal contact modules were used, C3 and C2 oligomers generated by HelixFuse were supplied as the WORMS building-block databases. At a later stage, experimentally validated crystal contact modules were added to these libraries, either as heterodimeric contacts embedded within C3 building blocks or as homodimeric C2 building blocks.

Unlike previously implemented point-group architectures^28^, which require the relevant symmetry axes to intersect, propagation of the *P*2_1_3 and *I*2_1_3 crystal lattices requires the corresponding cyclic symmetry axes to adopt defined skewed arrangements. Because these constraints can be satisfied over a substantially larger conformational search space, we applied a stringent off-architecture angular tolerance of 0.1 to preferentially retain candidates with high-quality fusion geometries. Detailed WORMS settings are provided in the corresponding flag files included in the Supplementary Data. Before sequence design, the asymmetric units of the crystal backbones generated by WORMS were visually inspected following symmetry expansion in PyMOL^47^ to identify and exclude models with potential backbone clashes between symmetry-related building blocks.

### Crystal backbone generation by RFdiffusion

Starting from experimentally validated crystal designs, we generated both isomorphous variants that retained the original lattice parameters and expanded-lattice variants using symmetry-aware motif scaffolding with the generative diffusion model RFdiffusion^23^. During backbone generation, the peripheral helices involved in the validated crystal contacts were fixed as structural motifs, whereas the intervening regions were regenerated by RFdiffusion to reconnect these helices while preserving the lattice-forming interactions.

For design *P*2_1_3-6-0A, sequence lengths ranging from 120 to 165 residues were sampled, with 100 diffusion trajectories generated for each target length. To generate expanded-lattice variants of *P*2_1_3-6-0B, the peripheral helix forming the crystal contact was retained together with a trimmed segment of the adjacent helix (Supplementary Data file). The single-chain component was displaced by 25 Å along the x axis relative to its position in the original model to increase the lattice spacing, and RFdiffusion was used to generate new backbones connecting the retained motifs. Sequence lengths ranging from 130 to 175 residues were sampled, with 100 diffusion trajectories generated for each target length.

### Sequence design and AlphaFold prediction of crystal components

Residues modified during WORMS-based backbone generation were initially redesigned using Rosetta^36^; in later design rounds, sequence design was performed using ProteinMPNN^37^. Residues within backbones generated by RFdiffusion were designed exclusively with ProteinMPNN. For each backbone, five sequences were generated at a sampling temperature of 0.1.

The resulting sequences were evaluated for monomer folding and oligomeric assembly using Superfold (github.com/rdkibler/superfold), a convenience wrapper for AlphaFold2^38^, with the initial-structure-guess protocol enabled^48^. Designs were retained if the predicted structures had a pLDDT score greater than 90, a backbone RMSD of less than 1.5 Å relative to the design model, and a tolerance value below 0.2. The retained models were subsequently inspected visually for backbone length or flexibility, steric clashes, and other structural defects before candidates were selected for experimental characterization.

### “Split” strategy for generating crystal contact modules

For WORMS-generated backbones with *P*2_1_3 symmetry, the fusion chain formed from two C3-symmetric modules was initially split at each structurally permissible loop. Loop regions were identified by visual inspection in PyMOL and removed in their entirety, after which the remaining residues were renumbered as separate chains. In subsequent designs, we preferentially generated “1+2 split” interfaces, in which a single helix on one side of the interface packs against two helices on the opposing side. The corresponding split sites were typically positioned after the HelixFuse junction, within a protruding helix that connected adjacent repeat-protein modules.

Following chain splitting, the newly generated interfaces were evaluated using Rosetta and retained if they had a shape-complementarity score greater than 0.6, a predicted change in Gibbs free energy upon complex formation (ΔΔG) below −20 kcal/mol. The remaining models were visually inspected for hydrophobic packing, and interfaces containing at least two pairs of well-packed, interdigitating hydrophobic side chains across the interface were selected.

### “Trim” strategy for generating crystal contact modules

For WORMS-generated backbones with *I*2_1_3 symmetry, the C2-symmetric modules were shortened to reduce the extent of their original helical contacts. The resulting chain termini were reconnected by newly modeled loops using the Rosetta ConnectChainsMover, followed by sequence redesign of the modified regions. The redesigned models were subsequently evaluated and filtered using AlphaFold2 predictions.

### “Extract and evolve” strategy for generating crystal contact modules

To generate weak, reversible homodimeric crystal-contact modules for *I*2_1_3 symmetry, we extracted protein–protein interfaces containing edge β-strands from either existing heterodimeric complexes (LHD101)^41^ or crystal contacts from screened crystals of de novo α/β proteins (PDB ID: 6WMK, contacts between chain B and D). These interfaces were then subjected to one to three iterative rounds of ProteinMPNN sequence design and AlphaFold2 structure prediction. For both heterodimeric and homodimeric starting structures, the sequences of the two interacting chains were tied during ProteinMPNN design. During each iteration, we selected high-confidence C2-symmetric AlphaFold2 predictions that retained the β-sheet-mediated interaction while exhibiting varying degrees of registry displacement between the interacting β-strand edges. The resulting AlphaFold2 models were subsequently used as C2-symmetric input modules for WORMS-based generation of *I*2_1_3 crystal backbones.

### Protein expression and purification

Synthetic genes were optimized for *E. coli* expression and purchased from IDT (Integrated DNA Technologies) as plasmids in pET29b(+) vector with a hexahistidine affinity tag. Plasmids were cloned into BL21* (DE3) *E. coli* competent cells (Invitrogen). Single colonies from agar plate with 100 mg/L kanamycin were inoculated in 50 mL of Studier autoinduction media^49^, and the expression continued at 37 °C for over 24 hours. The cells were harvested by centrifugation at 4000 g for 10 min, and resuspended in a 35 mL lysis buffer of 300 mM NaCl, 25 mM Tris pH 8.0 and 1 mM PMSF. After lysis by sonication and centrifugation at 14000 g for 45 min, the supernatant was purified by Ni^2+^ immobilized metal affinity chromatography (IMAC) with Ni-NTA Superflow resins (Qiagen). Resins with bound cell lysate were washed with 10 mL (bed volume 1 mL) of washing buffer (300 mM NaCl, 25 mM Tris pH 8.0, 60 mM imidazole). The resins were first pre-eluted with 0.5 mL of elution buffer (300 mM NaCl, 25 mM Tris pH 8.0, 300 mM imidazole). The bound protein was then eluted with the same buffer, and 2 ml fractions were collected. Both soluble fractions and full cell culture were checked by SDS-PAGE. Soluble designs were further purified by size exclusion chromatography (SEC). Concentrated samples were run in 150 mM NaCl, 25 mM Tris pH 8.0 on a Superose 6 Increase 10/300 gel filtration column (Cytiva). SEC-purified designs were concentrated by 10K concentrators (Amicon) and quantified by UV absorbance at 280 nm before further assembly and characterization.

### Crystallization of designs

For simplicity, protein concentrations in all crystallization experiments are reported in monomer equivalents. For example, 25 µM component A denotes a monomer-equivalent concentration of 25 µM; if component A forms a C3-symmetric trimer, this corresponds to an oligomer concentration of 25/3 = 8.3 µM.

Because the crystals were designed to assemble without requiring specific precipitants or additives, initial crystallization screens were performed under NaCl-based conditions. For preliminary tests, the two components of *P*2_1_3 designs were mixed at equal monomer-equivalent concentrations, whereas the single components of *I*2_1_3 designs were tested individually. Samples were incubated in 150 mM NaCl and 25 mM Tris-HCl, pH 8.0, overnight or for several days to allow crystal assembly. Designs that did not crystallize under these conditions were screened by hanging-drop vapor diffusion using 24-well VDXm crystallization plates (Hampton Research). For each crystal design, a 4 µl drop containing protein at a monomer-equivalent concentration of 12.5, 25, 50 µM was placed on a siliconized glass coverslip and equilibrated against a reservoir containing 0.5, 1.0, 1.5, 2.0, 2.5, 3.0 or 5.0 M NaCl. Crystallization plates were monitored for up to two weeks. Conditions yielding crystals were further optimized by varying the NaCl concentration, protein concentration and additives. For example, *I*2_1_3-0 required 10% glycerol to form crystals larger than 50 µm. Designs were also evaluated by batch crystallization through direct mixing of the protein components with NaCl solutions, guided by validated hanging-drop conditions. Crystallization conditions for all experimentally validated designs are summarized in Supplementary Table S2.

For guest incorporation, fluorescent proteins were genetically fused to the N terminus of the A component of the target crystal through a flexible (GGS)_4_ linker. The fluorescent-protein-fused A components were assembled with the corresponding B components under the same conditions used for the unfused parent crystals. Successful incorporation of the fluorescent proteins was assessed by the characteristic coloration of the crystals under bright-field microscopy.

### Self-assembly of multicomponent protein crystal “alloys” with spatial control

For core–shell crystal assembly, different isomorphous designs were first grouped according to their NaCl-dependent self-assembly and stability windows, as determined from prior crystallization screening experiments (Supplementary Table S3). For example, *P*2_1_3-6-v14 remained stable above 0.5 M NaCl, *P*2_1_3-6-v1, *P*2_1_3-6-v5 and *P*2_1_3-6-v23 remained stable above 1.5 M NaCl, and *P*2_1_3-6-0 and *P*2_1_3-6-v2 remained stable above 2.5 M NaCl. These distinct stability windows enabled sequential assembly and disassembly, with the most stable crystal used as the inner core and less stable variants added as outer layers.

The inner core crystals were first assembled by hanging-drop crystallization and then washed three times in a buffer containing the same NaCl concentration as the mother liquor. For shell growth, drops containing the outer-shell components were equilibrated overnight by hanging-drop vapor diffusion against the corresponding reservoir solution. The washed core crystal was then transferred into the shell-component drop using a loop and used as a macroseed for epitaxial overgrowth. Additional shell layers were assembled in the same manner, with washing and transfer steps performed between each growth stage.

For random mixed crystal-alloy assemblies, all crystal components were mixed at equal monomer-equivalent concentrations and set up by hanging-drop crystallization. A relatively high range of NaCl concentrations was screened to identify reservoir conditions that promoted cooperative assembly of all components. In mixed assemblies, components that did not crystallize independently at lower NaCl concentrations could still be incorporated into the crystal lattice when co-assembled with components that more readily promoted crystallization. For example, the uniformly distributed 6-component crystals shown in Supplementary Fig. S23 assembled from a 1.5 M NaCl condition, which was sufficient for independent crystallization of only a subset of the constituent components.

### Negative-stained transmission electron microscopy (nsEM)

Within the hanging drop, assembled crystals were first diluted with 6 µl of the corresponding reservoir solution and then fragmented into smaller pieces under an optical microscope using cryo-loops (MiTeGen dual-thickness MicroMounts). The resulting suspension was immediately applied to negatively glow-discharged, Formvar/carbon-coated 400-mesh copper grids (Ted Pella). Samples were incubated on the grids for up to 15 min to promote crystal attachment. During incubation, the grid held in tweezers was loosely covered with the crystallization plate lid, together with tissue paper dampened with Milli-Q water, to prevent the sample from drying. The grids were gently blotted with Grade 1 filter paper (Whatman), stained with 3 µL uranyl formate, blotted again, and then stained with a second 3 µL aliquot of uranyl formate for 20 s before final blotting. Uranyl formate concentrations of either 0.75% or 2% were used, depending on the sample. Negative-stain electron microscopy screening was performed using a 120-kV Talos L120C transmission electron microscope (Thermo Fisher Scientific).

### CryoEM sample preparation and data collection

Assembled crystals were fragmented as described for negative-stain electron microscopy and applied to glow-discharged 1.2/1.3-T C-flat holey carbon grids. The grids were vitrified using a Vitrobot Mark IV with a wait time of 5 s, a blot time of 7.5 s and a blot force of 0, followed by immediate plunge-freezing in liquid ethane. Images were collected manually using SerialEM on a 200-kV Thermo Fisher Scientific Glacios transmission electron microscope equipped with a K2 Summit direct electron detector.

### SAXS of protein oligomers

Protein oligomer samples were diluted in buffer containing 150 mM NaCl, 25 mM Tris-HCl (pH 8.0) and 2% glycerol, and then concentrated using thoroughly washed 10-kDa molecular-weight-cutoff centrifugal concentrators (Amicon). The corresponding flow-through was used as the blank for buffer subtraction. Solution scattering measurements were performed at the SIBYLS 12.3.1 beamline at the ALS in the Lawrence Berkeley National Laboratory. The SIBYLS beamline parameters and collection procedures have been previously described^50,51^. The beamline has a fixed sample-to-detector distance of 2 m. Samples were placed in a transmission geometry. The monochromatic 10 keV X-ray beam converges to a rectangular 0.5 × 1 mm^2^ beam at the sample and becomes a 100 × 100 μm^2^ beam at the detector. The area detector was a Pilatus 2 M with 172 × 172 μm^2^ pixels. SAXS data from the 2D area detector were integrated using beamline-specific software to create X-ray intensity versus momentum transfer one-dimensional curves (*I*(*q*)). A series of exposures, in equal sub-second time slices, were taken of each well: 0.3 s exposures for 10 s resulting in 32 frames per sample. For each sample, data were collected for two different concentrations to test for concentration-dependent effects. ‘Low’ concentration samples were ∼1 mg ml^−1^ and ‘high’ concentration samples were ∼5 mg ml^−1^. The data were processed using the SAXS FrameSlice online server. FoXS server^52,53^ was used to compare design models with experimental scattering profiles and to calculate the quality of fit (*χ*) values. In this work, all *I*(*q*) profiles were plotted on a logarithmic scale for clarity.

### Small-angle X-ray scattering (SAXS) of crystals

Crystal assemblies in this work were confirmed primarily by SAXS. SAXS characterization was carried out at the 12-ID-B beamline of the Advanced Photon Source at Argonne National Laboratory following a literature report^54^. The wavelengths of X-rays used at 12 was 0.9322 Å (13.3 keV), and the system was calibrated using silver behenate as a standard. Two sets of slits were used to define and collimate the X-ray beam, and parasitic scattering was removed using a pinhole. Typical exposure times of between 0.1 and 0.5 s were used. The scattered X-rays were collected with a charge-coupled device (CCD) detector, and one-dimensional scattering profiles were obtained by radially averaging the images into scattering intensities, *I*(*q*), against the scattering vector magnitude, *q*. In this work, all *I*(*q*) profiles were plotted on a logarithmic scale for clarity. One-dimensional SAXS profiles were indexed following a literature report ^55,56^. Crystallographic symmetries and lattice parameters are determined by indexing powder diffraction patterns.

### Crystal structure determination

Diffraction data was collected at the Advanced Photon Source beamline on 24-ID-C. X-ray intensities and data reduction were evaluated and integrated using XDS^57^ and merged/scaled using Pointless/Aimless in the CCP4 program suite^58^. Structure determination and refinement starting phases were obtained by molecular replacement using Phaser^59^ using the designed model for the structures. Following molecular replacement, the models were improved using phenix.autobuild^60^; efforts were made to reduce model bias by using simulated annealing. Structures were refined in Phenix^60^. Model building was performed using COOT^61^. The final model was evaluated using MolProbity^62^. Data collection and refinement statistics are recorded in Supplementary Table S4. Data deposition, atomic coordinates, and structure factors reported in this paper have been deposited in the Protein Data Bank (PDB), http://www.rcsb.org/ with accession code pdb_000037TD.

### Hyperspectral fluorescence microscopy

Hyperspectral fluorescence data cubes were collected using a Photon Etc. IMA system coupled to a Nikon Ni-U upright microscope equipped with a ×60 objective lens (Nikon, numerical aperture 0.6). Samples were excited at 500 nm with an appropriate fluorescence filter set. The emitted fluorescence was passed through a tunable volume Bragg grating filter with a wavelength step size of 1-2 nm and an integration time of 1 s per step. At each wavelength, the fluorescence signal was recorded using a charge-coupled device camera (Thorlabs, 1501M-US-TE). Hyperspectral data cubes were generated by accumulating wavelength-resolved images over the spectral range of 500–850 nm. The fluorescence spectra were background corrected and processed using Photon Etc. PhySpec software (version 2.27).

### Design models, example scripts and experimental data

Design models for all experimentally validated crystals, representative scripts for individual design steps, and experimental characterization data are available through Zenodo^63^.

---

**Extended Data Fig. 1.**
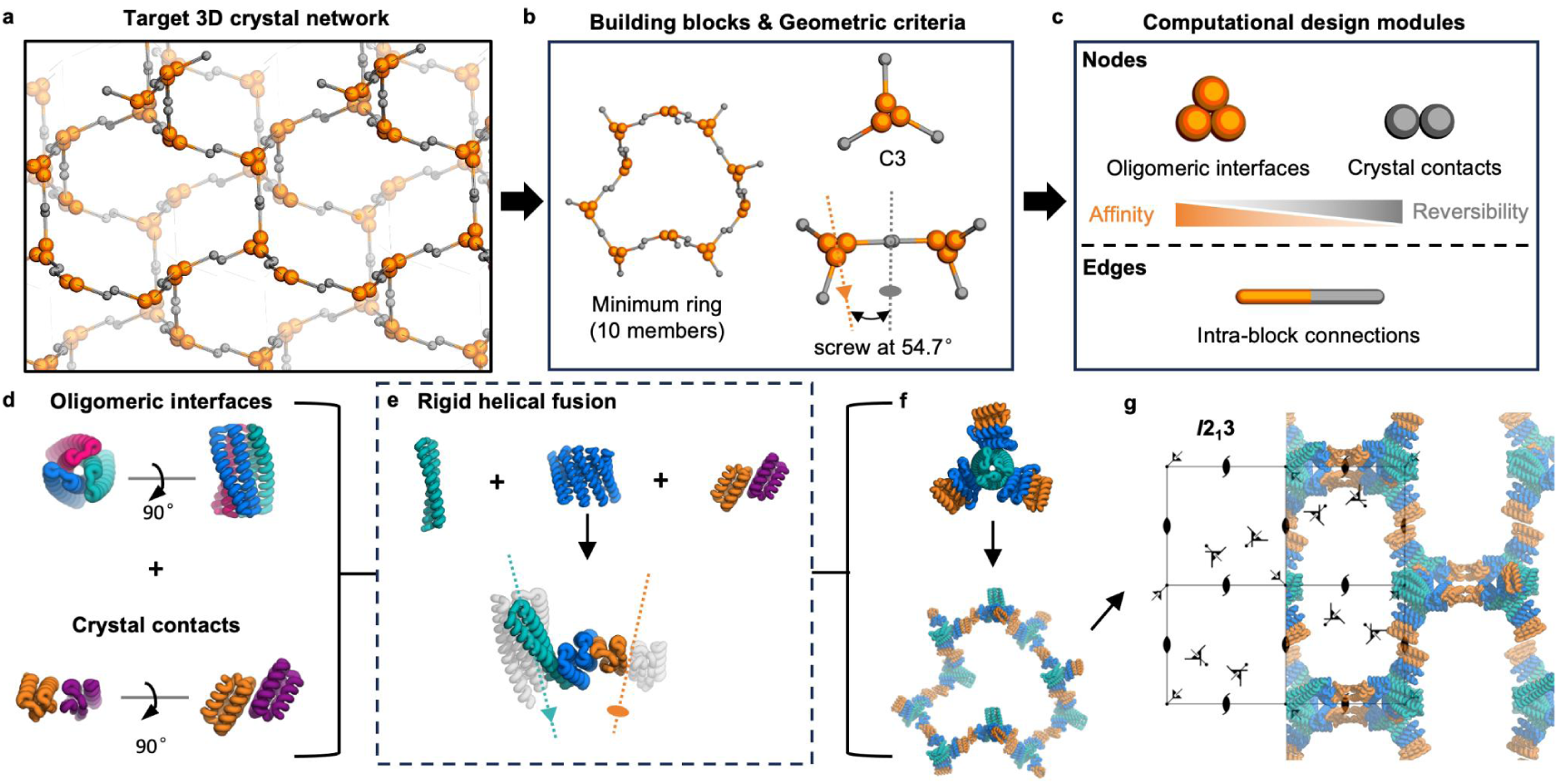
Modular computational design strategy for *I*2_1_3 space group. **a**, Schematic of the target three-dimensional crystal network, with spheres and rods as protein components, analogous to nodes and edges in a graph. **b**, Building blocks and geometric criteria for the network: the C3 cyclic axis of protein oligomers and the C2 cyclic axis of their interface skewed at 54.7°, forming a minimum 10-member ring assembly (left). **c**, Computationally designed modules building up the crystal lattice: nodes represent protein–protein interfaces, including high-affinity, low-reversibility oligomeric interfaces and low-affinity, high-reversibility crystal contacts, whereas edges correspond to covalent intra-block connections. **d**, Models of the oligomeric interface modules and the crystal contact modules. **e**, Architecture-guided rigid helical fusion for building block design. **f**, Models of designed protein building blocks and the minimum ring assembly indicated in panel b. **g**, Model of assembled 3D protein crystal superimposed with the symmetry elements of the *I*2_1_3 unit cell. In **d-g**, modules are color-coded; dash lines with triangles/ovals indicate the three-fold/two-fold cyclic axis of the corresponding modules.

**Extended Data Fig. 2.**
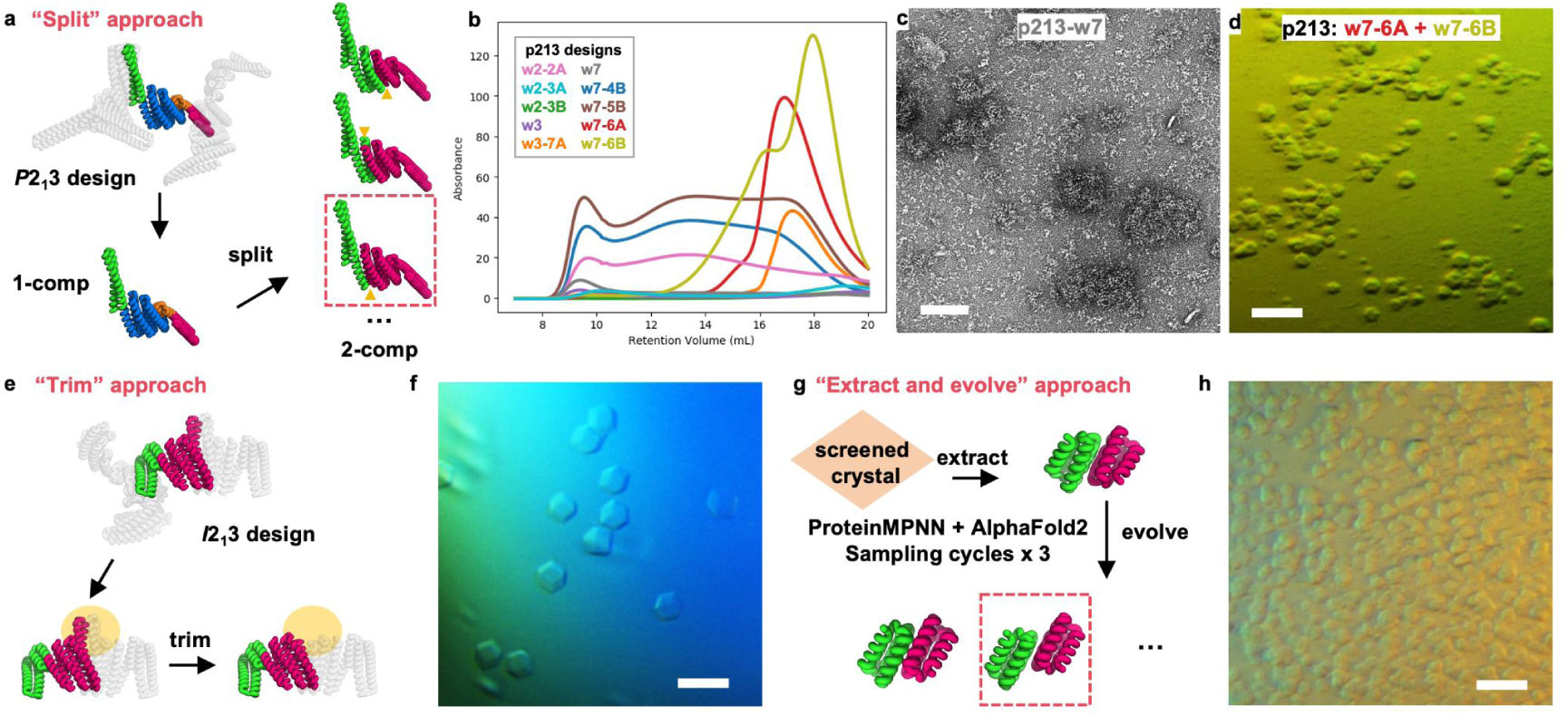
| Design strategies for weak, reversible crystal-contact modules and their experimental validation. **a**, Schematic of the “split” strategy, in which single-chain, one-component constructs were converted into two-component constructs by splitting the chain at different loop regions. **b**, SEC profiles of protein components before and after splitting. **c**, nsEM image of single-chain design p213-w7 after IMAC purification, showing that the protein formed disordered aggregates on its own. **d**, Optical microscopy image of crystals assembled from p213-w7-6A and p213-w7-6B, which were generated by splitting p213-w7. **e**, Schematic of the “trim” strategy, in which C2-symmetric protein interfaces were shortened and reconnected. **f**, Optical microscopy image of initially assembled crystals of design *I*2_1_3-0. **g**, Schematic of the “extract-and-evolve” strategy, in which iterative rounds of sequence design and structure prediction were used to evolve existing β-strand-edge contacts into modules with diverse interaction strengths. **h**, Optical microscopy image of initially assembled crystals of design *I*2_1_3-1. Scale bars: c, 100 nm; d, f and h, 100 µm.

**Extended Data Fig. 3.**
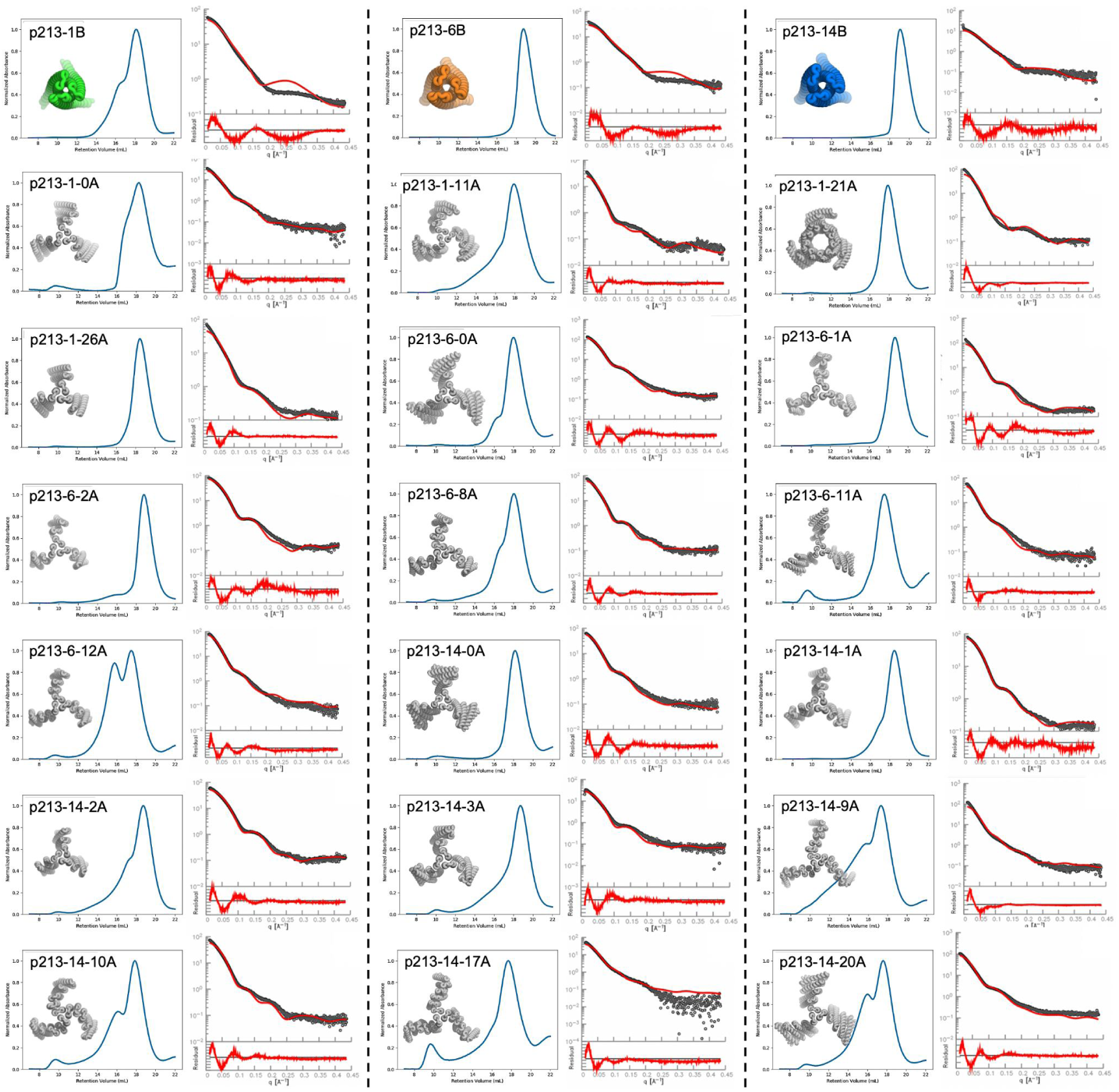
SEC and SAXS characterization of cyclic-symmetry protein components used for *P*2_1_3 crystal assembly. For each design, the computational model is shown alongside the SEC profile (Superose 6 Increase 10/300 column) and the experimental SAXS profile. SAXS data are shown as black circles and are compared with theoretical profiles calculated from the computational models using the Debye formula.

**Extended Data Fig. 4.**
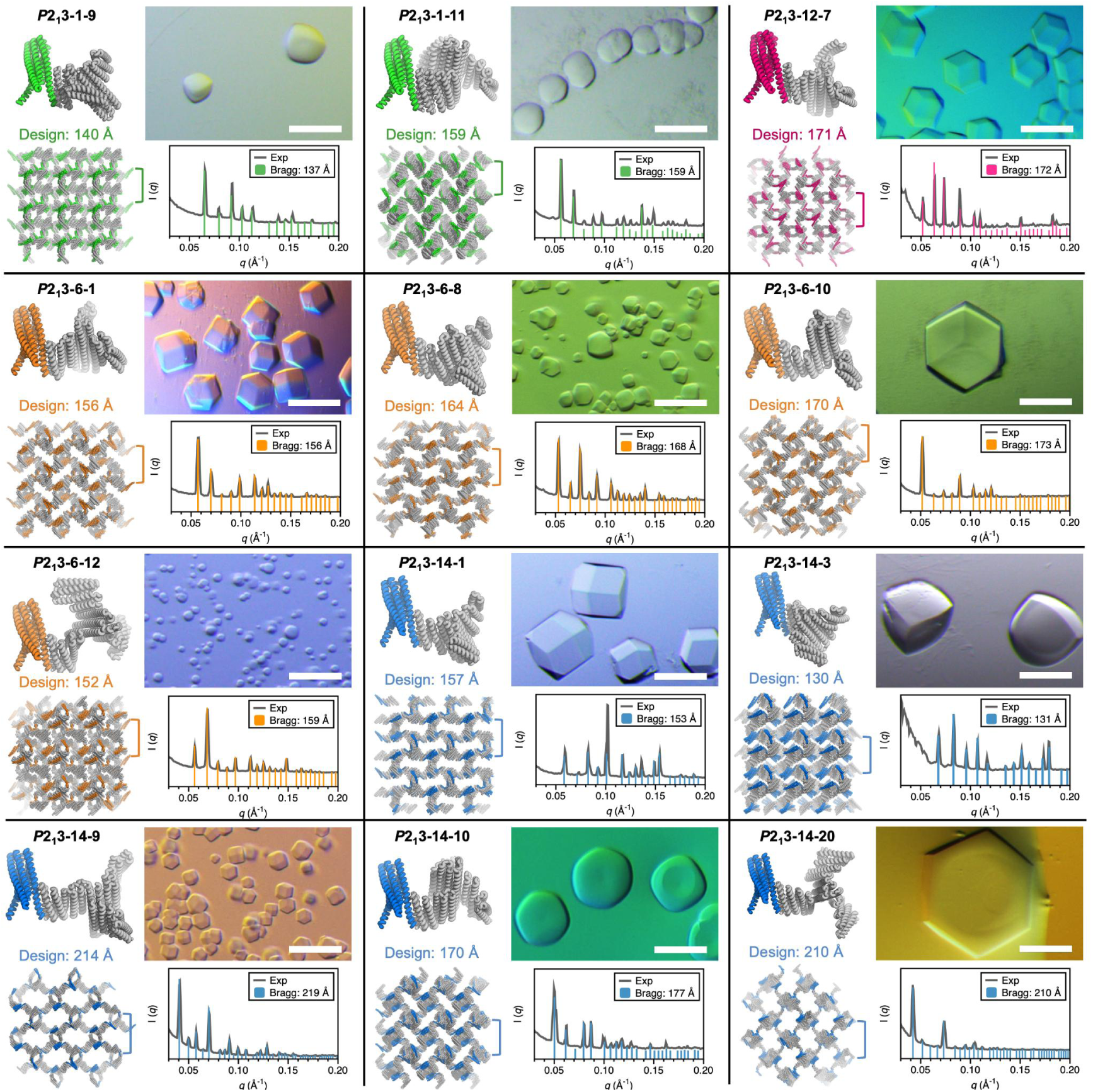
Computational design and experimental characterization of additional *P*2_1_3 crystals. Computational design models of the crystals alongside their experimental validation by optical microscopy and SAXS. For each crystal assembly: top left, two C3 components interacting through the designed crystal contacts; bottom left, symmetry-expanded protein crystal lattice viewed along the (001) direction with the designed unit cell parameter indicated above; top right, optical microscopy images of representative crystals. Scale bar, 100 µm; bottom right, experimental SAXS profile (Exp) fitted with theoretical Bragg diffraction peaks (Bragg), with corresponding experimental unit cell parameters shown in the legend.

**Extended Data Fig. 5.**
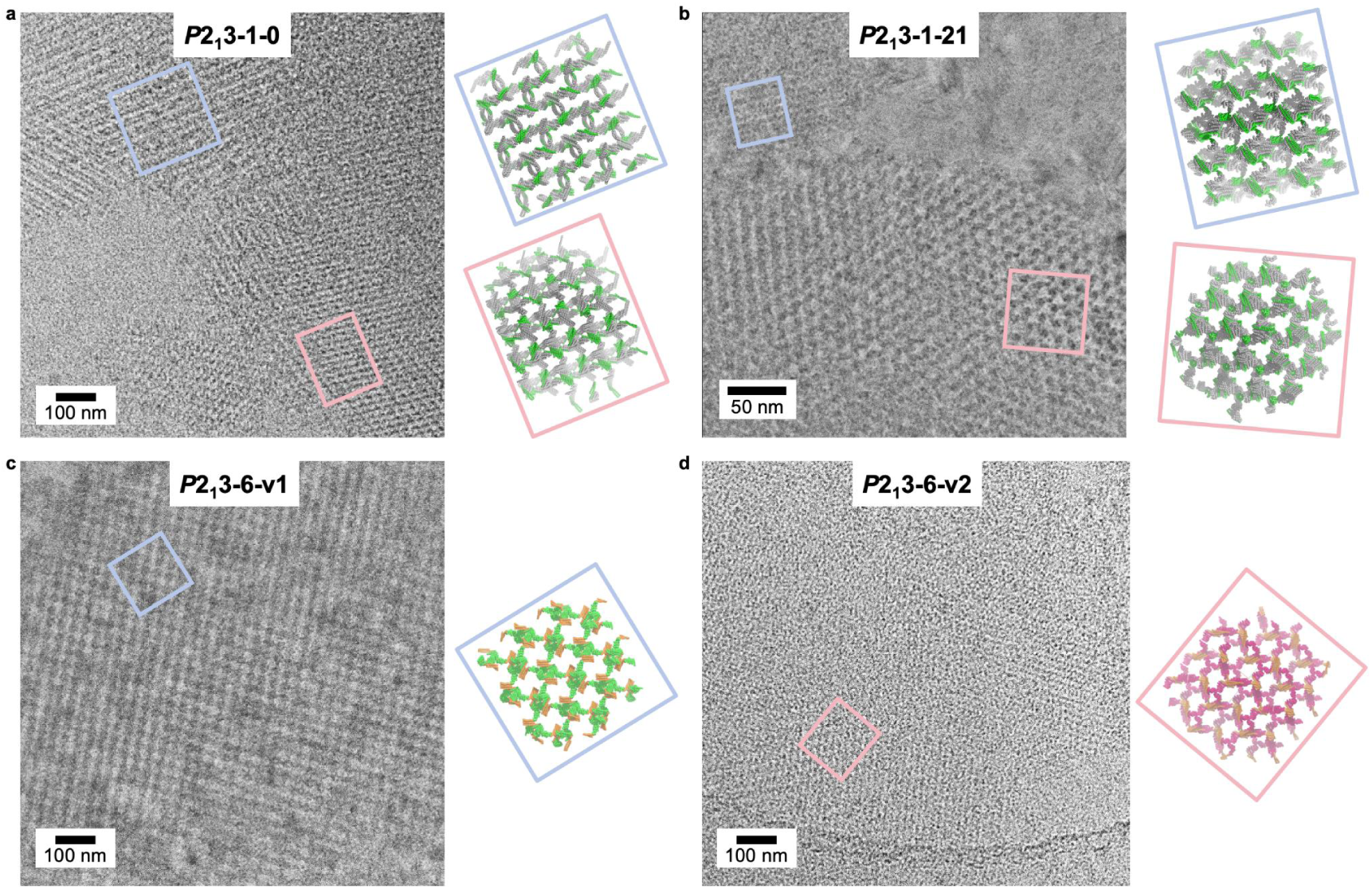
Representative cryoEM images of experimentally validated protein crystal designs. Lattice patches from the corresponding computational design models are shown in boxed insets alongside the micrographs.

**Extended Data Fig. 6.**
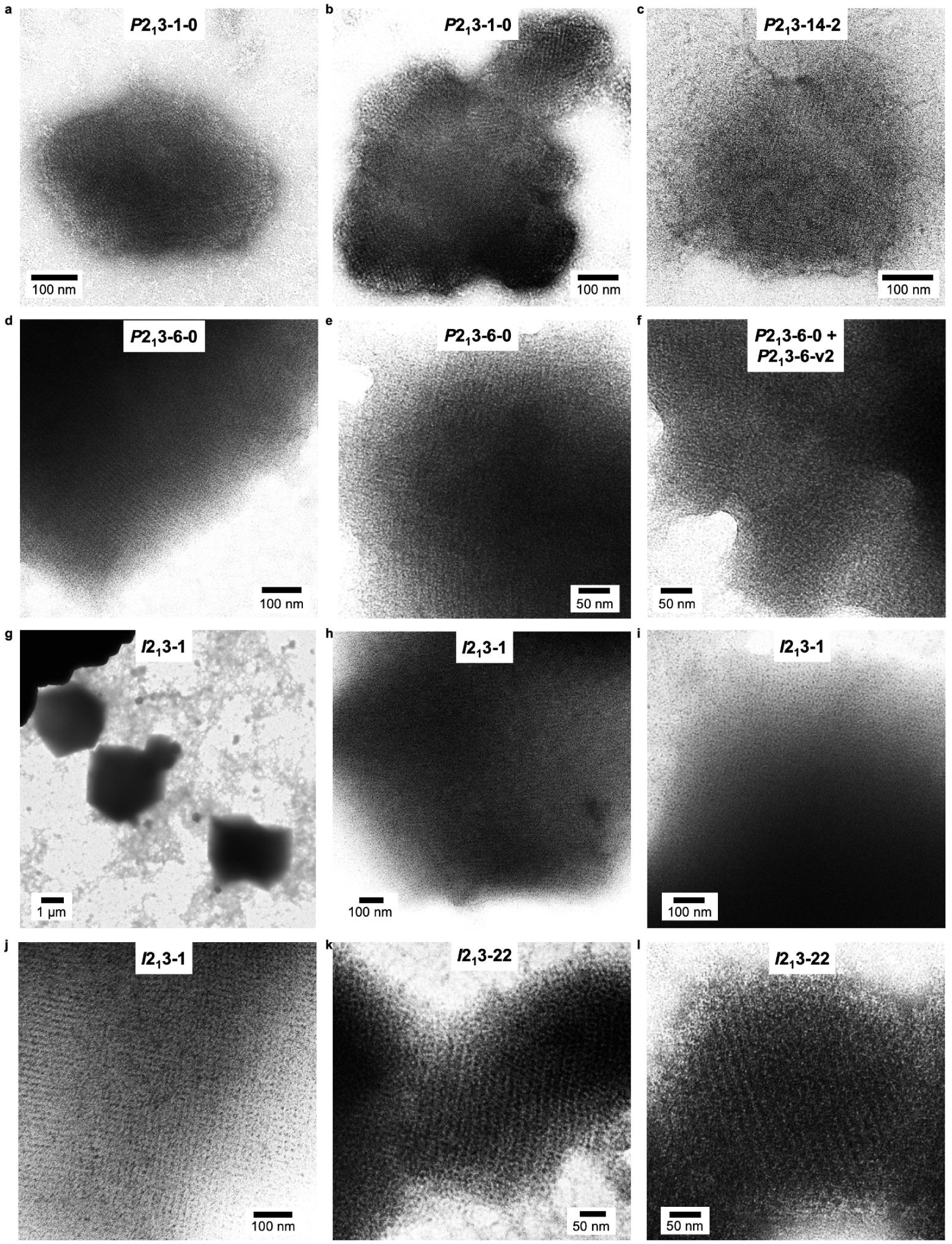
Representative nsEM images of validated protein crystal designs. **a-e**, *P*2_1_3 crystal lattices. **f**, *P*2_1_3 crystals of mixed, isomorphous lattices. **g-l**, *I*2_1_3 crystal lattices.

**Extended Data Fig. 7.**
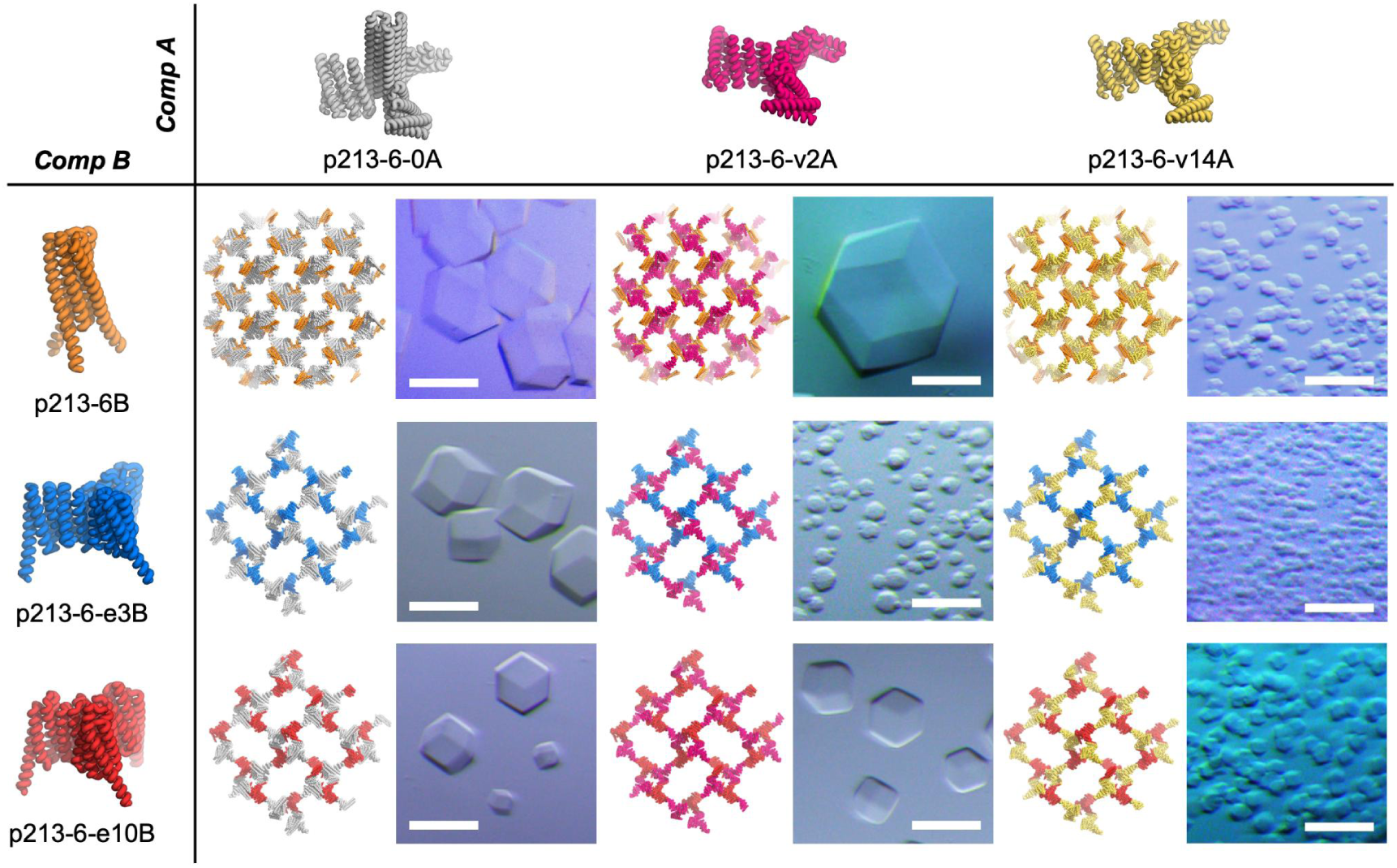
Modular self-assembly of protein crystals through recombination of individually engineered protein components. Three variants each of component A and component B, both C3-symmetric trimeric components of *P*2_1_3 crystals, were recombined in a row-by-column matrix to generate all nine pairwise combinations. Each combination resulted in macroscopic crystal assembly. Optical microscopy images are shown alongside the corresponding crystal-lattice design models viewed along the [100] direction. Scale bars, 100 µm.

**Extended Data Fig. 8.**
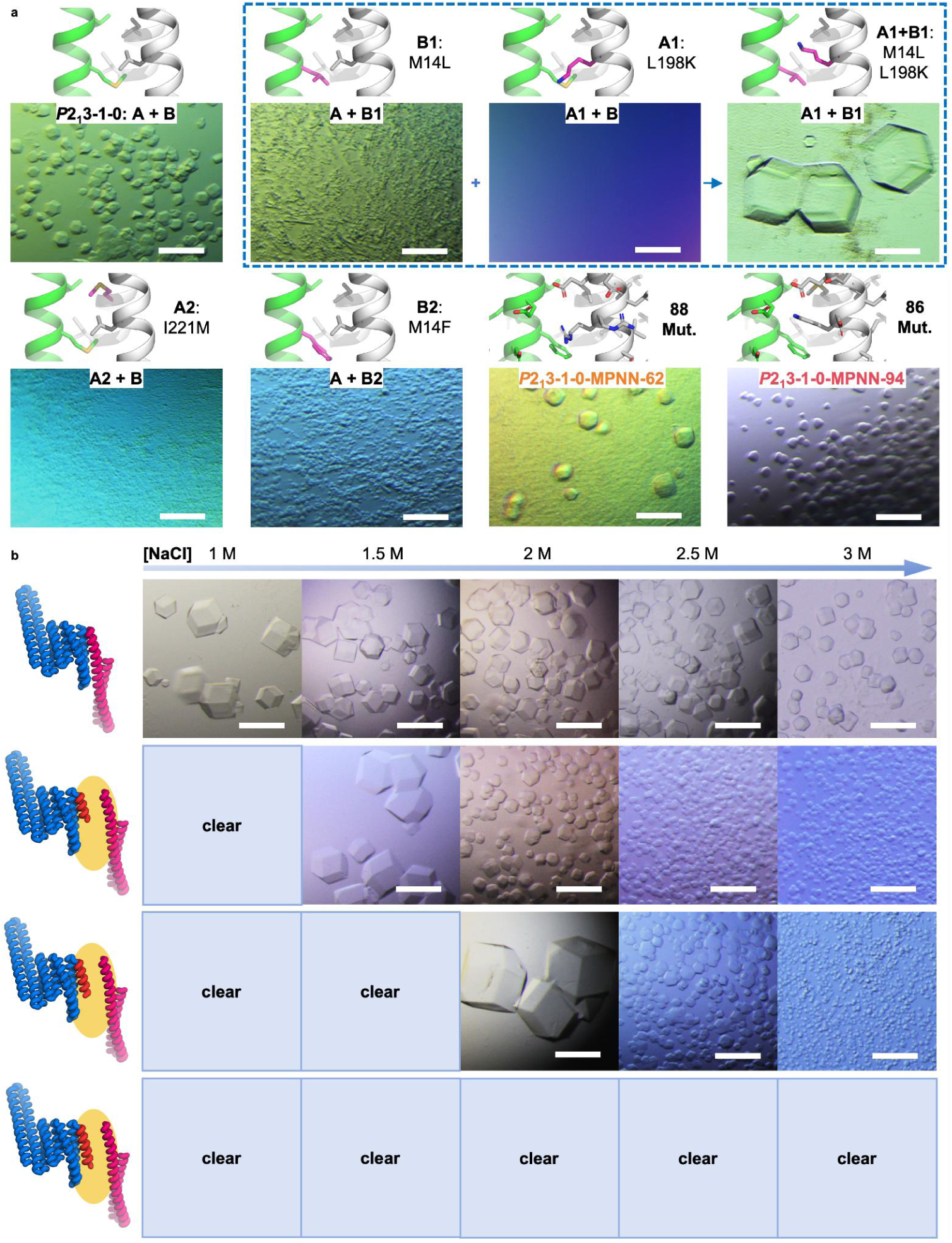
Modulation of protein crystal self-assembly by crystal-contact engineering. **a**, Optical microscopy images of crystals formed by interface point-mutant and ProteinMPNN-redesigned variants, shown alongside the corresponding design model and mutation labels. Point mutations are highlighted in pink in the model. Designs *P*2_1_3-1-0-MPNN-62 and *P*2_1_3-1-0-MPNN-94 contain extensive sequence redesigns of 88 and 86 positions, respectively. Mut., mutations. The dashed blue box highlights an example of compensatory interface mutations, in which mutation A1 eliminated self-assembly whereas mutation B1 enhanced self-assembly; combining the two produced intermediate assembly behavior with larger crystals. **b**, Modulation of crystal-assembly kinetics by shielding the crystal contact. Models are shown on the left, with shielding regions highlighted in yellow. Optical microscopy images on the right show crystal-assembly outcomes at elevated NaCl concentrations ranging from 1 to 3 M. Stronger shielding slowed crystal nucleation, requiring higher NaCl concentrations to initiate self-assembly and resulting in fewer crystals. Scale bars, 100 µm.

**Extended Data Fig. 9.**
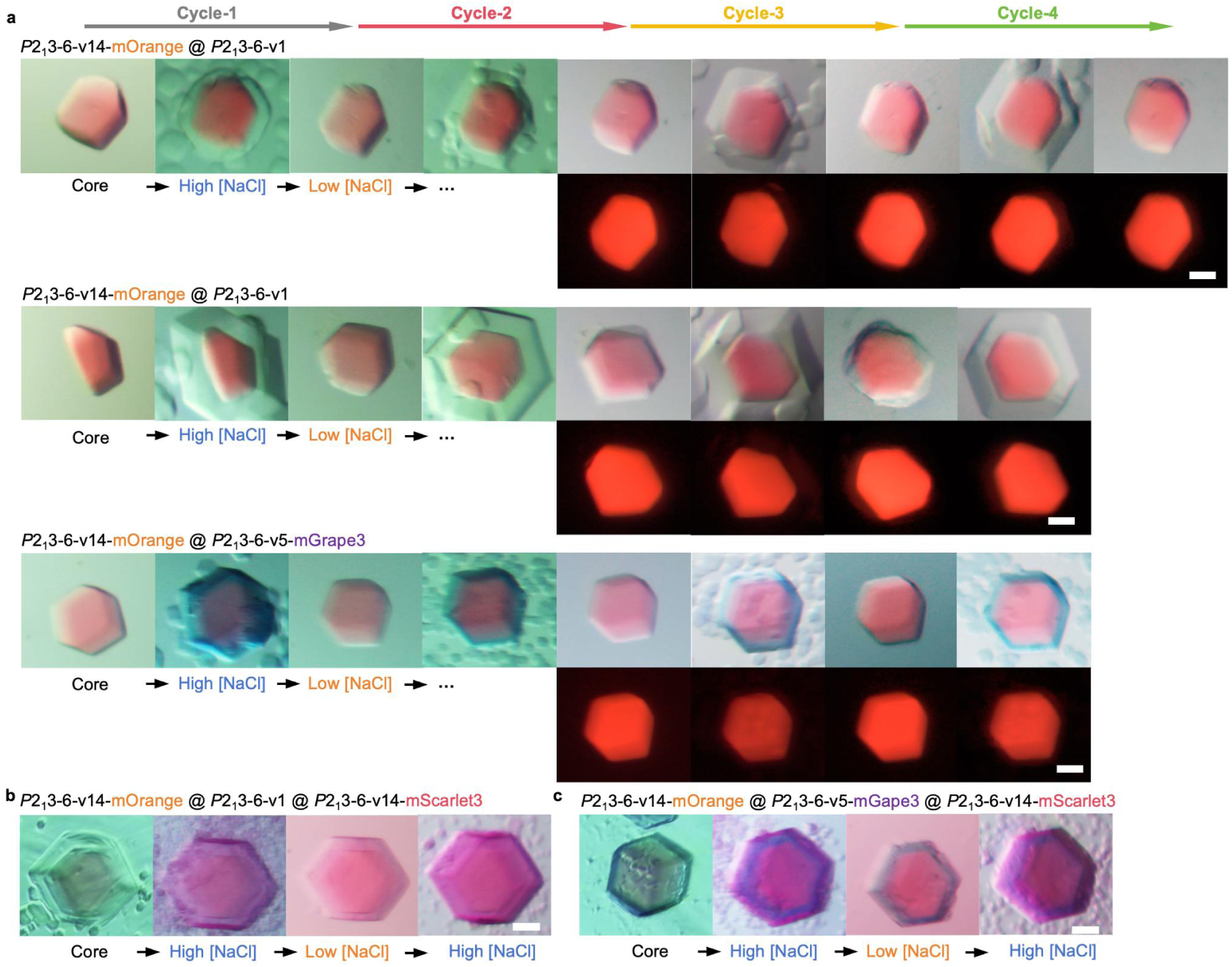
Reversible assembly and disassembly of core–shell crystals. **a**, Optical microscopy images showing reversible shell assembly and disassembly in two-layer core–shell crystals over four cycles. Two representative crystals are shown for *P*2_1_3-6-v14–mOrange@*P*2_1_3-6-v1, and one representative crystal is shown for *P*2_1_3-6-v14–mOrange@*P*2_1_3-6-v5–mGrape3. The *P*2_1_3-6-v1 shell assembled at 1.5 M NaCl and disassembled at 0.5 M NaCl, whereas the *P*2_1_3-6-v5–mGrape3 shell assembled at 1.5 M NaCl and disassembled at 0.75 M NaCl. Fluorescence images of mOrange protein fusion show that the core crystal remained intact throughout the cycles. **b**,**c**, Optical microscopy images showing reversible outer-shell assembly and disassembly in three-layer core–shell crystals: *P*2_1_3-6-v14–mOrange @ *P*2_1_3-6-v1 @ *P*2_1_3-6-v14–mScarlet3 (**b**) and *P*2_1_3-6-v14–mOrange @ *P*2_1_3-6-v5–mGrape3 @ *P*2_1_3-6-v14–mScarlet3 (**c**). In b and c, assembly and disassembly of the outermost *P*2_1_3-6-v14–mScarlet3 layer were performed at 2.5 M and 1.5 M NaCl, respectively. Scale bars, 50 µm.

**Extended Data Table 1.**
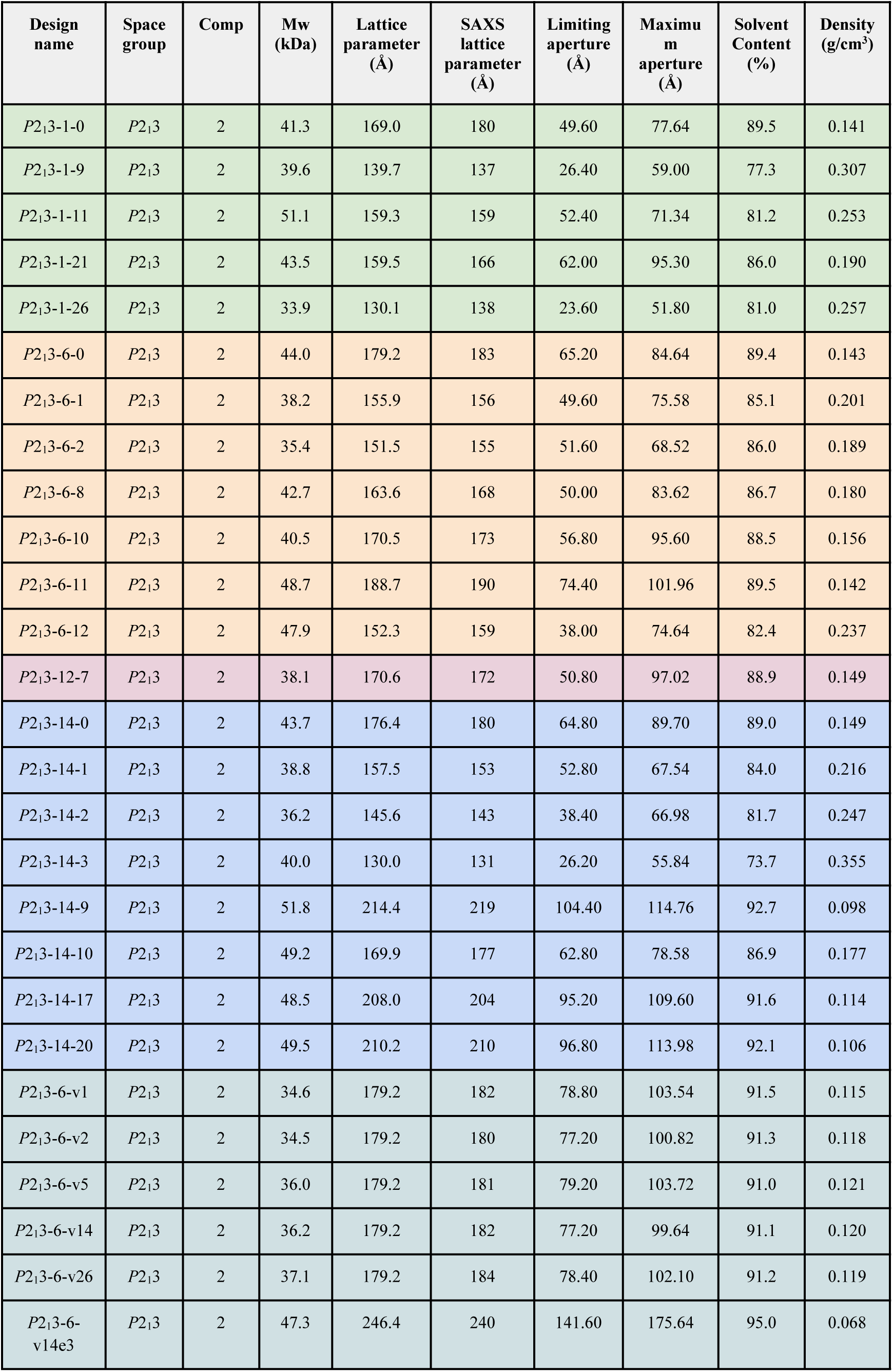
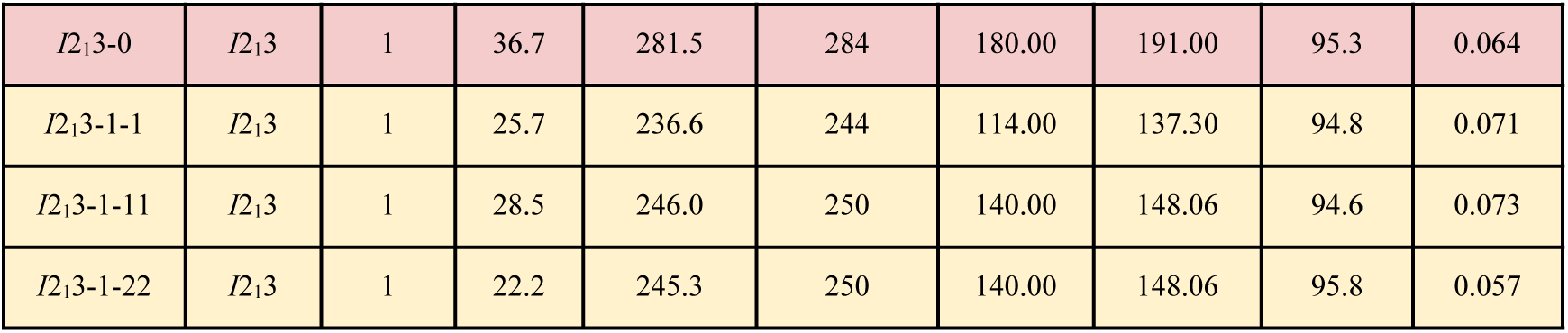
Structural and pore-architecture parameters of experimentally validated protein crystals.

## Notes

### Competing Interest Statement

The authors have declared no competing interest.

