## Supplementary_Information for "Programmable De Novo Design of Mesoporous Protein Crystal Frameworks"

**Supplementary Figures S1-S23**

**Supplementary Tables S1-S4**

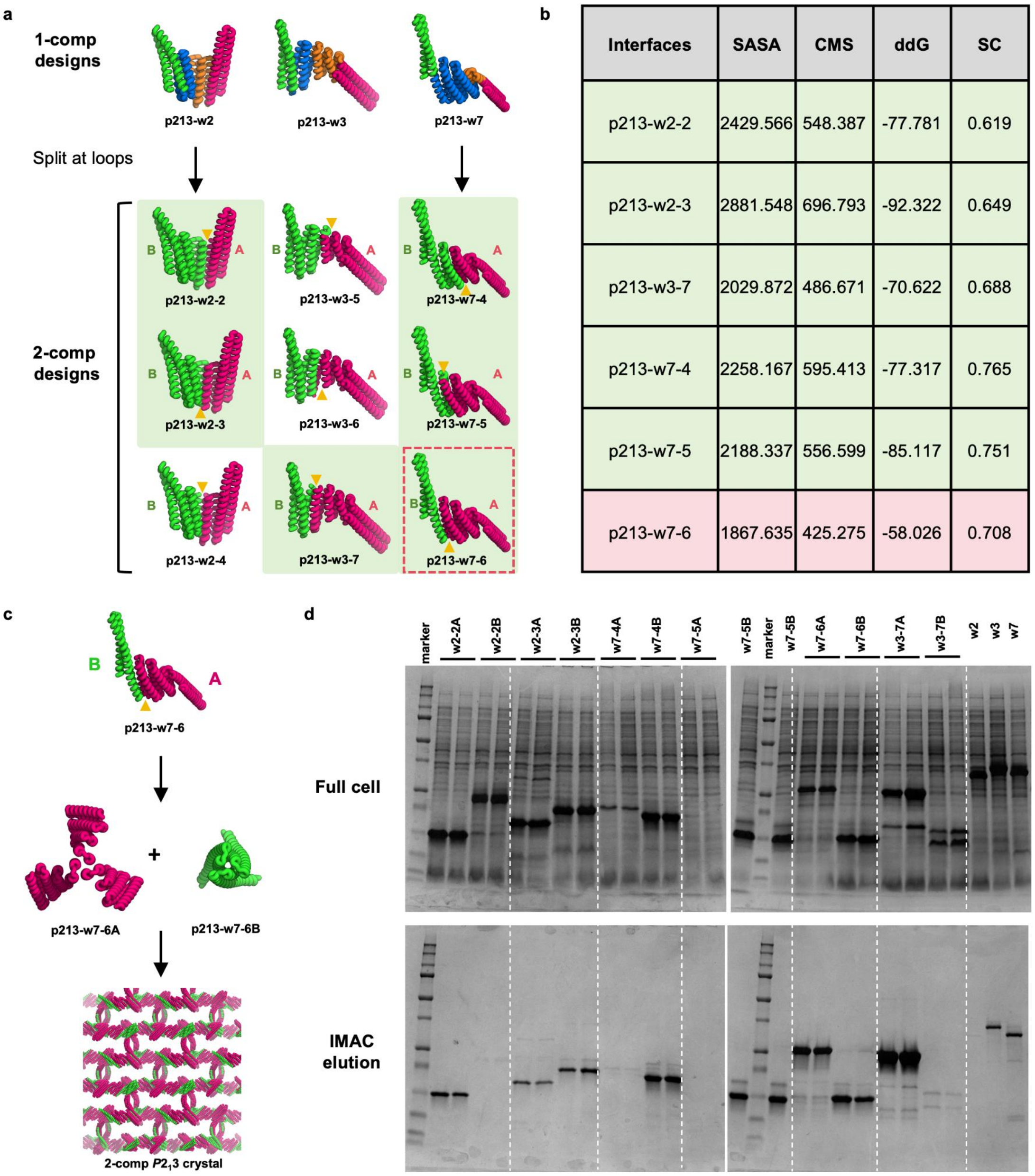


| Interfaces | SASA | CMS | ddG | SC |
|---|---|---|---|---|
| p213-w2-2 | 2429.566 | 548.387 | -77.781 | 0.619 |
| p213-w2-3 | 2881.548 | 696.793 | -92.322 | 0.649 |
| p213-w3-7 | 2029.872 | 486.671 | -70.622 | 0.688 |
| p213-w7-4 | 2258.167 | 595.413 | -77.317 | 0.765 |
| p213-w7-5 | 2188.337 | 556.599 | -85.117 | 0.751 |
| p213-w7-6 | 1867.635 | 425.275 | -58.026 | 0.708 |

**Supplementary Fig. S1 Design and experimental screening for the "split" crystal contacts of $P2_13$ protein crystals. a**, Three one-component $P2_13$ crystal designs were strategically split at distinct locations to generate two-component assemblies, introducing a newly formed protein–protein interface between them. Designs with green background color were selected for experimental characterization. Design in dashed red box was the working design for successful crystal self-assembly. **b**, Rosetta metric calculation results of the experimental tested interfaces by "split". **c,** Scheme of the self-assembly process showing two protein trimers generated by splitting the original design, associating into the target $P2_13$ crystal lattice. **c,** SDS-PAGE analysis for the protein expression and purification by IMAC. Two parallel samples were tested for each split design. White dashed lines added for visual guidance of band alignment.

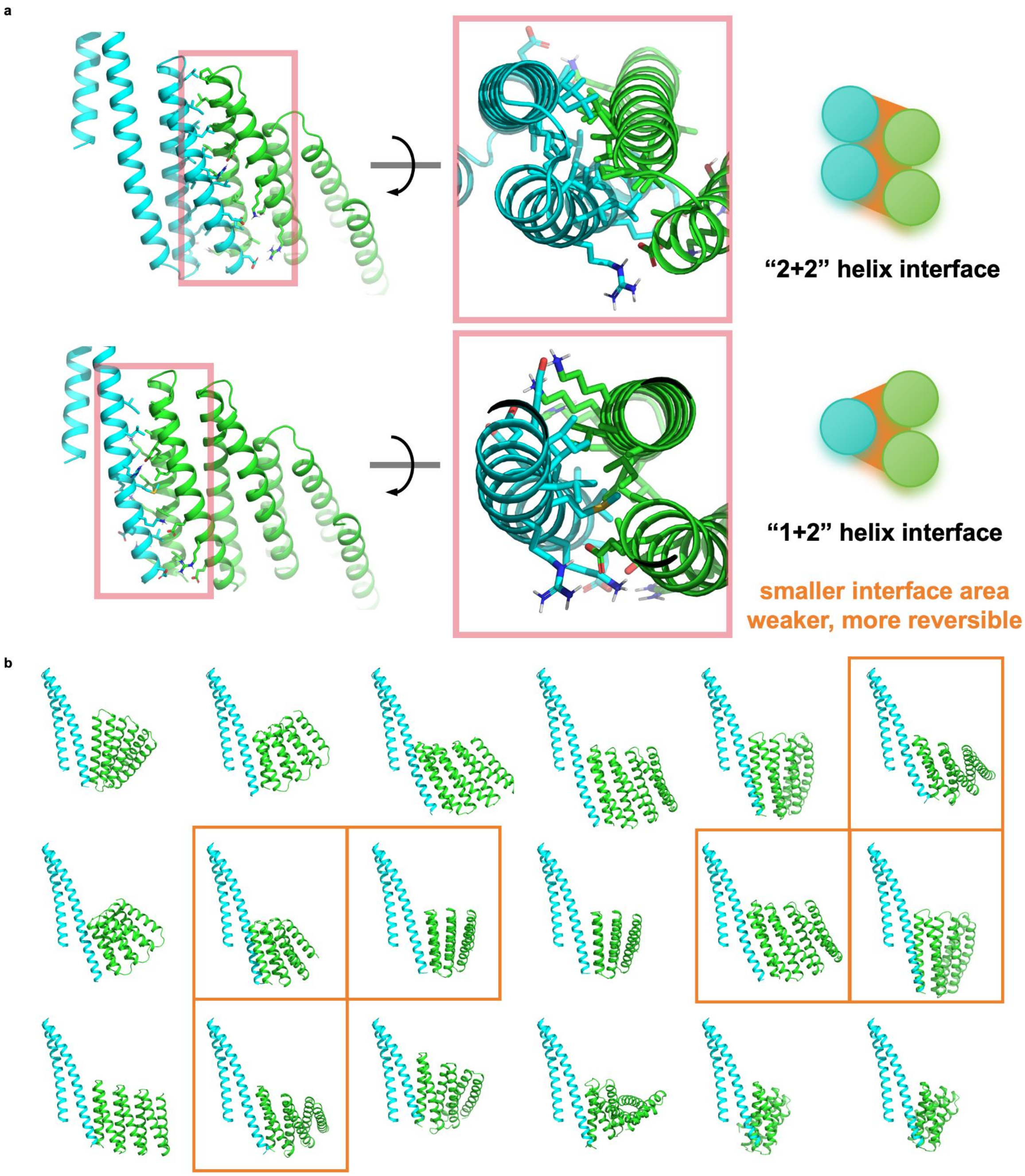


**Supplementary Fig. S2 Identification of “1+2” helix interfaces as functional crystal contacts and design of constructs with similar features. a**, Comparison between a more conventional “2+2” helix interface and the experimentally validated “1+2” helix interface observed in crystal assembly. The “1+2” interface has a smaller buried surface area, which may contribute to a weaker and more reversible protein–protein interaction. **b**, Design models of “1+2” helix-interface modules generated by applying the “split” strategy to a library of helical bundles fused with de novo helical repeat proteins. Modules with soluble, monodispersed bundle components (in cyan) are labeled in orange boxes.

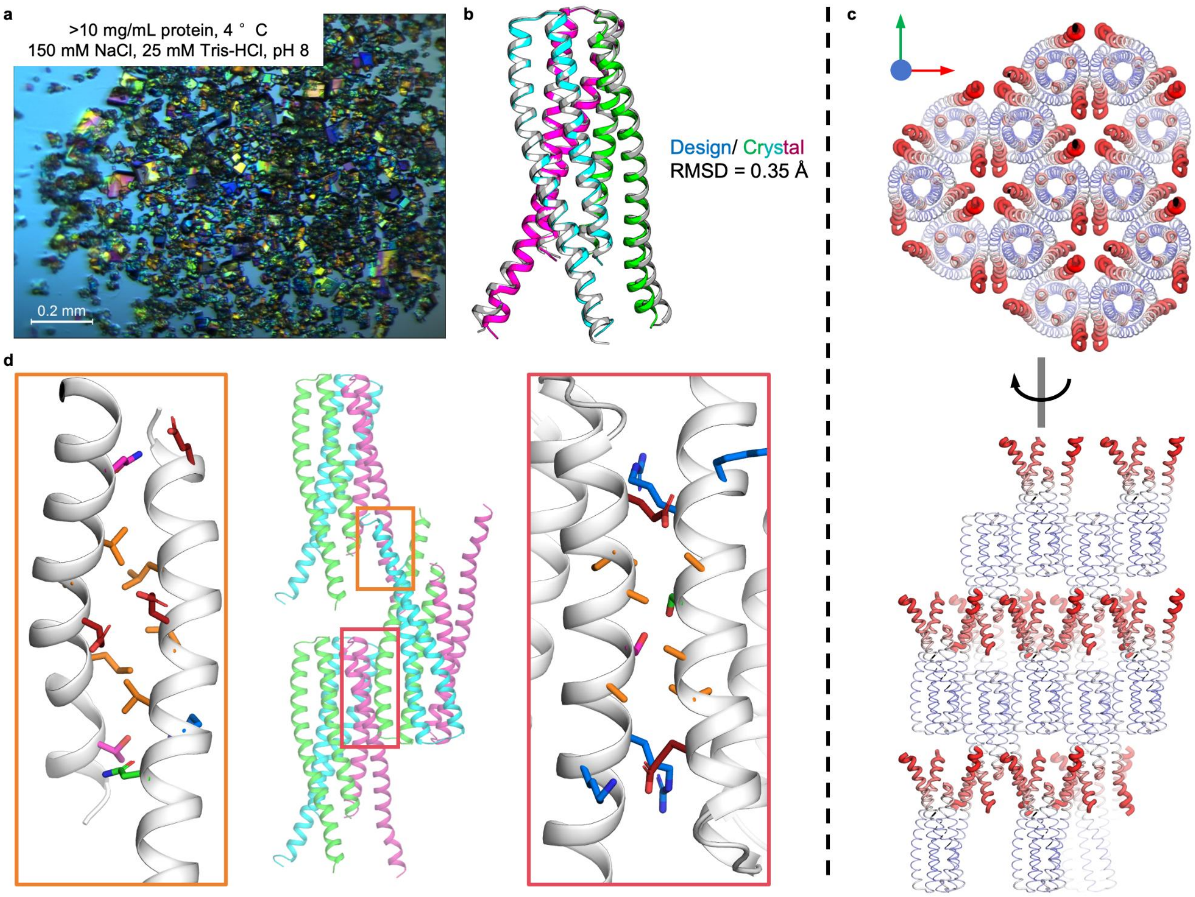


**Supplementary Fig. S3 X-ray crystallographic structure determination of component p213-14B. a**, Accidental crystals obtained from a concentrated stock solution of purified p213-14B protein. **b**, Alignment of the experimentally determined crystal structure with the computational design model, showing an all-atom root-mean-square deviation (RMSD) of 0.35 Å. **c**, Crystal packing of the trimeric components in the solved structure, viewed along the z and x axes. Regional B factors are shown in sausage representation, indicating higher flexibility in the single-helix region. **d**, Crystal contacts observed in the accidental p213-14B crystals. Two distinct contacts, located at the bundle and single-helix regions, both feature small hydrophobic patches.

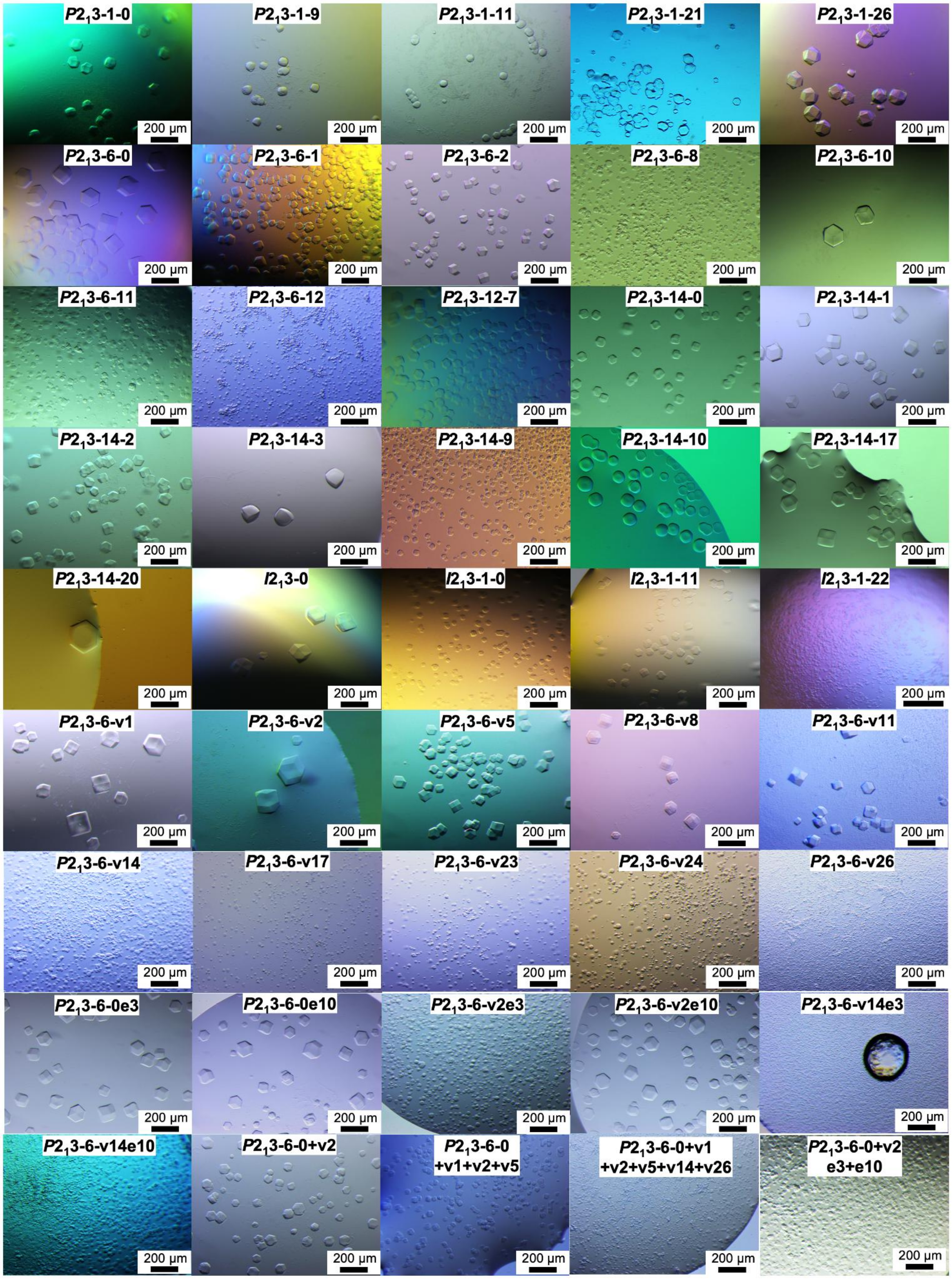


**Supplementary Fig. S4 Large-field optical microscopy images of assembled P213 and I213 crystals.**

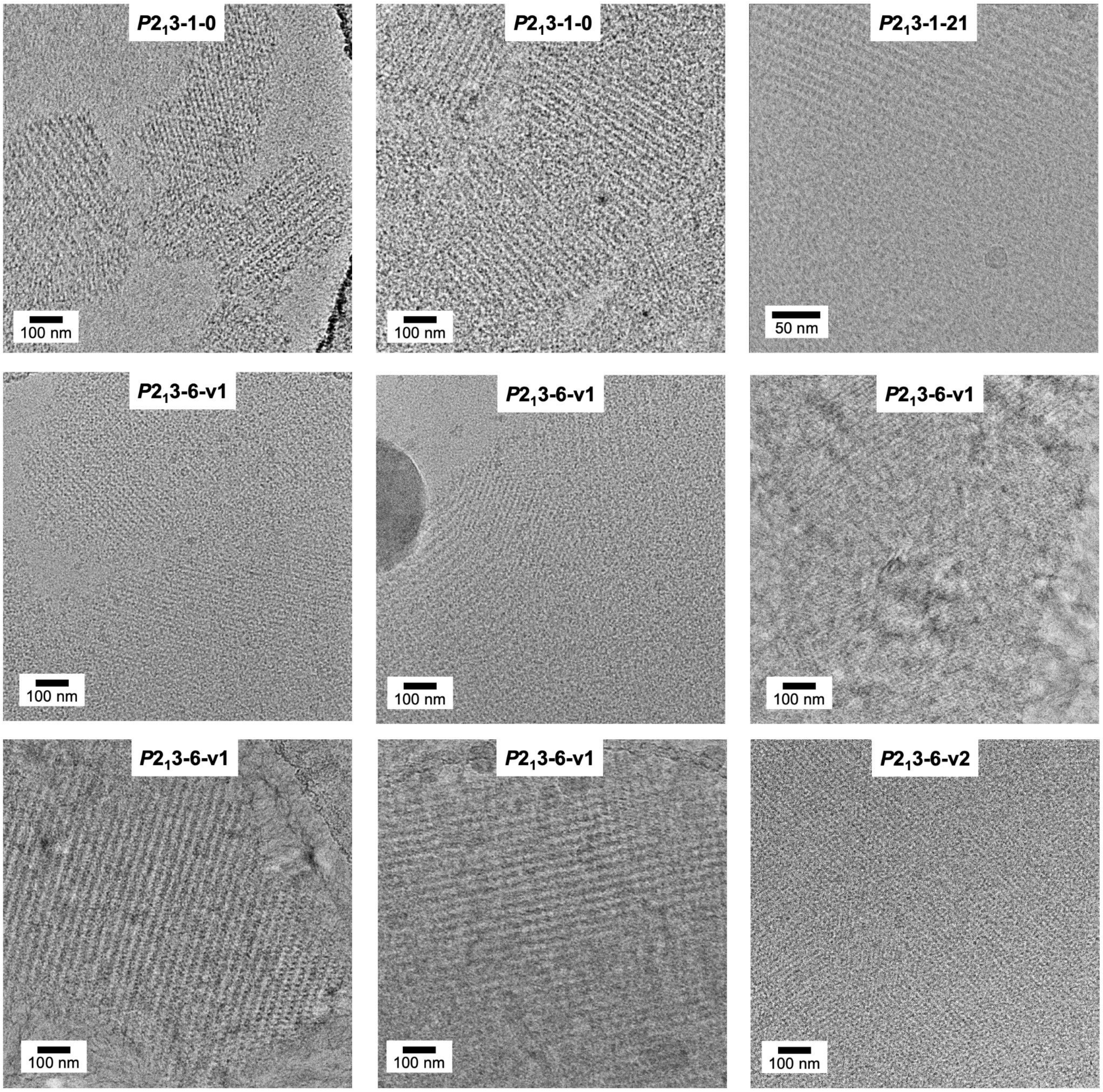


**Supplementary Fig. S5 Additional representative cryoEM images of experimentally validated protein crystal designs.**

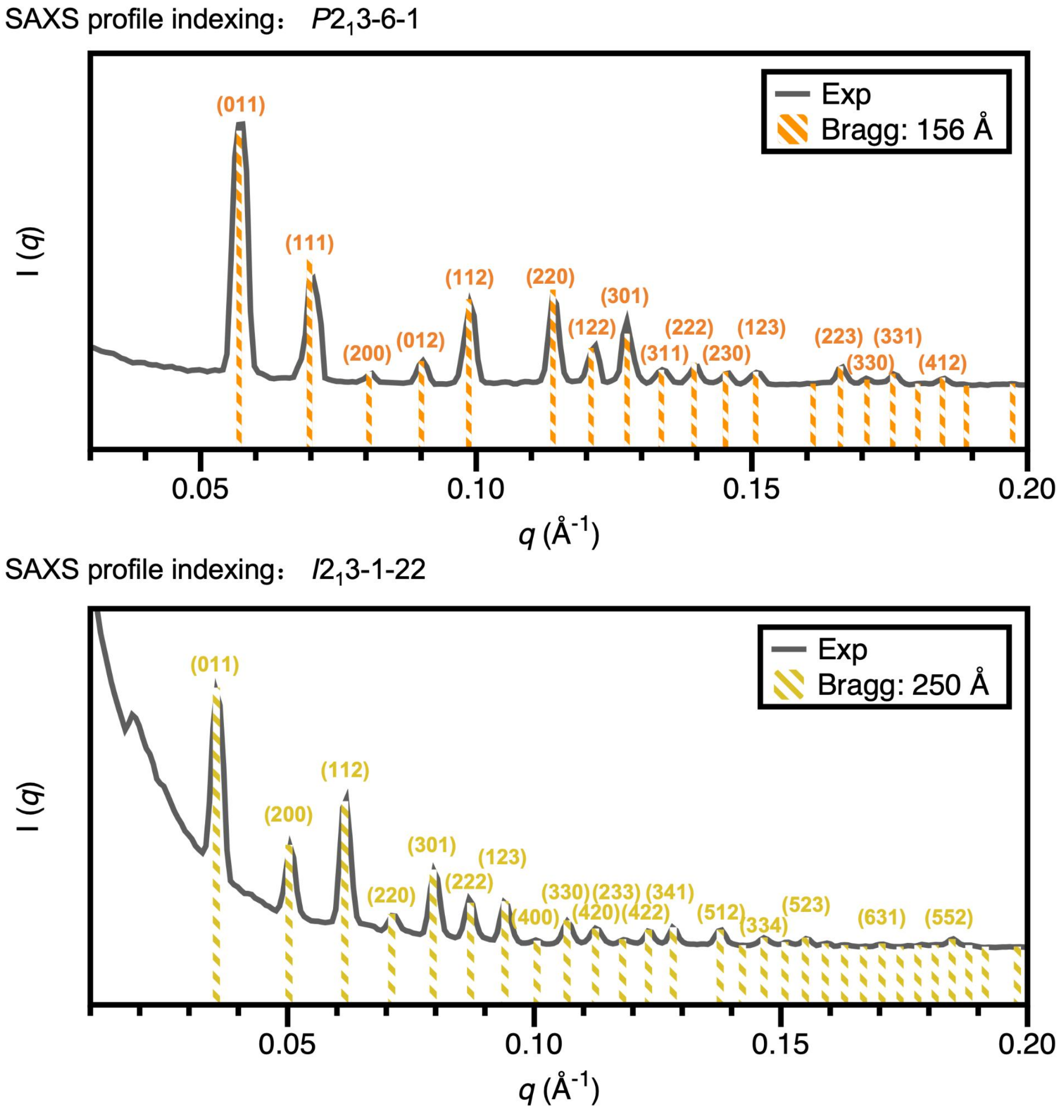


**Supplementary Fig. S6 Representative SAXS indexing examples showing the assignment of observed scattering peaks to their corresponding crystallographic indices.** Experimental SAXS profile (Exp) fitted with theoretical Bragg diffraction peaks (Bragg), with corresponding experimental unit cell parameters shown in the legend.

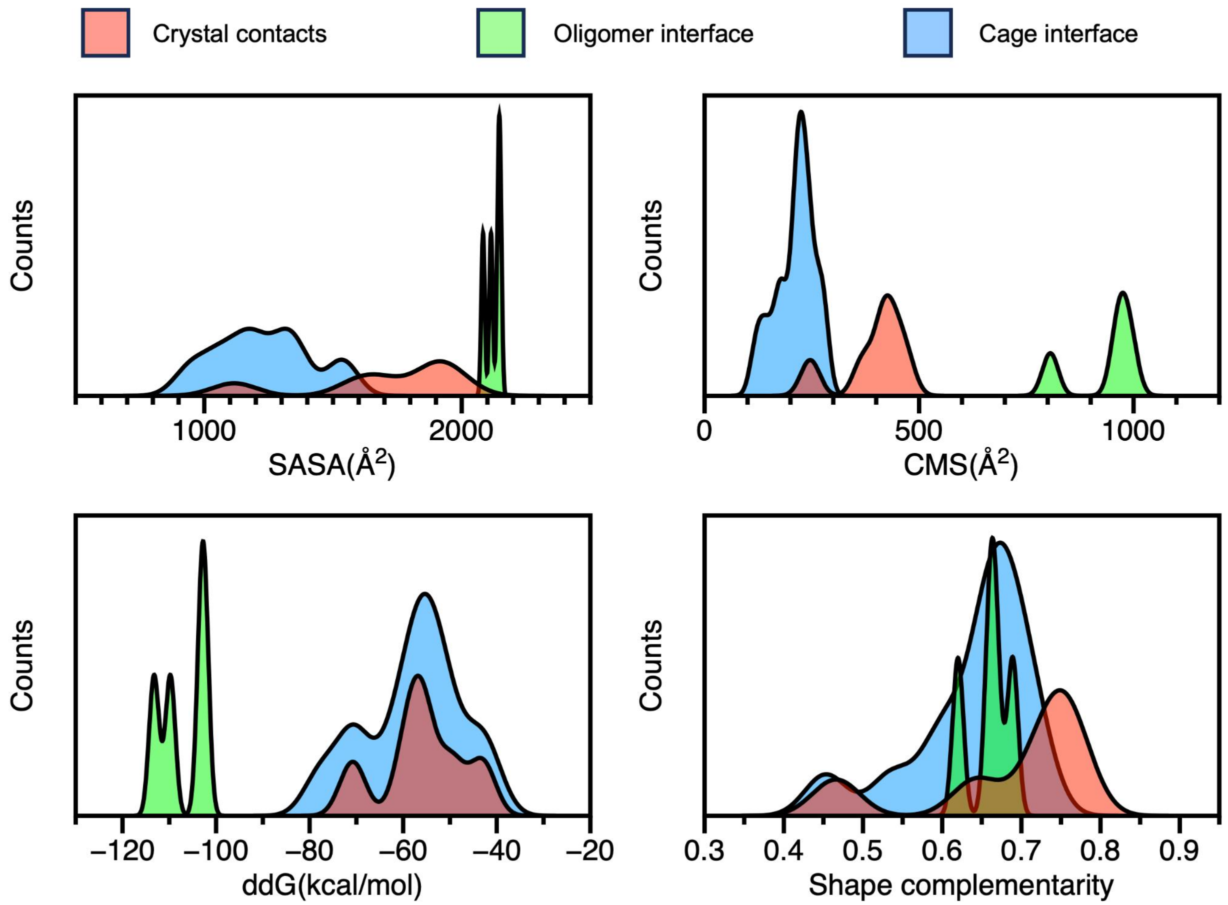


**Supplementary Fig. S7 Distributions of Rosetta metrics for validated crystal-contact modules and oligomeric modules, compared with previously designed de novo nanocage interfaces.** The validated crystal-contact modules exhibited interface areas intermediate between those of oligomeric modules and nanocage interfaces, while maintaining interface energies comparable to those of nanocage interfaces and slightly higher shape-complementarity scores.

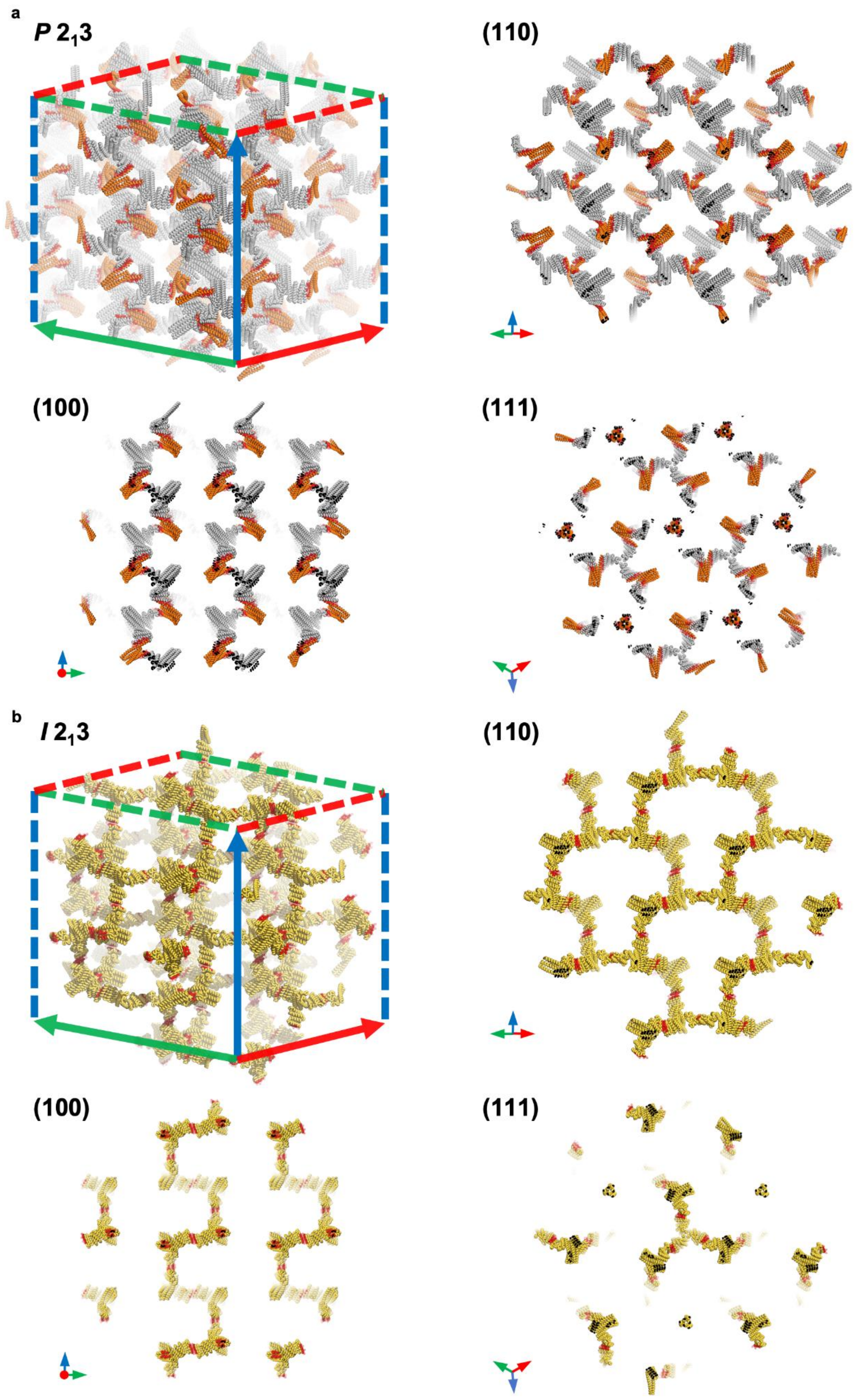


**Supplementary Fig. S8 Facet models of designed protein crystals. a**, Three-dimensional lattice model of the designed $P2_13$-6-0 crystal, showing representative (110), (100) and (111)

facet slices. **b**, Three-dimensional lattice model of the designed $I2_13$-1-0 crystal, showing representative (110), (100) and (111) facet slices. Crystal contacts are highlighted in red. In both lattices, the (110) slices contain the most densely packed protein-protein interaction networks, including interconnected 10-membered minimum assembly rings. Under a broken-bond approximation and Wulff construction, these more densely connected planes are expected to have lower surface energy and to be preferentially exposed, consistent with the observed rhombic dodecahedral crystal morphologies.

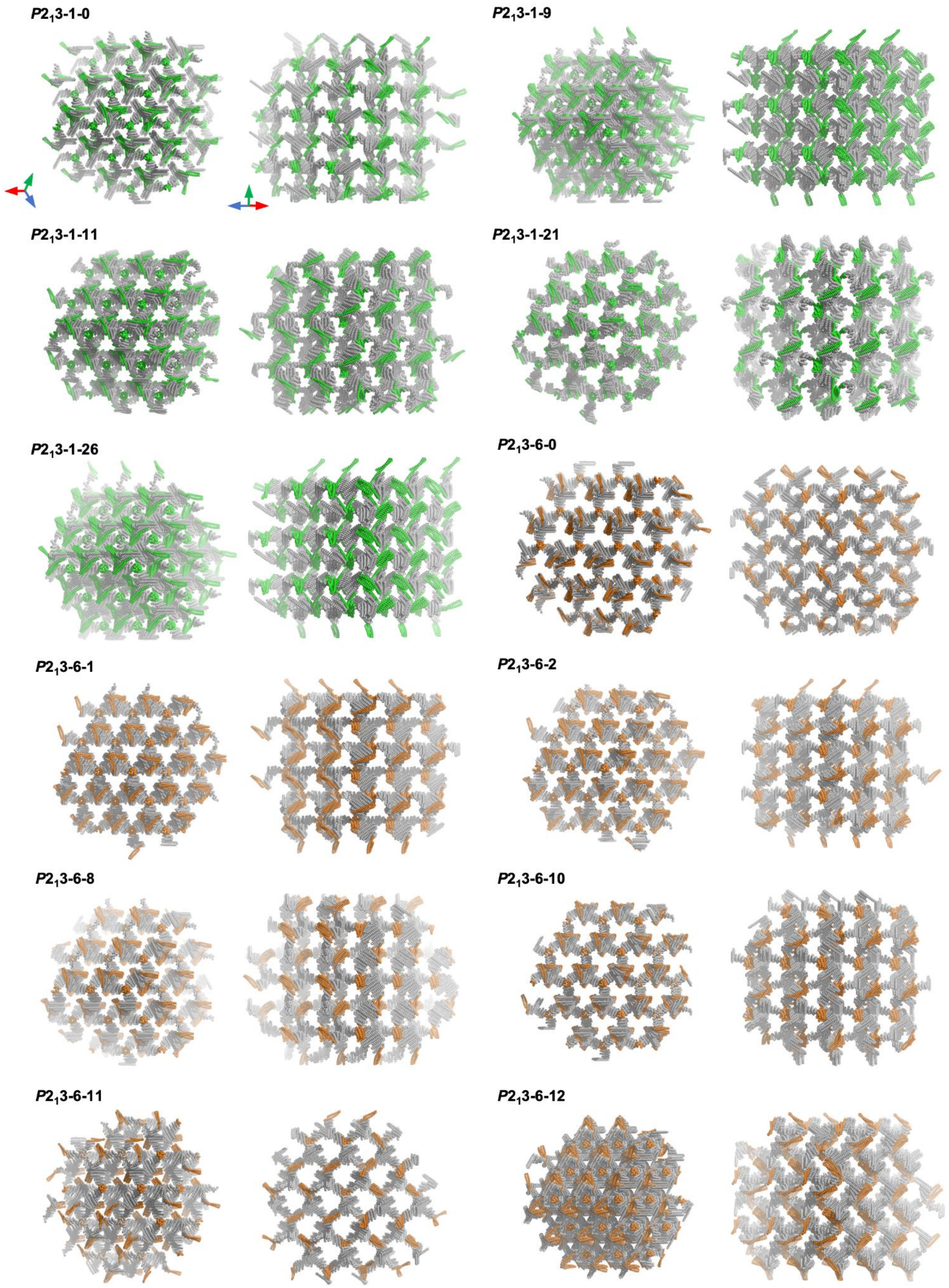


**Supplementary Fig. S9 Design models of validated protein crystal lattices viewed along the [111] and [110] directions.**

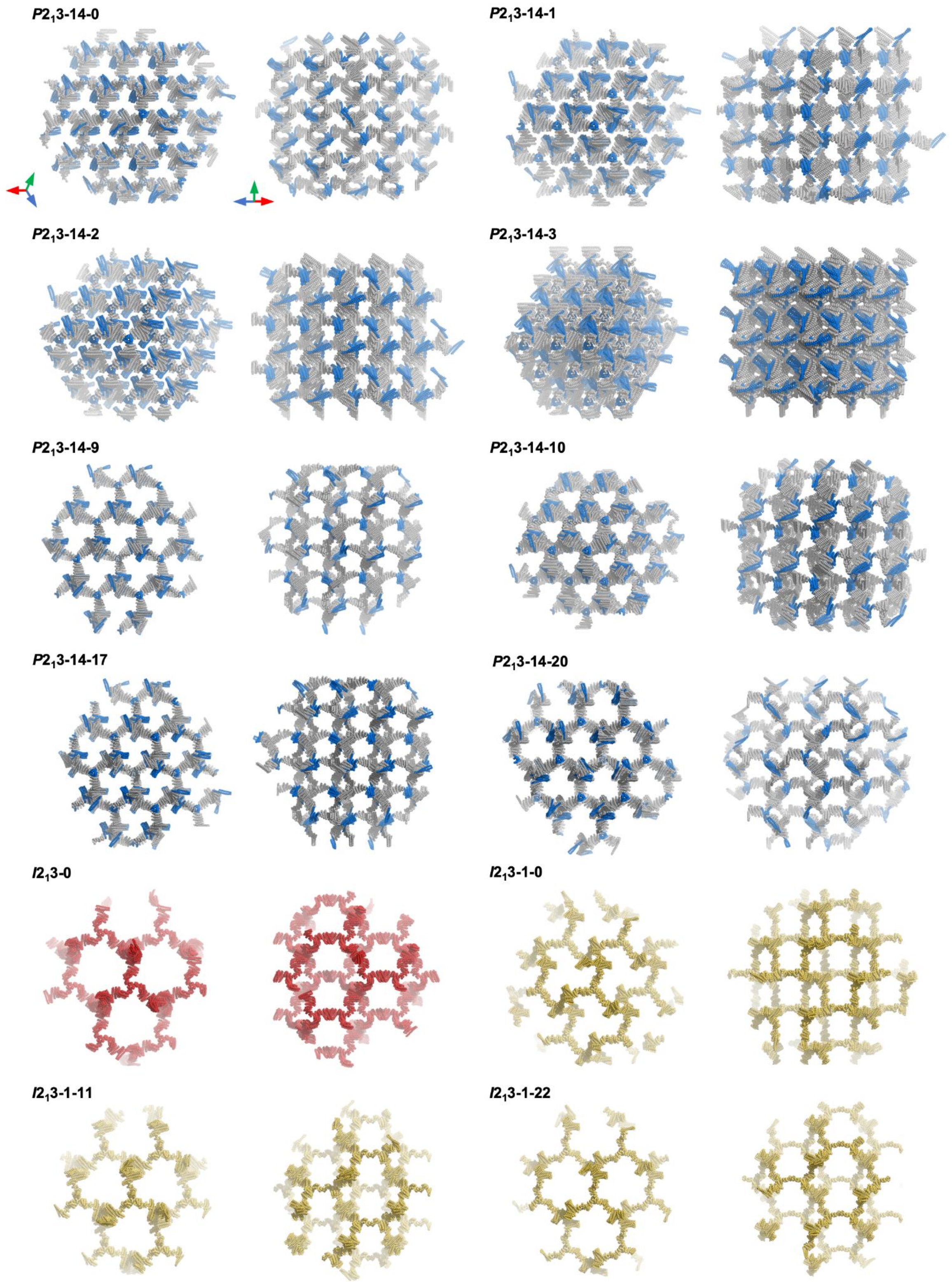


**Supplementary Fig. S10 Design models of additional validated protein crystal lattices viewed along the [111] and [110] directions.**

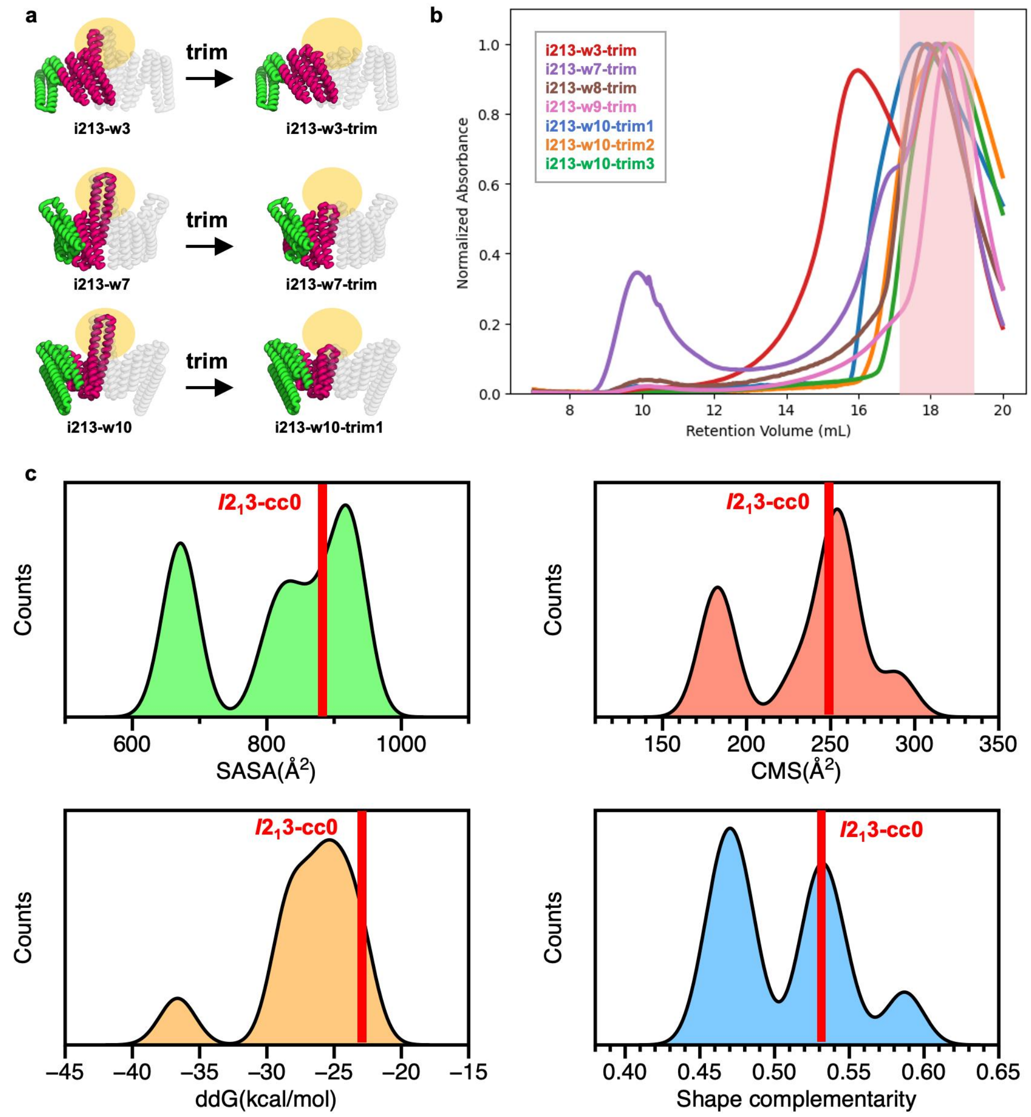


**Supplementary Fig. S11 Design and experimental screening of "trimmed" crystal contacts of $I2_13$ protein crystals.** **a**, Schematic of the "trim" strategy applied to different C2-symmetric protein interfaces. **b**, SEC profiles of crystal-assembly components designed with different trimmed crystal contacts. **c**, Rosetta metrics of the validated crystal contact $I2_13$-cc0 among all experimentally tested "trimmed" crystal contact modules. The validated contact has a relatively less negative predicted interface ddG.

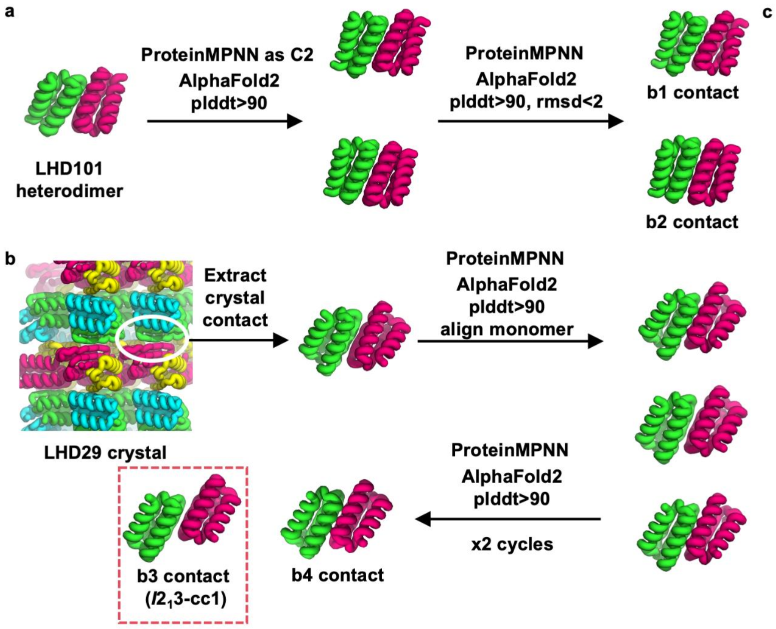


c

| Interfaces | SASA | CMS | ddG | SC |
|---|---|---|---|---|
| LC2-1-52 | 703.545 | 302.261 | -40.777 | 0.663 |
| LC2-1-57 | 772.746 | 322.624 | -37.244 | 0.646 |
| LC2-2-1 | 621.18 | 257.022 | -25.958 | 0.538 |
| LC2-2-31 | 661.465 | 258.306 | -35.61 | 0.648 |
| LC2-2-36 | 664.976 | 258.81 | -32.465 | 0.549 |
| LC2-2-7 | 642.58 | 262.769 | -28.171 | 0.549 |
| LC2-b4 | 642.203 | 236.329 | -21.738 | 0.549 |
| LC2-b3 ($I2_13$-cc1) | 454.346 | 123.081 | -7.044 | 0.487 |

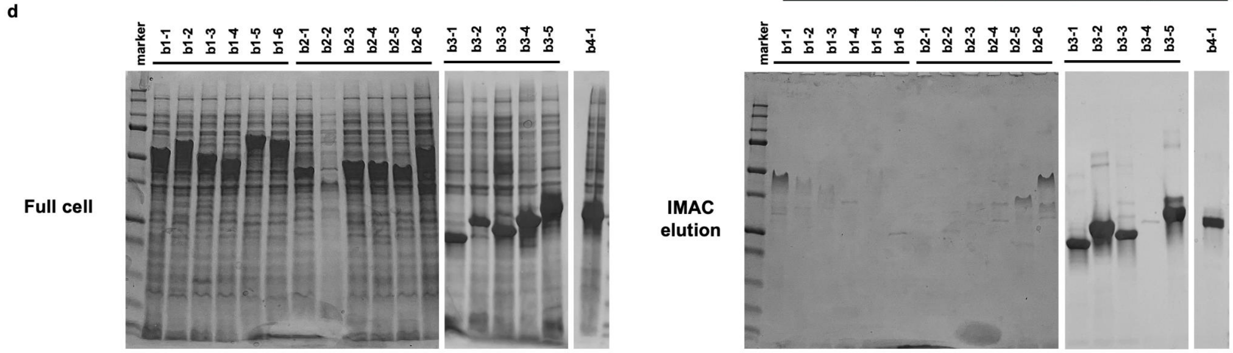


**Supplementary Fig. S12 Design and experimental screening of the "extract and evolve" crystal contacts of $\boldsymbol{I2_13}$ protein crystals. a**, **b**, Schematic of the "extract and evolve" strategy applied to (a) LHD101 heterodimer[1] and (b)LHD29 crystal contact (PDB ID: 6WMK). **c**, Rosetta metric calculation results of the experimental tested interfaces by "extract and evolve". **d**, SDS-PAGE analysis of the designed crystal-assembly components, for both full-cell protein expression and purification by IMAC. For the 12 designs featuring b1 and b2 contacts, expression was observed in the full-cell fraction, but no soluble protein was recovered following IMAC. This lack of solubility is likely due to inclusion body formation driven by strong inter-component interactions.

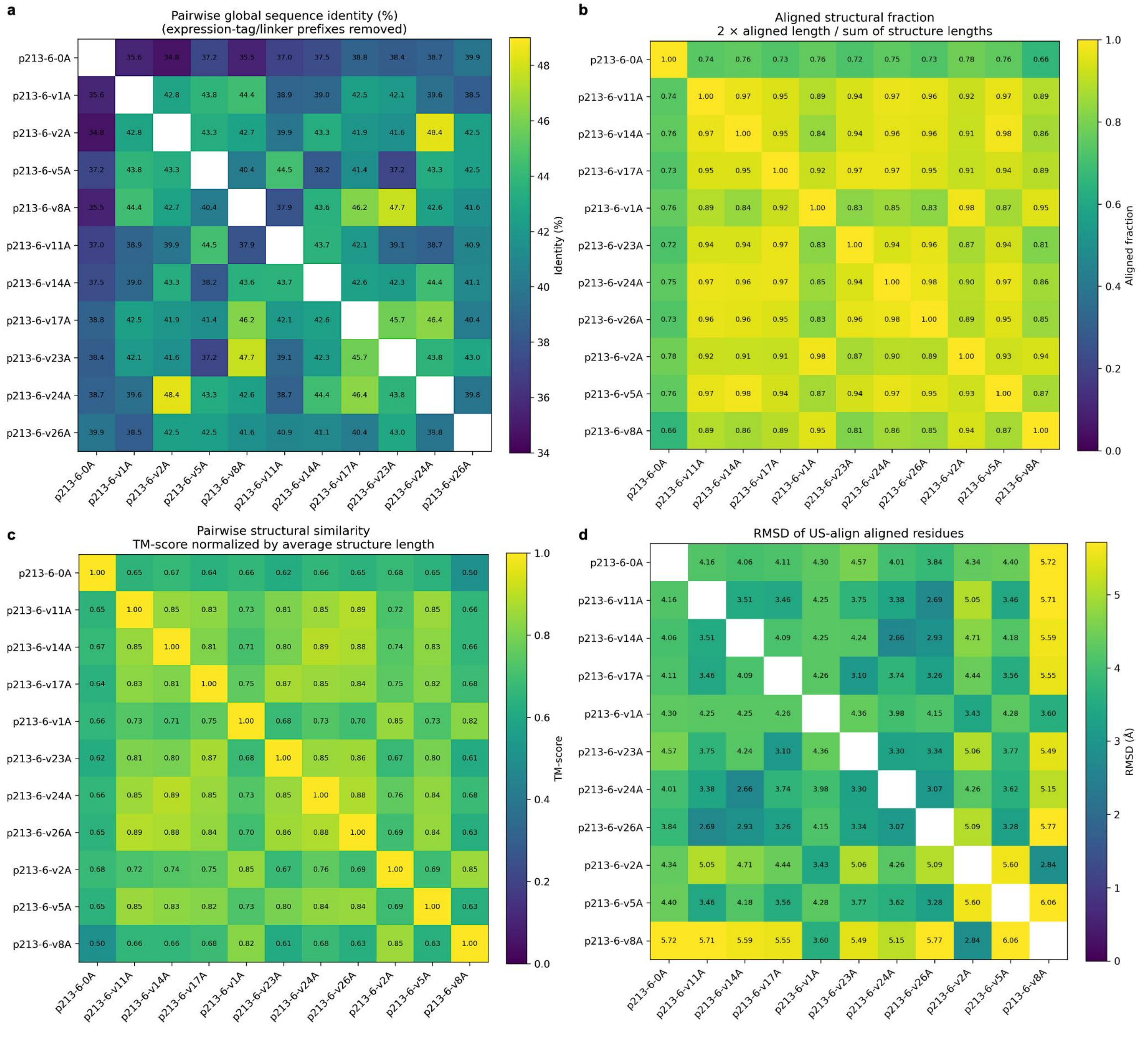


**Supplementary Fig. S13 Pairwise sequence and structural similarity of the RFdiffusion-generated crystal components. a**, Pairwise sequence identities were calculated after removing the N-terminal expression tags and linkers using Biopython PairwiseAligner[2]. Identity was defined as the number of exact residue matches divided by the total number of alignment columns, including gaps. Pairwise identities ranged from 34.8% to 48.4%, with a mean of 41.3%. **b-d**, Complete trimer assemblies were compared pairwise using US-align in multichain mode[3]. The aligned fraction, calculated as twice the number of aligned residues divided by the combined lengths of the two assemblies, ranged from 0.661 to 0.981, with a mean of 0.887. RMSDs over aligned residues ranged from 2.66 to 6.06 Å. Average-length-normalized TM-scores ranged from 0.504 to 0.891, with a mean of 0.750. Heatmap cells report pairwise sequence identity, aligned fraction, RMSD, or TM-score.

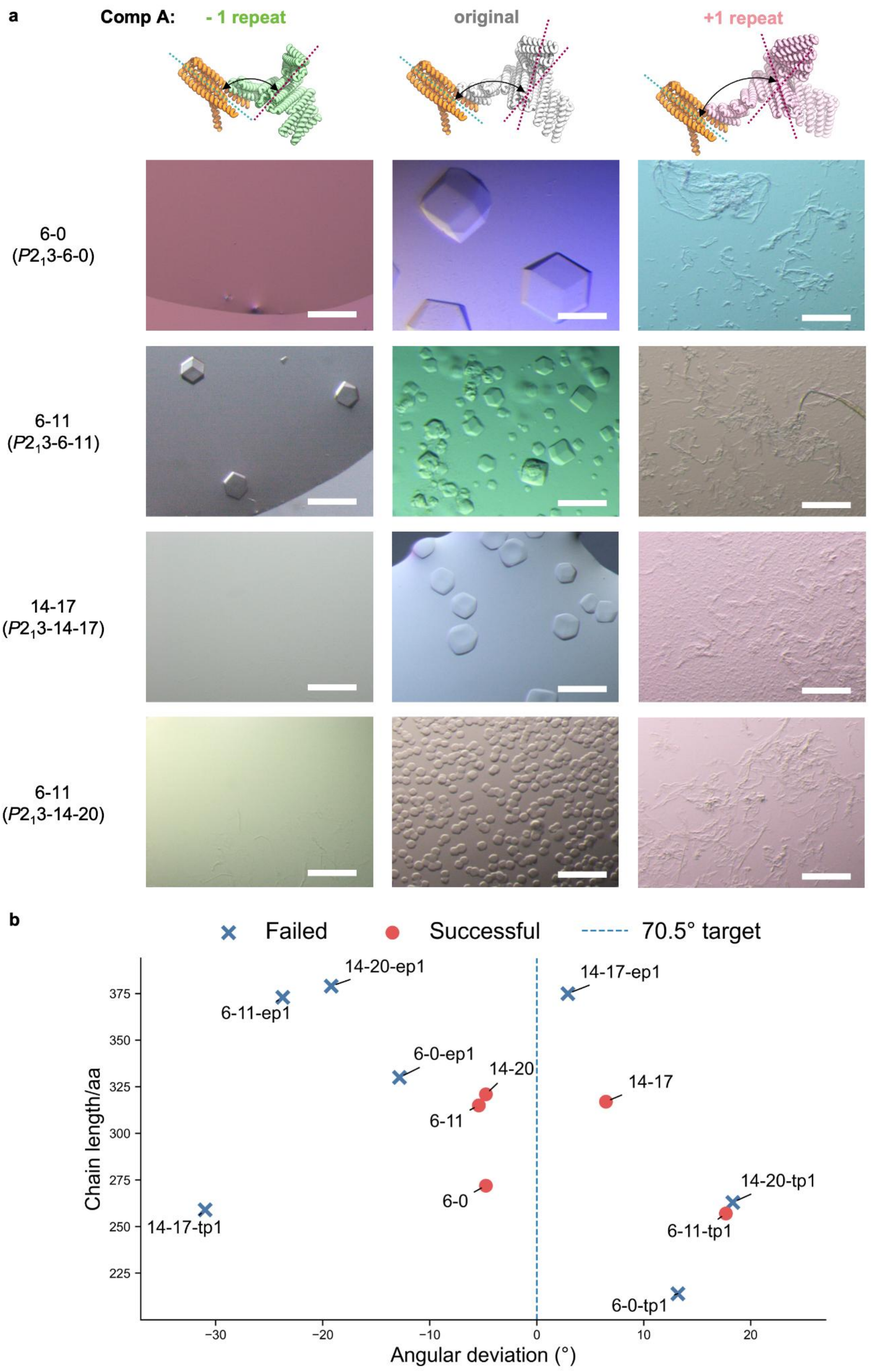

a
Comp A:
- 1 repeat
original
+1 repeat
6-0
($P2_13$-6-0)
6-11
($P2_13$-6-11)
14-17
($P2_13$-14-17)
6-11
($P2_13$-14-20)
b
Failed
Successful
70.5° target
Chain length/aa
Angular deviation (°)
14-20-ep1
6-11-ep1
14-17-ep1
6-0-ep1
14-20
6-11
14-17
6-0
14-17-tp1
14-20-tp1
6-11-tp1
6-0-tp1

**Extended Data Fig. S14 Accurate backbone geometry is important for lattice propagation.** **a**, Four $P2_13$ crystal designs were modified by adding or removing one de novo helical repeat from component A. In the upper schematic, the two interacting trimeric components are shown with their C3 axes indicated by dashed lines to illustrate deviations in the screw angle. Optical microscopy images of each modified design are shown below. Only one trimmed variant, derived from $P2_13$-6-11, crystallized after modification. Scale bar, 100 µm. **b**, Distribution of crystallization outcomes as a function of screw-angle deviation and chain length. Successful designs clustered near the target screw angle of 70.5° and at the lower end of the chain-length range. The suffixes tp1 and ep1 denote trimmed and extended variants, respectively. The failure of designs such as $P2_13$-14-17-ep1, which had one of the smallest screw-angle deviations, suggests that component flexibility may also contribute to assembly failure.

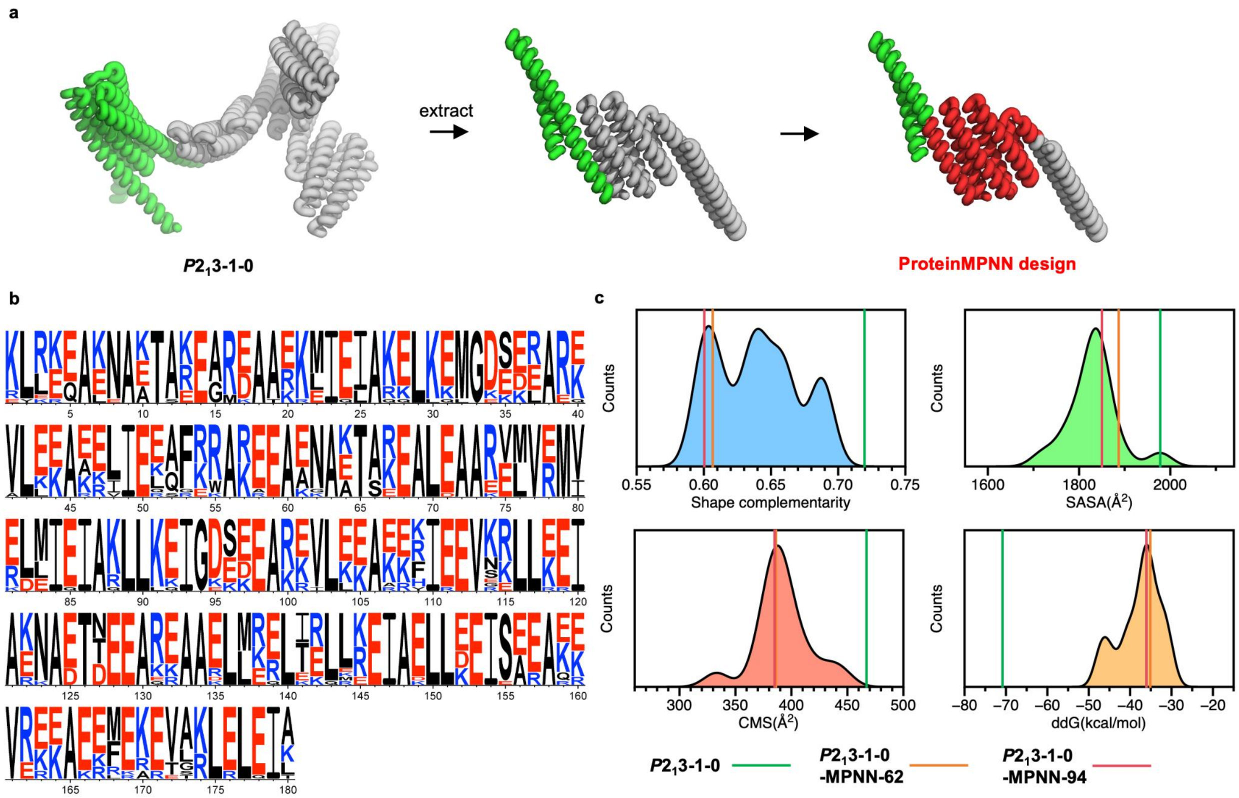


**Extended Data Fig. S15 ProteinMPNN redesign of protein sequences on a fixed protein backbone. a**, Single chains were extracted from the neighboring interacting component A and component B proteins. Backbone regions selected for sequence redesign are highlighted in red; residues forming the C3 homooligomeric interfaces were excluded from redesign. **b**, Sequence-logo representation of ProteinMPNN design outputs, showing amino-acid preferences at each redesigned position. **c**, Distributions of Rosetta metrics for the crystal-contact interfaces of experimentally tested designs. Individual lines indicate validated crystal-assembly designs, including the original *P*2₁3-1-0 design, which is shown for reference but excluded from the distribution calculation, and *P*2₁3-1-0-MPNN-62 and *P*2₁3-1-0-MPNN-94, which were included in the distribution calculation. ProteinMPNN redesigned crystal contacts featured relatively smaller interface and weaker interaction than the original design.

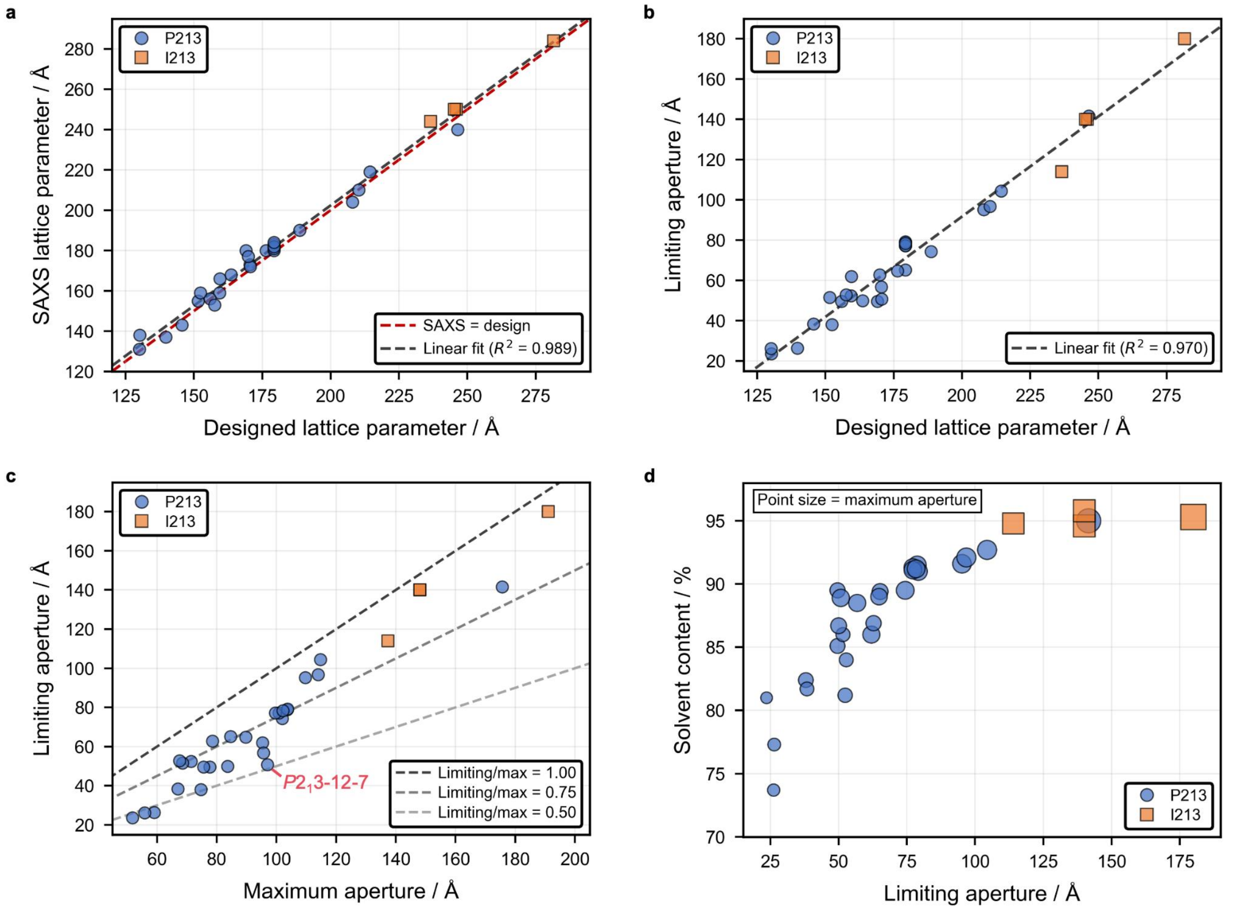


**Supplementary Fig. S16 Pore architecture and structural-property analysis of designed three-dimensional protein crystals.** All data correspond to Extended Data Table 1. **a**, SAXS-derived lattice parameters plotted against designed lattice parameters, showing close agreement between experimental and computational lattice sizes. **b**, Limiting aperture plotted against designed lattice parameters, showing that larger unit cells generally give rise to larger limiting apertures with an approximately linear relationship. **c**, Limiting aperture plotted against maximum aperture. Most designs fall within a limiting-to-maximum aperture ratio of 0.5–1.0. $P2_1$3-12-7, which has a low ratio, is labeled and may exhibit permanent guest-confinement properties. **d**, Solvent-content percentage plotted against limiting aperture, with maximum aperture indicated by point size. A positive correlation is observed, with limiting apertures ranging from 2 to 18 nm and solvent contents ranging from 74% to 95%.

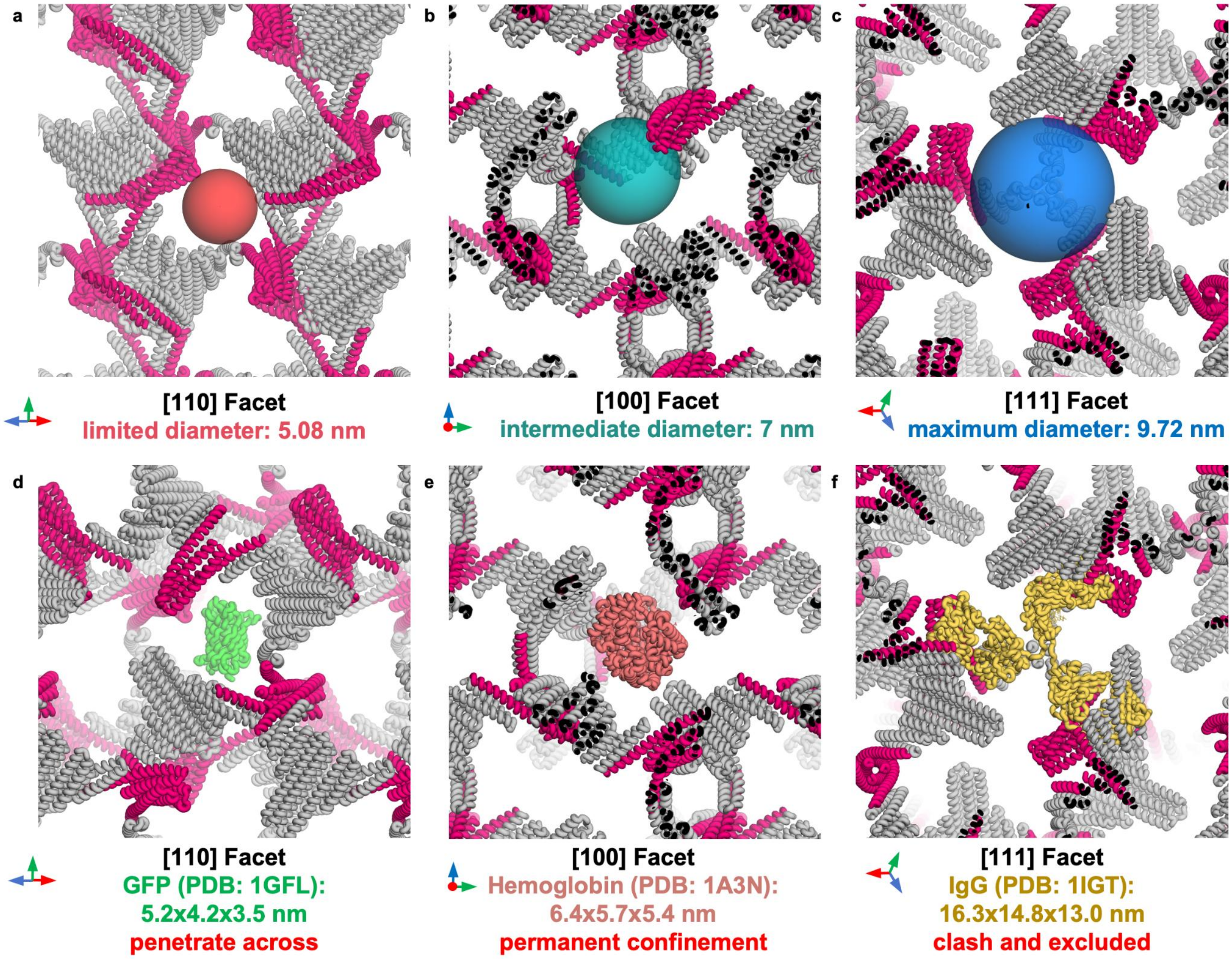


**Supplementary Fig. S17 Permanent guest confinement within the $P2_13$-12-7 lattice. a–c**, Sphere-based simulations illustrating the limiting diameter (**a**), an intermediate diameter (**b**) and the maximum diameter (**c**) of the crystal pore. **d–f**, Representative guest-protein simulations using GFP (**d**), hemoglobin (**e**) and IgG (**f**), with molecular dimensions estimated using Draw_Protein_Dimensions script (https://pymolwiki.org/Draw_Protein_Dimensions). GFP is smaller than the maximum pore diameter and has two dimensions below the limiting diameter, consistent with penetration through the lattice. IgG exceeds the maximum pore diameter in at least one dimension and is therefore predicted to be excluded. Hemoglobin is smaller than the maximum diameter but larger than the limiting diameter in at least two dimensions, suggesting that it could be incorporated during crystal assembly but remain permanently confined within the pore.

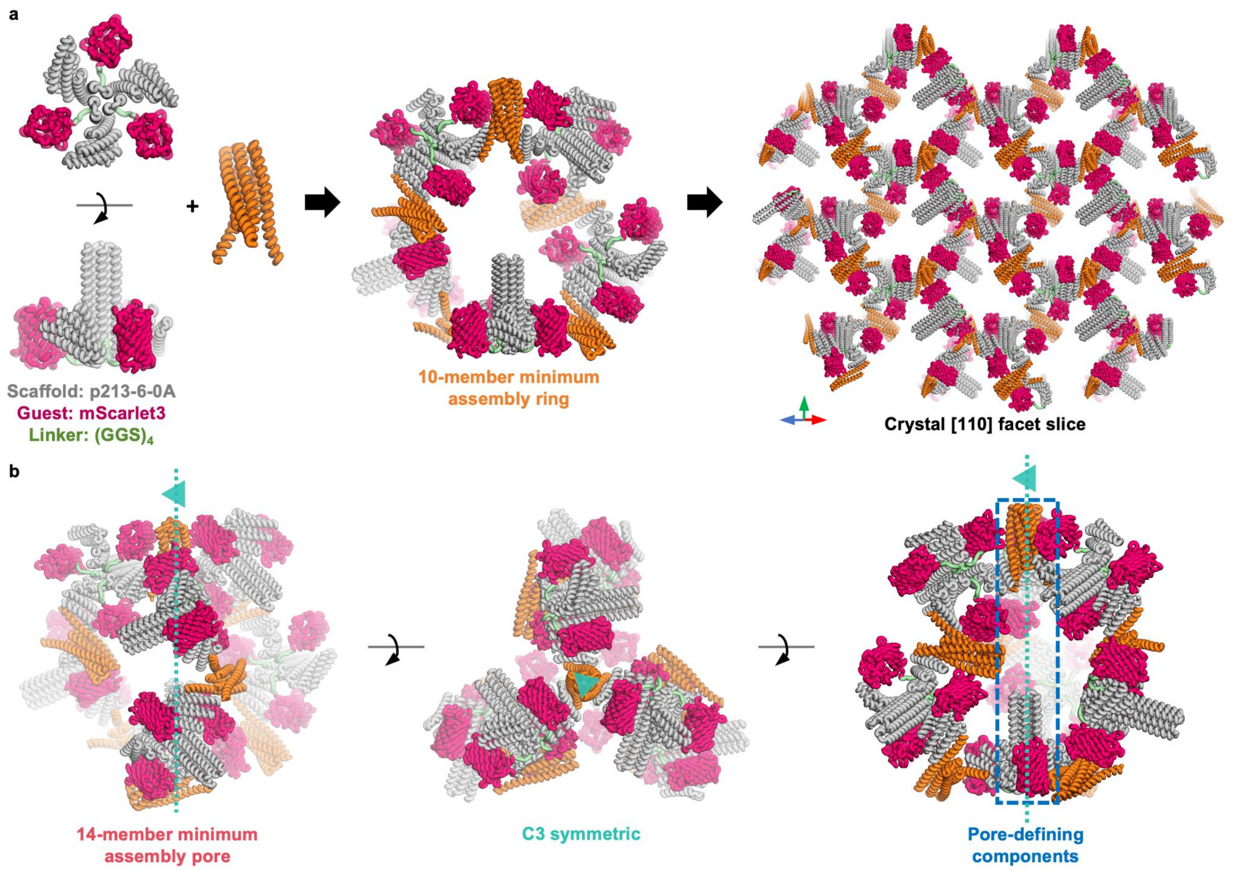


**Supplementary Fig. S18 Accommodation of GFP-like β-barrel fluorescent proteins within the *P*2$_1$3-6-0 crystal lattice by covalent fusion. a**, Schematic of crystal self-assembly with fused guest proteins. The *P*2$_1$3-6-0 component A was fused at its N terminus to mScarlet3 through a flexible (GGS)$_4$ linker and co-assembled with component B. The assembly first forms a 10-membered minimum assembly ring containing five copies of each component, which propagates into the crystal lattice, shown here as a [110] facet slice. The fused fluorescent proteins are multivalently displayed within the lattice, and the pore space allows positional flexibility of the linker-tethered guest proteins. **b**, Model of a 14-membered minimum assembly pore containing seven copies each of components A and B. The pore has C3 symmetry and is formed by three connected 10-membered rings. Within each pore, paired trimers of components A and B are aligned along the C3 axis, defining a cavity that accommodates three fused guest proteins per A-component trimer.

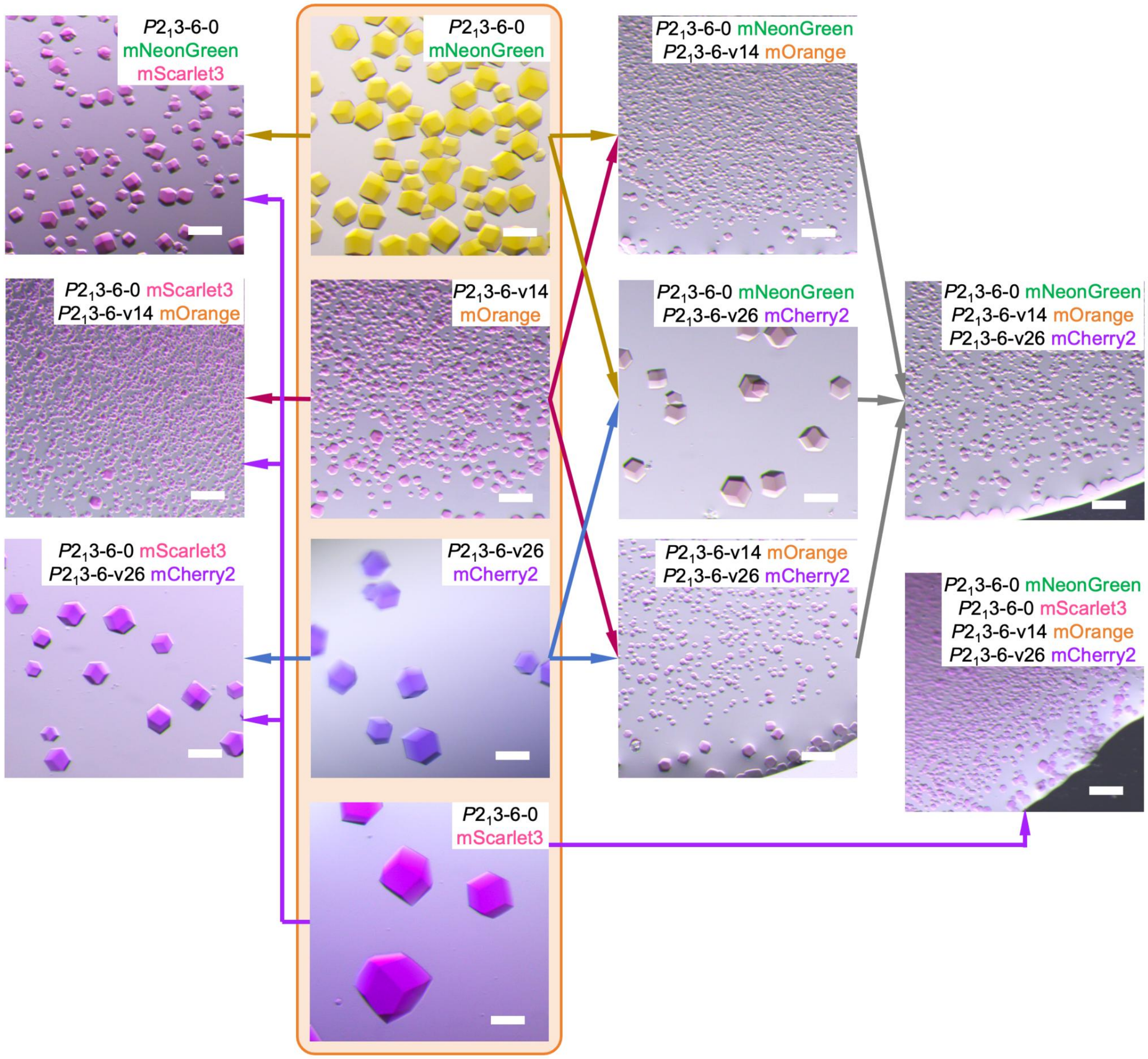


**Supplementary Fig. S19 Optical microscopy images of protein crystals incorporating fluorescent protein fusions.** Primary crystals, each containing a single fluorescent-protein fusion, are highlighted in the orange box. Surrounding images show co-assembled crystals generated by combining different primary crystal components, with the specific combinations indicated by arrows. The co-assembled crystals displayed mixed colors under bright-field microscopy.

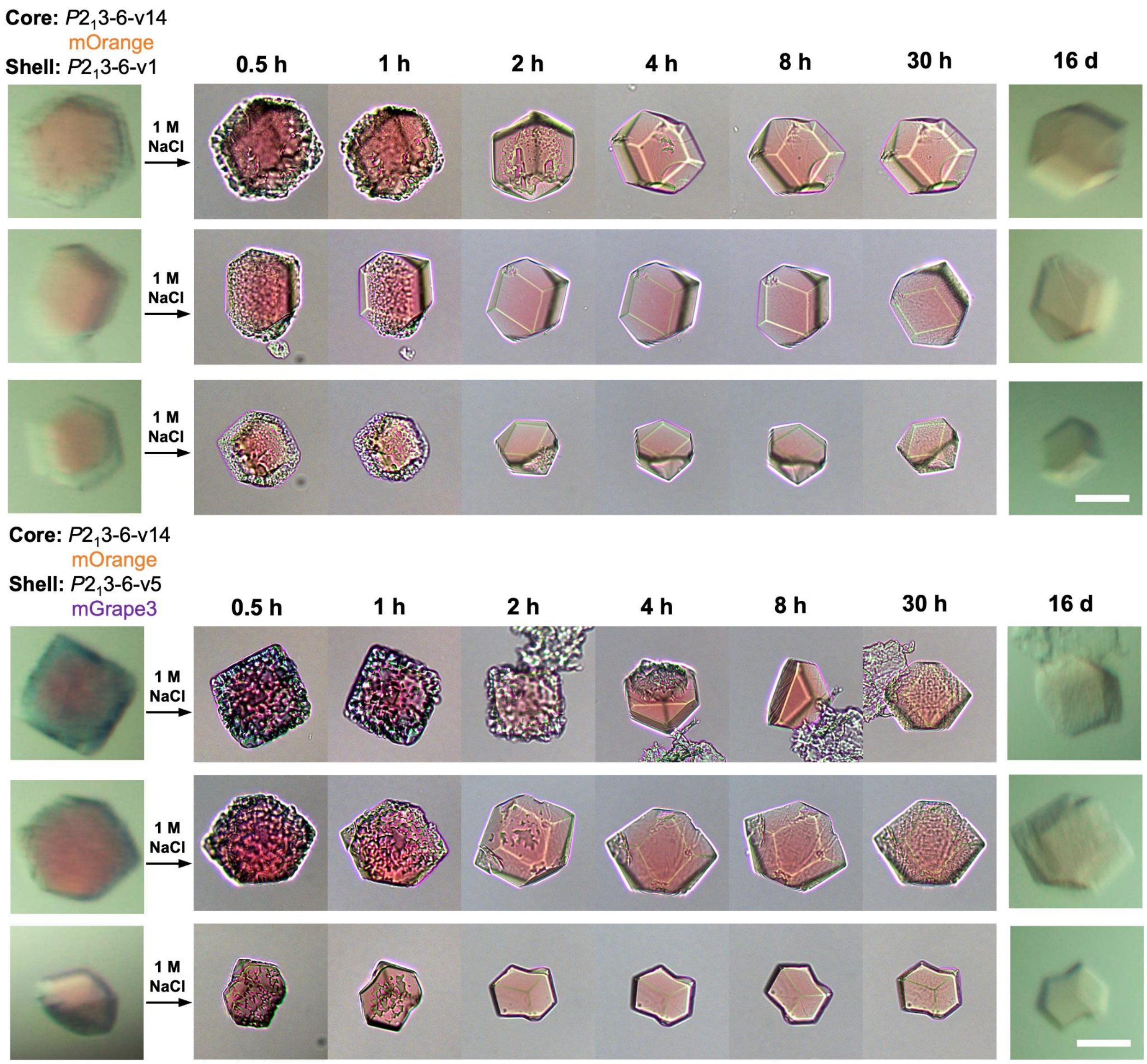


**Supplementary Fig. S20 Monitoring the disassembly of core–shell crystal assemblies.** Three representative crystals from two core–shell designs, $P2_13$-6-v14–mOrange@$P2_13$-6-v1 and $P2_13$-6-v14–mOrange@$P2_13$-6-v5–mGrape3, are shown during disassembly over time. At 1 M NaCl, the outer shell layer gradually disassembled over 2 h, whereas the inner core layer remained stably assembled and largely intact even after 16 d of incubation. Scale bars, 50 µm.

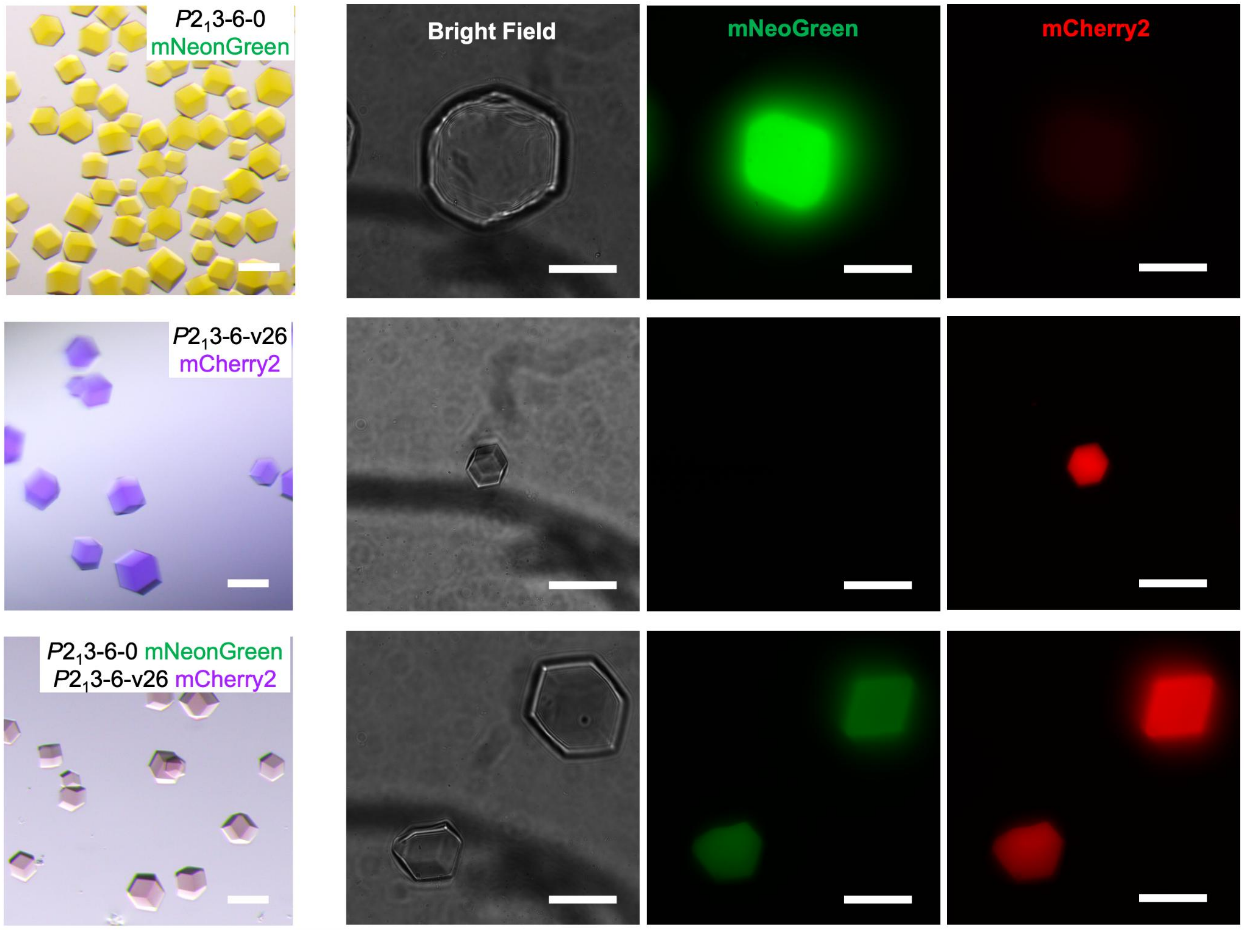


**Supplementary Fig. S21 Fluorescence microscopy images of co-assembled crystals containing fluorescent protein fusions.** From top to bottom, images show $P2_1$3-6-0–mNeonGreen crystals, $P2_1$3-6-v26–mCherry2 crystals and co-assembled crystals containing both constructs. Comparison of bright-field, green-channel and red-channel images shows that the co-assembled crystals incorporated both mNeonGreen and mCherry2, with both fluorescent components distributed uniformly throughout the crystals.

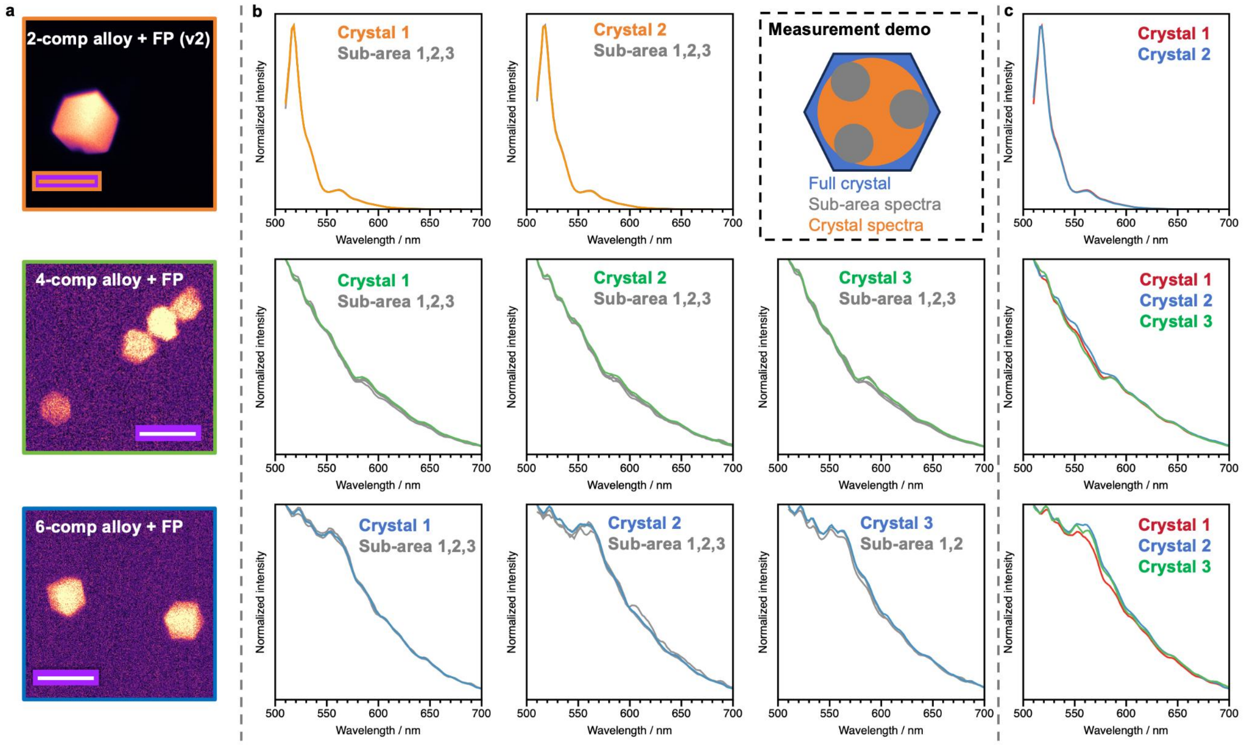


**Supplementary Fig. S22 Hyperspectral fluorescence analysis of protein crystal alloys. a**, Fluorescence images of 2-component, 4-component and 6-component crystal alloys incorporating fluorescent proteins. The 2-component alloy shown here is version 2, assembled from *P*2$_1$3-6-0–mNeonGreen and *P*2$_1$3-6-v23–mOrange, distinct from the version shown in Fig. 5o, which was assembled from *P*2$_1$3-6-0–mNeonGreen and *P*2$_1$3-6-v26–mCherry2. Scale bars, 50 µm. **b**, Fluorescence spectra measured from different subregions within individual crystals. Two to three crystals were measured for each sample, and spectra from each subregion were compared with the corresponding whole-crystal spectrum. Inset, schematic showing representative measurements from three subregions and the whole crystal. The consistent spectra across subregions indicate uniform distribution of the different fluorescent components within each assembled crystal alloy. **c**, Overlaid fluorescence spectra from different individual crystals, showing high spectral consistency across crystals.

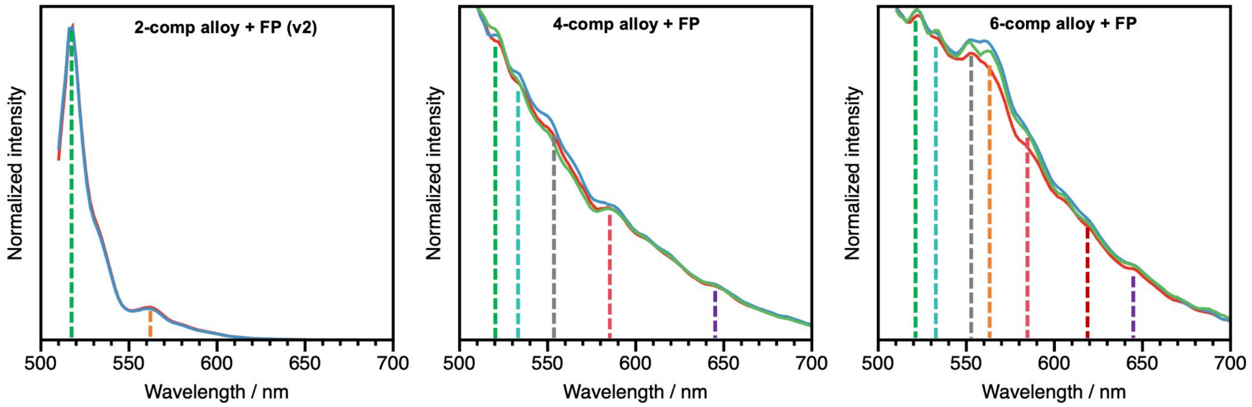


| | Excitation / nm | Emission / nm |
|---|---|---|
| mNeonGreen | 506 | 517 |
| mCitrine | 516 | 529 |
| mOrange | 548 | 562 |
| mScarlet3 | 569 | 592 |
| mCherry2 | 589 | 610 |
| mGrape3 | 608 | 646 |

Crosslinking reagent
glutaraldehyde: ~560 nm

**Supplementary Fig. S23 Assignment of fluorescent-protein emission peaks in the spectra of 2-component, 4-component and 6-component crystal alloys incorporating fluorescent proteins.** The table below lists the excitation and emission maxima of all six fluorescent proteins. Characteristic emission wavelengths of the constituent fluorescent-protein fusions identified in the spectra are indicated by dashed lines of the corresponding colors. Possible Förster resonance energy transfer (FRET) between fluorescent guests may have contributed to the reduced signal-to-noise ratio observed in the spectra. Crystals were mildly crosslinked with glutaraldehyde before characterization, resulting in an additional peak around 560 nm, indicated by grey dashed lines[4].

**Supplementary Table S1. Experimentally tested protein sequences in this study**

N terminal Met are included in all sequences; His-tag, Trp (added for A280 absorbance) and flexible linkers are underscored; Fluorescent protein fusions and corresponding engineered regions are highlighted.

| Name | Sequence* | Notes |
|---|---|---|
| p213-w2 | MEADLRFVAVLQKLNIELARLLLEAVARLQELNIDLVRKTSELT<br>DEKTIREEIRKVKEESKRIVEKAEELIRLAKLVSDVAVKLAELA<br>TKARDEEVRRIAAEVADQAMRLATEAAESGSQKCLELATLVLEA<br>VRAMAKAAKKSTDEEKIRKLLEEARQAIEEAQRACRSNDEKRLA<br>RAALKVARLLVKAAKEAGDPKAVEKAAKAALKVAEEANKKGDIQ<br>ILTEALEAAAEVARRLAEVARKRGEPRLEESAIKIALDALKLST<br>EALEKLTASLRAITEELKKNPSEDALVEHNRAIVEHNDMIRVNN<br>FLIAVVLELIVLRIKGSWSGLEHHHHHH | $P2_13$ design, single chain |
| p213-w3 | MDRELEVVALLQRLNIELARKLLEAVARLQELNIDLVRKTSELT<br>DEKTIREEIRKVKEESKRIVEEAEEEIRKAKKMSEILVRAADKD<br>AESIREAVRLAEELLRRDPSTEAEDLLRFAIQMAVEAPDPEAIR<br>EAVRAARELLRENPSEEAFRLLERAIEAAAQSPDPRARKEALEA<br>AEEMAREAKKELEEAEKRGDPKRALRAVEIVVRAAELLMKIAKA<br>EGSEEAKERAAKVAAEAARLAKRVAELAAKAGDKDVAKKALELA<br>LRALEIMLEILRDMLRKLKESLEELKKNPSEDALVRNNELIVKV<br>LEIIVMVLEAIAAVLKINALLVGSWSGLEHHHHHH | $P2_13$ design, single chain |
| p213-w7 | MDDELEAVAELQELNIQLAKKLLEAVARLQELNIDLVRKTSELT<br>DEKTIREEIRKVKEESKRIVEEAEEEIRRAKQISKAIEEMARLL<br>RRAKNAETPEEAARAAKELIKIAEKLQREGNKEQAEKVLRKATE<br>IIKRAAEMAEKEAKNARTPEEALRAAEVLVRLIKLLIEIAKLLQ<br>EQGNKEEAEKVLREAEELIDRVLELLEKIAENADTEELAKRAAE<br>LIKRLIELLKEIAKLLREAGLEDEAEKVEEKAKEMEERVDRLEL<br>KIALDELEKALQELREMLRKLKESLEELKKNPSEDALVRNNELI<br>VEVLRVIVEVLSIIAKVLKLNAKLAGSWSGLEHHHHHH | $P2_13$ design, single chain |
| p213-w2-2A | MRGHHHHHHGSWSGEADLRFVAVLQKLNIELARLLLEAVARLQE<br>LNIDLVRKTSELTDEKTIREEIRKVKEESKRIVEKAEELIRLAK<br>LVSDVAVKLAELATKARDEEVRRIAAEVADQAMRLATEAAES | Design $P2_13$-w2 split at the 2nd loop of the connector, component A |
| p213-w2-2B | MQKCLELATLVLEAVRAMAKAAKKSTDEEKIRKLLEEARQAIEE<br>AQRACRSNDEKRLARAALKVARLLVKAAKEAGDPKAVEKAAKAA<br>LKVAEEANKKGDIQILTEALEAAAEVARRLAEVARKRGEPRLEE<br>SAIKIALDALKLSTEALEKLTASLRAITEELKKNPSEDALVEHN<br>RAIVEHNDMIRVNNFLIAVVLELIVLRIKGSWSGLEHHHHHH | Design $P2_13$-w2 split at the 2nd loop of the connector, component B |
| p213-w2-3A | MRGHHHHHHGSWSGEADLRFVAVLQKLNIELARLLLEAVARLQE<br>LNIDLVRKTSELTDEKTIREEIRKVKEESKRIVEKAEELIRLAK<br>LVSDVAVKLAELATKARDEEVRRIAAEVADQAMRLATEAAESGS | Design $P2_13$-w2 split at the 3rd loop of the connector, component A |

| | | |
|---|---|---|
| | QKCLELATLVLEAVRAMAKAAKK | |
| p213-w2-3B | MEEKIRKLLEEARQAIEEAQRACRSNDEKRLARAALKVARLLVKAAKEAGDPKAVEKAAKAALKVAEEANKKGDIQILTEALEAAAEVARRLAEVARKRGEPRLEESAIKIALDALKLSTEALEKLTASLRAITEELKKNPSEDALVEHNRAIVEHNDMIRVNNFLIAVVLELIVLRIKGSWSGLEHHHHHH | Design $P2_13$-w2 split at the 3rd loop of the connector, component B |
| p213-w3-7A | MRGHHHHHHGSWSGDRELEVVALLQRLNIELARKLLEAVARLQELNIDLVRKTSELTDEKTIREEIRKVKEESKRIVEEAEEEIRKAKKMSEILVRAADKDAESIREAVRLAEELLRRDPSTEAEDLLRFAIQMAVEAPDPEAIREAVRAARELLRENPSEEAFRLLERAIEAAAQSPDPRARKEALEAAEEMAREAKKELEEAEKRGDPKRALRAVEIVVRAAELLMKIAKAE | Design $P2_13$-w3 split at the 7th loop of the connector, component A |
| p213-w3-7B | MEEAKERAAKVAAEAARLAKRVAELAAKAGDKDVAKKALELALRALEIMLEILRDMLRKLKESLEELKKNPSEDALVRNNELIVKVLEIIVMVLEAIAAVLKINALLVGSWSGLEHHHHHH | Design $P2_13$-w3 split at the 7th loop of the connector, component B |
| p213-w7-4A | MRGHHHHHHGSWSGDDELEAVAELQELNIQLAKKLLEAVARLQELNIDLVRKTSELTDEKTIREEIRKVKEESKRIVEEAEEEIRRAKQISKAIEEMARLLRRAKNAETPEEAARAAKELIKIAEKLQREGNKEQAEKVLRKATEIIKRAAEMAEKEAKNARTPEEALRAAEVLVRLIKLLIEIAKLLQEQ | Design $P2_13$-w7 split at the 4th loop of the connector, component A |
| p213-w7-4B | MKEEAEKVLREAEELIDRVLELLEKIAENADTEELAKRAAELIKRLIELLKEIAKLLREAGLEDEAEKVEEKAKEMEERVDRLELKIALDELEKALQELREMLRKLKESLEELKKNPSEDALVRNNELIVEVLRVIVEVLSIIAKVLKLNAKLAGSWSGLEHHHHHH | Design $P2_13$-w7 split at the 4th loop of the connector, component B |
| p213-w7-5A | MRGHHHHHHGSWSGDDELEAVAELQELNIQLAKKLLEAVARLQELNIDLVRKTSELTDEKTIREEIRKVKEESKRIVEEAEEEIRRAKQISKAIEEMARLLRRAKNAETPEEAARAAKELIKIAEKLQREGNKEQAEKVLRKATEIIKRAAEMAEKEAKNARTPEEALRAAEVLVRLIKLLIEIAKLLQEQGNKEEAEKVLREAEELIDRVLELLEKIAEN | Design $P2_13$-w7 split at the 5th loop of the connector, component A |
| p213-w7-5B | MEELAKRAAELIKRLIELLKEIAKLLREAGLEDEAEKVEEKAKEMEERVDRLELKIALDELEKALQELREMLRKLKESLEELKKNPSEDALVRNNELIVEVLRVIVEVLSIIAKVLKLNAKLAGSWSGLEHHHHHH | Design $P2_13$-w7 split at the 5th loop of the connector, component B |
| p213-1-0A (p213-w7-6A) | MRGHHHHHHGSWSGDDELEAVAELQELNIQLAKKLLEAVARLQELNIDLVRKTSELTDEKTIREEIRKVKEESKRIVEEAEEEIRRAKQISKAIEEMARLLRRAKNAETPEEAARAAKELIKIAEKLQREGNKEQAEKVLRKATEIIKRAAEMAEKEAKNARTPEEALRAAEVLVRLIKLLIEIAKLLQEQGNKEEAEKVLREAEELIDRVLELLEKIAENADTEELAKRAAELIKRLIELLKEIAKLLREA | Design $P2_13$-w7 split at the 6th loop of the connector, design $P2_13$-1-0, component A |
| p213-1B | MEDEAEKVEEKAKEMEERVDRLELKIALDELEKALQELREMLRK | Design $P2_13$-w7 split at |

| | | |
|---|---|---|
| (p213-w7-6B) | LKESLEELKKNPSEDALVRNNELIVEVLRVIVEVLSIIAKVLKLNAKLAGSWSGLEHHHHHH | the 6th loop of the connector, component B, utilized across all *P*2₁3-1-* assemblies |
| p213-1-9A | MRGHHHHHHGSWSGSLELEAVALLQELNIELARLLLEAVARLQELNIDLVRKTSELTDEKTIREEIRKVKEESKRIVEEAELAIRAAKAISEAIAAAARAAQKGNEEVAKAALFVLKAALELAKAKRDKEAIERVERIAREAKKAAELAEKGDDARALEILLELAVKIIEELIRAAKEAQEQGNKEEAEKILRKAEELIDRALELAERIAENARTEEAAKRAAEMIKKLIELLKEIAKLLREA | Design *P*2₁3-1-9, component A |
| p213-1-11A | MRGHHHHHHGSSGSEELKAVALLQRLNIELARLLLEAVARLQELNIDLVRKTSELTDEKTIREEIRKVKEESKAIVLWAELAILAAKAASLVIAAEALLKAGDKEEAEKVLRKAAELIREATRKAEELAKNAATPEVALLAAEILVRLIKLLIEIAKLLQEAGNKEEAEKVLREATELIKRVTELLEKIAKNSDTPELALRAAELLVRLIKLLIEIAKLLREQGNKEEARKVLKEAQELIKRVAELLEKIAKNAKTPEEALRAAELLVRLIKLLIEIAKLLQESGNKEEAEKVLREAEELIDRVLELLEKIAENADTEELAKRAAELIKRLIELLKEIAKLLREA | Design *P*2₁3-1-11, component A |
| p213-1-21A | MRGHHHHHHGSSGPDLFLKALELLIKLAESLAETARERGDEKALEEAARIAEKAAELAEKLARKARKEGNLELALQALRAMVEAARVLAEIARERGNEELLKYAWELARRAAEQALQIAAEAALRGNEELALIALEILVEALKVLSHIAREKGDPKLLEESMLWQRILVALVEALLALLEGDAETAERAAKEAARIAEEAGLGEALEAVAALAIAIAKFAQEAGQKEEAEKVLRKAEEIIDKALEILEKQAENAETEEEAKKAAEKIKKLIELLKIIAKLLREA | Design *P*2₁3-1-21, component A |
| p213-1-26A | MRGHHHHHHGSWSGSEELKAVALLQRLNIELARKLLEAVARLQELNIDLVRKTSELTDEKTIREEIRKVKEESKRIVERAEAEIQEAKAESELIAAKIAMEHNPSEDALKLAVEAIKKAIEAAKEMQRLGDKERAEKILRKAEEAIDEALEAAERLAENASTEEQAKKAAELIKRLIELLKEIAKLLREA | Design *P*2₁3-1-26, component A |
| p213-6-0A | MRGHHHHHHGSWSGSKELEAVARLQELNIELARKLLEAVARLQELNIDLVRKTSELTDEKTIREEIRKVKEESKRIVEEAEEEIRRAKAISLHIAIRALEDAARQVEEAIKRNPDNDEAVETAVRIARLLKEAAEKAQELAKRLGDPELLKDALRALEVAVRAVELAIRSNPNNEEAVRTAVRLAEELMKVARELIERAEKTGDPELLKLARRAAEMAVRAVEMAIKANPDDDKAVEAAVRLARVLKEIAERLQEEAQKRGDEELLKEAERALEVAVRAVELAKKS | Design *P*2₁3-6-0, component A |
| p213-6B | MEEAEETAKRLARELTRVKLLQALAELEKALRELKKSLDELERSLEELEKNPSEDALVENNRLNVENNKIIVEVLRIIAEVLKLNAQAGGSWSGLEHHHHHH | Design *P*2₁3-6-0, component B, utilized across all *P*2₁3-6-* assemblies |
| p213-6-1A | MRGHHHHHHGSWSGSEEERKVKELQKLNIELARILLEAVAKLQE | Design *P*2₁3-6-1, |

| | | |
|---|---|---|
| | LNIDLVRKTSELTDEKTIREEIKKVKIQSRVIVQNAELAIEVAK<br>AISEIIAGGDPKQAIEKIAEKAEALLREVERLGGPPEALREAAQ<br>VAKEAMEAAERAARKAGAPPEVQEKARILGELLELIARVARVKR<br>RIKENPDDDKAVEEAVRLARELKKVAERAQEKAQKTGDEKMLEA<br>AEKALEVAVRAVELAKKS | component A |
| p213-6-2A | MRGHHHHHHGSWSGSEEERKVKRLQKLNIELARILLEAVARLQE<br>LNIDLVRKTSELTDEKTIREEIKRVKARSEAIVKEAEIAIEEAK<br>ARSEEEAGGDPKEAKEKLIRKLKEAAERLQELAKQLGDPELLKK<br>ALEMLEKAVRVAEELIKRNPDDDKAVELAVRLARALKKVAEELQ<br>ERAQKTGDEELLKLAERALEVAVRAVELAKKS | Design $P2_13$-6-2, component A |
| p213-6-8A | MRGHHHHHHGSWSGSDELEAVHLLQQLNIELARLLLEAVARLQE<br>LNIDLVRKTSELTDEKTIREEIRKVKEQSKRIVQVAEAAIRAAE<br>AASEAISRIAKTDDPDSLRKVVRELARKLKQIVEEAARLNGPGE<br>MIELLARFAIELLKRAAEFAERAMTDERVREQLRAEALVAAAEI<br>LKLAARALQEEAKQTGDPELLKQALQLLEEAVRMVEEAIKRQPD<br>DDKAVETAVRLARELKKVAEELQERAQKTGDEELLKLAERALEV<br>AVRAVELAKKS | Design $P2_13$-6-8, component A |
| p213-6-10A | MRGHHHHHHGSWSGSEHAKAVYKLQRLNIELARKLLEAVARLQE<br>LNIDLVRKTSELTDEKTIREEIRKVKEESKRIVEEAEKQIEAAK<br>LISEIIYILAEARASGTEESLRQAIERVLQIARKGDDREALRAA<br>AKVIEEIARLSGSEQAKIEAARALKEIAEILQEKAKKTGDPELL<br>KEALEALERAARLLEEAIKRNPDDDKAVEMAVRIARELKKVAEE<br>LQERAQKTGDEELLKLAERALEVAVRAVELAKKS | Design $P2_13$-6-10, component A |
| p213-6-11A | MRGHHHHHHGSWSGSEQEEIVNRLQRLNIELARQLLEAVARLQE<br>LNIDLVRKTSELTDEKTIREEIRKVKEESKRIVEEAEIRIRAAE<br>VISEALALQAELPSEEAKKAVEEIEEAAKAAERAAEKGQTEVAK<br>QALELLKRAIEEAKRKRSEEALRKVERIARAAKLIARAAELKAE<br>AERLIKKARKTGDPELLRKALEALEEAVRAAEEAIKTNPDDDMA<br>VKMAVELARELKKVAEELQERAKKTGDPELLKLALRALEVAVRA<br>VELAIKSNPDNDEAVETAVRLARELKKVAEELQERAQKTGDEEL<br>LKLAERALEVAVRAVELAKKS | Design $P2_13$-6-11, component A |
| p213-6-12A | MRGHHHHHHGSWSGSEQEEIVNRLQRLNIELARQLLEAVARLQE<br>LNIDLVRKTSELTDEKTIREEIRKVKEESKRIVEEAEIRIRAAE<br>VISEALALQAELPSEEAKKAVERIEEAAKAAERAAEQGKTEVAK<br>LALKVLKKAIEMAKRLRSEEALKIVEAIAEAAKRAAKEAEKGDT<br>EEAKKALRLAEELLRLIEEVFKAEERIKTNPDDDEAVERAVRLA<br>RKLKEVAERLQEQAKKKGEPALLKLALIALEIAVRAVELAIKSN<br>PDNDEAVETAVRLARELKKVAEELQERAQKTGDEELLKLAERAL<br>EVAVRAVELAKKS | Design $P2_13$-6-12, component A |
| p213-12-7A | MRGHHHHHHGSWSGSKELEAVARLQRLNIELARKLLEAVARLQE<br>LNIDLVRKTSELTDEKTIREEIRKVKEESKRIVLRAKAQILAAK<br>AQSAIIAAEAALTAGDPETAREAVREALELVEQLRKLAKKAGDK<br>EALEAAAELARQVAKVAREVGSPETALEALKVAAEAAKESGNKE | Design $P2_13$-12-7, component A |

| | | |
|---|---|---|
| | EAEKILREATELIKRATEELEKRAKNARTSEEAKRAAEKLKKLIEMLKEIAKLLEEA | |
| p213-12B | MEDEAEKVKEEAKELERRVTKLEIEILLRELKASTAELKRATASLRAITEELKKNPSEDALVEHNRAIVEHNAIIVENNRIIAKVLEAIVRAIKGSWSGLEHHHHHH | Design $P2_13$-12-0, component B, utilized across all $P2_13$-12-* assemblies |
| p213-14-0A | MRGHHHHHHGSWSGSEELKAVADLQKLNIELARKLLEAVARLQELNIDLVRKTSELTDEKTIREEIRKVKEESKRIVEEAEKEILQAKVISLEIAVRAFEEAIKKDPDSDSAVEAAVRLARDLKKAAERLQEEAKRKGDPELLKSALRALEVAVRAVELAIRSNPDNEEAVETAKRLAKELQKVAKELLERAKKTGDPELLKLAKRALEVAVRAAEAAAKAMEERIKKNPDDDRAVEEAVKIARVLKELAELLQEVAKALGDEEALKLAERALEVAVRAVELAKKS | Design $P2_13$-14-0, component A |
| p213-14B | MEEAEETAKRLAEELKKVKILQALAELRKSTAELKRATASLRAITEELKKNPSEDALVEHNRAIVEHNAIIVENNRIIAKVLELIVRAIKGSWSGLEHHHHHH | Design $P2_13$-14-0, component B, utilized across all $P2_13$-14-* assemblies |
| p213-14-1A | MRGHHHHHHGSWSGSEELKAVAELQRLNLELALKLLEAVARLQELNIDLVRKTSELTDEKTIREEIEKVRDESALIVAEAAVEIIKAAIESAKIAQEAGSKEAAEKILRAAERAIEEIRRLLEELRKNAYTPEARERAKKLLERLVKLLAEIAKLLEEQGNLEKALKVAEKAVREVEQLIKENPDDDKAVELAVELARLLKEIAERLQEEAKKTGDERLLKLAERALEVAVRAVELAKKS | Design $P2_13$-14-1, component A |
| p213-14-2A | MRGHHHHHHGSWSGSEELKAVAELQRLNLELALKLLEAVARLQELNIDLVRKTSELTDEKTIREEIEKVRDESALIVAEAAVEIIKAAIESARIAFEAGSDEAAAKILQAALRAIEEIARLLFELAQNAYTPEARKRAEELLERLAKLLKEAAEMAKEMIKKNPDDDKAVKLAVEIAKALKKMAEMLQRLAKQTGDERLLKLAEEALEVAVQAVELAKKS | Design $P2_13$-14-2, component A |
| p213-14-3A | MRGHHHHHHGSWSGSEELKAVAELQRLNLELALKLLEAVARLQELNIDLVRKTSELTDEKTIREEIEKVRDESALIVAEAAVEIIKAAIESAKIAQEAGSKEAAEKILRAARRAIREITRLLEELAKNAYTPEAAARAAELLAELIKLLIEIARLLKEQGNKEEARKVLEEAKELAKRAAEIVKKLVRKAEERIKTNPDDDRAVELAVRLAKIAKKLAEAAQEIAKQTGDEEMLKLAEELLEVAVRAVELAKKS | Design $P2_13$-14-3, component A |
| p213-14-9A | MRGHHHHHHGSWSGSLELEAVAVLQKINIELARLLLEAVAQLQELNIDLVRKTSELTDEKTIREEIRKVKERSKAIVELAEEAIELAKEASEAVTRIAKTDDPDELRKVARELAERAEALVREAARKDLPEEAIVKFAEYAMKLLVTATELAMSFMTDKEVVTELAREAIESLRRIAEYVLRAATDGETARRAMESGEKEIRRLEKAAQKYTTDPRSELKLRRRALEAMIRLVELMIKKNPDDDKAVEKAVELAREAKRAAEEAQEQAKKTGDPEMLKEALKMLELAVRAVELAIKSNPDNDEAVETAVRLARELKKVAEELQERAKKTGDEELLKLAERALEVAVRAVELA | Design $P2_13$-14-9, component A |

| | | |
|---|---|---|
| | KKS | |
| p213-14-10A | M<u>RGHHHHHHGSWSG</u>SLELEAVAVLQKINIELARLLLEAVAQLQE<br>LNIDLVRKTSELTDEKTIREEIRKVKERSKAIVELAEEAIELAK<br>EASEAVTRIAKTDDPDELRKVARELAERAEALVREAARKDLPEE<br>AIVKFAEYAMKLLATAAELAMKFMRDARVVLELAREAEESVKRI<br>AEYVKRAATDGETKRRAEKSGLEERLRIAEAAVRALEEALKQNP<br>DDDKAVKALVRAARQLKRIAEEMQEEAKKQGDPELLKLALRALE<br>LAVRAVELAIKSNPDNDEAVETAVRLARELKKVAEELQERAKKT<br>GDEELLKLAERALEVAVRAVELAKKS | Design $P2_13$-14-10, component A |
| p213-14-17A | M<u>RGHHHHHHGSWSG</u>SEEQQRVAALQKLNIELARALLEAVARLQE<br>LNIDLVRKTSELTDEKTIREEIRKVKEESKKIVKAAEIIIRIAK<br>YASIAINKNGSDEARQAVKEIARLAKEALEEGCCDAAKFAIRAL<br>ELLARVFDGSDVQRLAEEAIEKIAETAARNGGDETAKEAARALK<br>RLAEELIKKARKTGDPELLRQALRALEAAARAAELAIRFNPDDD<br>EAVELAVRLARELKKVAEELQERAKKTGDPELLKLALRALEVAV<br>RAVELAIKSNPDNDEAVETAVRLARELKKVAEELQERAKKTGDE<br>ELLKLAERALEVAVRAVELAKKS | Design $P2_13$-14-17, component A |
| p213-14-20A | M<u>RGHHHHHHGSWSG</u>SYQLEIVARLQELNIRLARQLLEAVARLQE<br>LNIDLVRKTSELTDEKTIREEIRKVKEESKRIVEEAELLIEAAR<br>VASEILTLAAKAEEEGNPELRDSARKLYQALKEALEIAKRAGEP<br>EVIREVTREAEEAAKVIKEAIEAEKQGDRERHKKAQVRVLILIA<br>EILKKAAEAAIKKARKTGDPELLRKALELLEKAARAAEEAIKRN<br>PDDDKAVEMAVRLARELKKVAEELQERAKKTGDPELLKLALRAL<br>EVAVRAVELAIKSNPDNDEAVETAVRLARELKKVAEELQERAKK<br>TGDEELLKLAERALEVAVRAVELAKKS | Design $P2_13$-14-20, component A |
| p213-6-v1A | M<u>RGHHHHHHGSWSG</u>ETIKKIIKLLSLITLLLIREAIKKVKLGEL<br>SAEEVKIIVEAAALQIELDKENAPELLEELKKLAKEVVRKIPEV<br>VIHIARKIEELVKAAPKENIPHLEELVEELIEEVEREGVLTLEL<br>VIEIGRAVLTLLKKGSSERTVELAVRLARILKRMAEELQKRAQE<br>TGDERLLKKAEEALEVAVELVEAAKEA | Design $P2_13$-6-v1, component A, isomorphous lattice to $P2_13$-6-0 |
| p213-6-v2A | M<u>RGHHHHHHGSWSG</u>DLALLAKTIAEIIVILAEVLVRLGALTPEL<br>AKRLAELVAKLLKAFPETAPQAIRFAEVLLEAVRRDPRFLPVLV<br>ELLEAMAEEVERLTPEDKRRLARVARAALEAAREQGAPLEVLLR<br>IERVLARFLLALVEDGDEEAVEEAVRVARRLKELAEELQKRAQE<br>TGDEELLKLAEEALEEAVRLVEAAKRF | Design $P2_13$-6-v2, component A, isomorphous lattice to $P2_13$-6-0 |
| p213-6-v5A | M<u>RGHHHHHHGSWSG</u>MEARLRAILLALILENLLKLGEDPEAVREV<br>LEALSRLCIEHPETAPLAFRLIAKIMKTVKVSPEVAAVIAEVMT<br>RTLIALIEQGVPLELMAEAMEYAAEALIALAKKVPEVLPVLLEV<br>LTRVAEALLAAPEYLVPAMPALLRAAEALAEKMAGGDERTVEAA<br>VRLARVLKRAAEALQERAQETGDERLLKLAERALEVAVRLVEIA<br>KAA | Design $P2_13$-6-v5, component A, isomorphous lattice to $P2_13$-6-0 |
| p213-6-v8A | M<u>RGHHHHHHGSSG</u>ELEELKKEFEKMTSIILLLLAEITLTLAKIV | Design $P2_13$-6-v8, |

| | | |
|---|---|---|
| | AKVLKDEEEVLRLLERAKWIAKELSKQEIVDAELLKRLLERLKELIEEVELNEEQLEEALKLLKELIKVMKKVPQEDFPAVAKALDEALEALLEQAKKVGEELALEALLAKAELLKLCIERGGSEEYVEKAVRLARELKKLAEKYQKKAQETGDEELLKKAEEALEKAVELVELAKKA | component A, isomorphous lattice to $P2_13$-6-0 |
| p213-6-v11A | MRGHHHHHHGSSGIEEEIRLLLLRVLLVELRVLALEPDAPEEEAELVSKLLSTLISKYPIDNEEAELITKALARVKGRAPVLLKGILKTIIELANADNETILNLLWMATEALKAAAEAGINERLPEVLKELAKAVKVVIKNNKDIEVEVALVELVEEALKWAEKVKIDEEGVEALVRLARLLKEIAERFQERAQETGDERLLKLAERALEIAVRLVELAKKA | Design $P2_13$-6-v11, component A, isomorphous lattice to $P2_13$-6-0 |
| p213-6-v14A | MRGHHHHHHGSWSGMLLEKLAARLFGVELRLLALEGAPVEAAIALVEEFLEMGVNPRELLKVACELARELQPEDPRAAAVMAAALVTLAIAALKAGTLTEEELKEALKVIAELLKKTEDIDTETAMKLVELALELLKLVKDASLEVLKEAAEVAKEVAKLLKKLVKEGVEEAVEKAVRLARLLKEIAERFQEEAQKTGDEELLKLAEEALETAVELVEVAKEF | Design $P2_13$-6-v14, $P2_13$-6-v14-e3 and Design $P2_13$-6-v14-e10, component A |
| p213-6-v17A | MRGHHHHHHGSSGRFFAETTAEILKIIAGILYLVLKAVPNVDKELANEILKELAKLLKKLLEKYPELVPLIAEAAKELAWAAKLLKELSVETVKEFLELLKLLAKQDKELALEAAKEAAWALKKLENPEALAEGLKGLLKLLKELKDGDLVLLLEALIEAAEAAKRALELGVVEEAVEILVRLARLLKEVAEKLQKKAQETGDEELLKLAEEALEAAVEAVEIAKEA | Design $P2_13$-6-v17, component A, isomorphous lattice to $P2_13$-6-0 |
| p213-6-v23A | MRGHHHHHHGSSGEIEKKILELTIKIAKLAFVAAAIVAKTTKDKEWLDELLKLACELLKEIIKNPLITPEQLEELCEALKEMLEESSELISPETAKLILETLKEIIKNPKVTVKALENLGLAAVELLPIVPEEALEVLEEILKKLREEDLEVAAEVLLRAAEEILKILKEEKKPIAVEIAVRLARKLKELAEELQKKAQETGDEELLKVAEKLLEKAVELVEAAKEL | Design $P2_13$-6-v23, component A, isomorphous lattice to $P2_13$-6-0 |
| p213-6-v24A | MRGHHHHHHGSWSGRLLKIELTLLENLAYLAAAFKEHKEVVRRLARELRELALELAKENPEEVVERLAETLELLLKSGLLSAEGAAEFAILVCELALEVRDNEEALRRLLETLAALLERLTELVEEADEELLVRLARRVLEALRELRDADLVALARAARAAGRLALAAIERGAGEEAVELAVRLARLLKELAERLQEEAQRTGDEELLKLAEELLEEAVRLVEAAKRR | Design $P2_13$-6-v24, component A, isomorphous lattice to $P2_13$-6-0 |
| p213-6-v26A | MRGHHHHHHGSWSGEEKKKLVLKLFEINCNIVIIYSLCGIDDEELLRKLCEEIEELFKELIDDLELLTEALRELARMVKRANPANAGIVLKMALRIATYVIKKQPEAAEAVTELLRALAEATAKLLPEQALEVAPAALELARTILADPRAPVELLKAAVELLIALAEVLAKARTPEAVEALVRVARVLKEFAERLQELAQETGDEELLKLAERALEAAVRAVEIAKEL | Design $P2_13$-6-v26, component A, isomorphous lattice to $P2_13$-6-0 |
| p213-6-e3B | MEEAEETAKRLARELLRVKLLQALAELEKLLAELRKAPTPEREA | Design $P2_13$-6-v14-e3, |

| | | |
|---|---|---|
| | ALAAALAELARLAEEVAAEDPAYAREVAALAAEVAGALAEIAPHLSPEAARAIGAALGRLVALATSSPELSLEDLRAVLEALRALLLGMSPEVADAFLAGFAESMTEYVRKTGKVSEHLVLMAEFLLHLALVLGASRETQLAVLELHLLAKLALLATGSWSGLEHHHHHH | component B, expanded lattice of $P2_13$-6-0 |
| p213-6-e10B | MEEAEETAKRLAQELLRVKLLQLLAEAEKVAEKLKKDITPENVKTLLETLEKIEEALEKLKPIIDKELAKKLGKAFAKIAKALAEALKALFEAGADTETLVTAVRAAMLAIALLLEMVAIAADDKELAKEAAKALAEVAKAVALGTAEVTRAALEGLEEAGKRVARYPEGAEAVAKVAREARREVLRYSLDLTLEQRLNATTVLLALTTLELEALRAEVRAGSWSGLEHHHHHH | Design $P2_13$-6-v14-e10, component B, expanded lattice of $P2_13$-6-0 |
| p213-6-0A-mScarlet2 | MRGHHHHHHGSSGDSTEAVIKEFMRFKVHMEGSMNGHEFEIEGEGEGRPYEGTQTAKLRVTKGGPLPFSWDILSPQFMYGSRAFTKHPADIPDYWKQSFPEGFKWERVMNFEDGGAVSVAQDTSLEDGTLIYKVKLRGTNFPPDGPVMQKKTMGWEASTERLYPEDVVLKGDIKMALRLKDGGRYLADFKTTYRAKKPVQMPGAFNIDRKLDITSHNEDYTVVEQYERSVARHSTGGSGGSGGSGGSSKELEAVARLQELNIELARKLLEAVARLQELNIDLVRKTSELTDEKTIREEIRKVKEESKRIVEEAEEEIRRAKAISLHIAIRALEDAARQVEEAIKRNPDNDEAVETAVRIARLLKEAAEKAQELAKRLGDPELLKDALRALEVAVRAVELAIRSNPNNEEAVRTAVRLAEELMKVARELIERAEKTGDPELLKLARRAAEMAVRAVEMAIKANPDDDKAVEAAVRLARVLKEIAERLQEEAQKRGDEELLKEAERALEVAVRAVELAKKS | Design $P2_13$-6-0, component A, fused with mScarlet2 on N terminal |
| p213-6-0A-mNeonGreen | MRGHHHHHHGSSGVSKGEEDNMASLPATHELHIFGSINGVDFDMVGQGTGNPNDGYEELNLKSTKGDLQFSPWILVPHIGYGFHQYLPYPDGMSPFQAAMVDGSGYQVHRTMQFEDGASLTVNYRYTYEGSHIKGEAQVKGTGFPADGPVMTNSLTAADWCRSKKTYPNDKTIISTFKWSYTTGNGKRYRSTARTTYTFAKPMAANYLKNQPMYVFRKTELKHSKTELNFKEWQKAFTDGGSGGSGGSGGSSKELEAVARLQELNIELARKLLEAVARLQELNIDLVRKTSELTDEKTIREEIRKVKEESKRIVEEAEEEIRRAKAISLHIAIRALEDAARQVEEAIKRNPDNDEAVETAVRIARLLKEAAEKAQELAKRLGDPELLKDALRALEVAVRAVELAIRSNPNNEEAVRTAVRLAEELMKVARELIERAEKTGDPELLKLARRAAEMAVRAVEMAIKANPDDDKAVEAAVRLARVLKEIAERLQEEAQKRGDEELLKEAERALEVAVRAVELAKKS | Design $P2_13$-6-0, component A, fused with mNeoGreen on N terminal |
| p213-6-v1A-mCitrine | MRGHHHHHHGSSGVSKGEELFTGVVPILVELDGDVNGHKFSVSGEGEGDATYGKLTLKFICTTGKLPVPWPTLVTTFGYGLMCFARYPDHMKQHDFFKSAMPEGYVQERTIFFKDDGNYKTRAEVKFEGDTLVNRIELKGIDFKEDGNILGHKLEYNYNSHNVYIMADKQKNGIKVNFKIRHNIEDGSVQLADHYQQNTPIGDGPVLLPDNHYLSYQSKLSKDPNEKRDHMVLLEFVTAAGITGGSGGSGGSGGSETIKKIIKLLSLITLLLIREAIKKVKLGELSAEEVKIIVEAAALQIELDKENAPELLEELKKLAKEVVRKIPEVVIHIARKIEELVKAAPKENIPHLEELVEELIEEVEREGVLTLELVIEIGRAVLTLLKKGSSERTVELAVRLARILKRMAEELQKRAQETGDERLLKKAEEALEVAVELVEAAKEA | Design $P2_13$-6-v1, component A, fused with mCitrine on N terminal |

| | | |
|---|---|---|
| p213-6-v2A-mScarlet3 | MRGHHHHHHGSSGDSTEAVIKEFMRFKVHMEGSMNGHEFEIEGE<br>GEGRPYEGTQTAKLRVTKGGPLPFSWDILSPQFMYGSRAFTKHP<br>ADIPDYWKQSFPEGFKWERVMNFEDGGAVSVAQDTSLEDGTLIY<br>KVKLRGTNFPPDGPVMQKKTMGWEASTERLYPEDVVLKGDIKMA<br>LRLKDGGRYLADFKTTYRAKKPVQMPGAFNIDRKLDITSHNEDY<br>TVVEQYERSVARHSTGGSGGSGGSGGSDLALLAKTIAEIIVILA<br>EVLVRLGALTPELAKRLAELVAKLLKAFPETAPQAIRFAEVLLE<br>AVRRDPRFLPVLVELLEAMAEEVERLTPEDKRRLARVARAALEA<br>AREQGAPLEVLLRIERVLARFLLALVEDGDEEAVEEAVRVARRL<br>KELAEELQKRAQETGDEELLKLAEEALEEAVRLVEAAKRF | Design $P2_13$-6-v2, component A, fused with mScarlet3 on N terminal |
| p213-6-v5A-mGrape3 | MRGHHHHHHGSSGVSKGEENNMAVIKEFMRFKTHMEGSVNGHEF<br>EIEGKGEGRPYEGTQTAKLKVTKGGPLPFAWDILSPQMMYGSKA<br>YVKHPPDIPDYMKLSFPEGFKWERVMNFEDGGVVTVTQDSSLQD<br>GEFIYKVKLHGTNFPSDGPVMQKKTMGWEASSERLYPEDGALKG<br>EVKMRLKLKDGGHYDAEVKTTYMAKKPVQLPGAYKLDYKLDITS<br>HNEDYTIVEQYERAEGRHSTGGSGGSGGSGGSMEARLRAILLAL<br>ILENLLKLGEDPEAVREVLEALSRLCIEHPETAPLAFRLIAKIM<br>KTVKVSPEVAAVIAEVMTRTLIALIEQGVPLELMAEAMEYAAEA<br>LIALAKKVPEVLPVLLEVLTRVAEALLAAPEYLVPAMPALLRAA<br>EALAEKMAGGDERTVEAAVRLARVLKRAAEALQERAQETGDERL<br>LKLAERALEVAVRLVEIAKAA | Design $P2_13$-6-v5, component A, fused with mGrape3 on N terminal |
| p213-6-v14A-mOrange | MRGHHHHHHGSSGVSKGEENNMAIIKEFMRFKVRMEGSVNGHEF<br>EIEGEGEGRPYEGFQTAKLKVTKGGPLPFAWDILSPQFTYGSKA<br>YVKHPADIPDYFKLSFPEGFKWERVMNFEDGGVVTVTQDSSLQD<br>GEFIYKVKLRGTNFPSDGPVMQKKTMGWEASSERMYPEDGALKG<br>EIKMRLKLKDGGHYTSEVKTTYKAKKPVQLPGAYIVGIKLDITS<br>HNEDYTIVEQYERAEGRHSTGGSGGSGGSGGSMLLEKLAARLFG<br>VELRLLALEGAPVEAAIALVEEFLEMGVNPRELLKVACELAREL<br>QPEDPRAAAVMAAALVTLAIAALKAGTLTEEELKEALKVIAELL<br>KKTEDIDTETAMKLVELALELLKLVKDASLEVLKEAAEVAKEVA<br>KLLKKLVKEGVEEAVEKAVRLARLLKEIAERFQEEAQKTGDEEL<br>LKLAEEALETAVELVEVAKEF | Design $P2_13$-6-v14, component A, fused with mOrange on N terminal |
| p213-6-v26A-mCherry2 | MRGHHHHHHGSSGVSKGEEDNMAIIKEFMRFKVHMEGSVNGHEF<br>EIEGEGEGRPYEGTQTAKLKVTKGGPLPFAWDILSPQFMYGSKA<br>YVKHPADIPDYLKLSFPEGFNWERVMNFEDGGVVTVTQDSSLQD<br>GEFIYKVKLRGTNFPSDGPVMQCRTMGWEASTERMYPEDGALKG<br>EIKQRLKLKDGGHYDAEVKTTYKAKKPVQLPGAYNVDIKLDILS<br>HNEDYTIVEQYERAEGRHSTGGSGGSGGSGGSEEKKKLVLKLFE<br>INCNIVIIYSLCGIDDEELLRKLCEEIEELFKELIDDLELLTEA<br>LRELARMVKRANPANAGIVLKMALRIATYVIKKQPEAAEAVTEL<br>LRALAEATAKLLPEQALEVAPAALELARTILADPRAPVELLKAA<br>VELLIALAEVLAKARTPEAVEALVRVARVLKEFAERLQELAQET<br>GDEELLKLAERALEAAVRAVEIAKEL | Design $P2_13$-6-v26, component A, fused with mCherry2 on N terminal |
| i213-0 (i213-w3-trim) | MSAEKLIELLERSLRKQEQPLTERERELLALAQKVLVLILKAAE<br>LALQGNKEEAEKVLRRAAEEIREATRLAEEAAKNSETPEEALEA | Design $I2_13$-0 by trimming interface |

| | | |
|---|---|---|
| | AEILVLLIEALIAIAELLQLQGNKEEAEKVLREATELIKRVTEL<br>LEKIAKNADTPELAERAAELLQRLIRLLEEIARLLKEQGNKEEA<br>RKVQEEAKELKKRVAELLAELAKEVAREVGDPELIKLAEAARKL<br>GDAEAAQAILEAAKKAKEAKERGDELKIALARLEAALALLKASL<br>RLLERSLEELEKNPSEDALVENNRLNVENNKLIVKVLEIIAEVL<br>KLNAKAVGSWSGLEHHHHHH | |
| i213-1-1 | MTLRVRIENATLEQAAKIIAKAQEVAAAEGGTTDYEFKKGTLTV<br>TMANIQGKAAAEVLAFAIEMALPNSELAREAKELAEEAKKSTDE<br>EQEKVVFLALAIVLQLPDTELAEIALRVAKLAVEADDQEALKKA<br>YEALQRVQDKPNSKEAIKALLEAALAALEVALRKLKKSLDELER<br>SLEELEKNPSEDALVENNRLNVENNKIIVKVLEIIAIVLKINAL<br>LGSWSGLEHHHHHH | Design $I2_13$-1-1 by extracted interface |
| i213-1-11 | MTRTKTFEEVSLEAAAEIIKKAQEVAAREGGTTDYEFKKGTLTV<br>TMANISEEAKKEVEAFAAIVKAKASPDPKLQELAKEAEEIMQLV<br>REAREKQRQGDKEEAEKVLRKATELIKKAFVKAAEVALRSSDPK<br>QAKLAAELMKALVMLLEIIAKLLEEEGNEDEAEKVKKAAEVLEA<br>LAELLRALLELREMLRKLKESLEELKKNPSEDALVRNNELIVEV<br>LRVIVEVLEIILKVLALLAALVGSWSGLEHHHHHH | Design $I2_13$-1-11 by extracted interface |
| i213-1-22 | MTSTVRIEGATLEAAAKIIAKAQEVAAAEGGTTDYEFKKGTLTV<br>TLANISEEAEAEVELYAAELMAKSPDPRALKVAIKLAEKAVRKN<br>PGDERAERAVEVIEEAAKLLLKSPDPEAQKEALKALLKVLEYAL<br>RKLKKSLDELERSLEELEKNPSEDALVENNRLNVENNKIIVEVL<br>EIIAKVLKAAAKVGGSWSGLEHHHHHH | Design $I2_13$-1-12 by extracted interface |
| p213-6-0A-tp1 | MRGHHHHHHGSWSGSKELEAVARLQELNIELARKLLEAVARLQE<br>LNIDLVRKTSELTDEKTIREEIRKVKEESKRIVEEAEEEIRRAK<br>AISLHIAIRALEDAARQVEEAIKRNPDNDEAVETAVRIARLLKE<br>AAEKAQELAKRLGDPELLKDALRALEVAVRAVELAIRSNPDDDK<br>AVEAAVRLARVLKEIAERLQEEAQKRGDEELLKEAERALEVAVR<br>AVELAKKS | Design $P2_13$-6-0-tp1, component A, one helical repeat trimmed from p213-6-0A |
| p213-6-0A-ep1 | MRGHHHHHHGSWSGSKELEAVARLQELNIELARKLLEAVARLQE<br>LNIDLVRKTSELTDEKTIREEIRKVKEESKRIVEEAEEEIRRAK<br>AISLHIAIRALEDAARQVEEAIKRNPDNDEAVETAVRIARLLKE<br>AAEKAQELAKRLGDPELLKDALRALEVAVRAVELAIRSNPNNEE<br>AVRTAVRLAEELMKVARELIERAEKTGDPELLKLARRAAEMAVR<br>AVEMAIKANPNNEEAVRTAVRLAEELMKVARELIERAEKTGDPE<br>LLKLARRAAEMAVRAVEMAIKANPDDDKAVEAAVRLARVLKEIA<br>ERLQEEAQKRGDEELLKEAERALEVAVRAVELAKKS | Design $P2_13$-6-0-ep1, component A, one helical repeat extended from p213-6-0A |
| p213-6-11A_tp1 | MRGHHHHHHGSWSGSEQEEIVNRLQRLNIELARQLLEAVARLQE<br>LNIDLVRKTSELTDEKTIREEIRKVKEESKRIVEEAEIRIRAAE<br>VISEALALQAELPSEEAKKAVEEIEEAAKAAERAAEKGQTEVAK<br>QALELLKRAIEEAKRKRSEEALRKVERIARAAKLIARAAELKAE<br>AERLIKKARKTGDPELLRKALEALEEAVRAAEEAIKTNPDNDEA<br>VETAVRLARELKKVAEELQERAQKTGDEELLKLAERALEVAVRA | Design $P2_13$-6-11-tp1, component A, one helical repeat trimmed from p213-6-11A |

| | | |
|---|---|---|
| | VELAKKS | |
| p213-6-11A_ep1 | MRGHHHHHHGSWSGSEQEEIVNRLQRLNIELARQLLEAVARLQELNIDLVRKTSELTDEKTIREEIRKVKEESKRIVEEAEIRIRAAEVISEALALQAELPSEEAKKAVEEIEEAAKAAERAAEKGQTEVAKQALELLKRAIEEAKRKRSEEALRKVERIARAAKLIARAAELKAEAERLIKKARKTGDPELLRKALEALEEAVRAAEEAIKTNPDDDMAVKMAVELARELKKVAEELQERAKKTGDPELLKLALRALEVAVRAVELAIKSNPDDDMAVKMAVELARELKKVAEELQERAKKTGDPELLKLALRALEVAVRAVELAIKSNPDNDEAVETAVRLARELKKVAEELQERAQKTGDEELLKLAERALEVAVRAVELAKKS | Design *P*2$_1$3-6-11-ep1, component A, one helical repeat extended from p213-6-11A |
| p213-14-17A_tp1 | MRGHHHHHHGSWSGSEEQQRVAALQKLNIELARALLEAVARLQELNIDLVRKTSELTDEKTIREEIRKVKEESKKIVKAAEIIIRIAKYASIAINKNGSDEARQAVKEIARLAKEALEEGCCDAAKFAIRALELLARVFDGSDVQRLAEEAIEKIAETAARNGGDETAKEAARALKRLAEELIKKARKTGDPELLRQALRALEAAARAAELAIRFNPDNDEAVETAVRLARELKKVAEELQERAKKTGDEELLKLAERALEVAVRAVELAKKS | Design *P*2$_1$3-14-17-tp1, component A, one helical repeat trimmed from p213-14-17A |
| p213-14-17A_ep1 | MRGHHHHHHGSWSGSEEQQRVAALQKLNIELARALLEAVARLQELNIDLVRKTSELTDEKTIREEIRKVKEESKKIVKAAEIIIRIAKYASIAINKNGSDEARQAVKEIARLAKEALEEGCCDAAKFAIRALELLARVFDGSDVQRLAEEAIEKIAETAARNGGDETAKEAARALKRLAEELIKKARKTGDPELLRQALRALEAAARAAELAIRFNPDDDEAVELAVRLARELKKVAEELQERAKKTGDPELLKLALRALEVAVRAVELAIKSNPDDDEAVELAVRLARELKKVAEELQERAKKTGDPELLKLALRALEVAVRAVELAIKSNPDNDEAVETAVRLARELKKVAEELQERAKKTGDEELLKLAERALEVAVRAVELAKKS | Design *P*2$_1$3-14-17-ep1, component A, one helical repeat extended from p213-14-17A |
| p213-14-20A_tp1 | MRGHHHHHHGSWSGSYQLEIVARLQELNIRLARQLLEAVARLQELNIDLVRKTSELTDEKTIREEIRKVKEESKRIVEEAELLIEAARVASEILTLAAKAEEEGNPELRDSARKLYQALKEALEIAKRAGEPEVIREVTREAEEAAKVIKEAIEAEKQGDRERHKKAQVRVLILIAEILKKAAEAAIKKARKTGDPELLRKALELLEKAARAAEEAIKRNPDNDEAVETAVRLARELKKVAEELQERAKKTGDEELLKLAERALEVAVRAVELAKKS | Design *P*2$_1$3-14-20-tp1, component A, one helical repeat trimmed from p213-14-20A |
| p213-14-20A_ep1 | MRGHHHHHHGSWSGSYQLEIVARLQELNIRLARQLLEAVARLQELNIDLVRKTSELTDEKTIREEIRKVKEESKRIVEEAELLIEAARVASEILTLAAKAEEEGNPELRDSARKLYQALKEALEIAKRAGEPEVIREVTREAEEAAKVIKEAIEAEKQGDRERHKKAQVRVLILIAEILKKAAEAAIKKARKTGDPELLRKALELLEKAARAAEEAIKRNPDDDKAVEMAVRLARELKKVAEELQERAKKTGDPELLKLALRALEVAVRAVELAIKSNPDDDKAVEMAVRLARELKKVAEELQERAKKTGDPELLKLALRALEVAVRAVELAIKSNPDNDEAVETAVRLARELKKVAEELQERAKKTGDEELLKLAERALEVAVRAVELAKKS | Design *P*2$_1$3-14-20-ep1, component A, one helical repeat extended from p213-14-20A |
| p213-1-0- | MRGHHHHHHGSWSGDDELEAVAELQELNIQLAKKLLEAVARLQE | Design *P*2$_1$3-1-0-MPNN- |

| | | |
|---|---|---|
| MPNN-62A | LNIDLVRKTSELTDEKTIREEIRKVKEESKRIVEEAEEEIRRAK QISKAIEEMARLRREAENAETAREAREAAERLIELARELKEMGD SERAREVLEEAERLIEEAFRWAREEAANAETAREALEAARELVR MVELLIEIARLLKEIGDSEEARRVLEKAARFIEEVRRLLEEIAR NAETTEEAREAAELLRELTRLLREIAELLEEI | 62, component A, redesigned by ProteinMPNN |
| p213-1-0-MPNN-62B | MSARAREVREKAEEFEREVARLELEIALDELEKALQELREMLRK LKESLEELKKNPSEDALVRNNELIVEVLRVIVEVLSIIAKVLKL NAKLAGSWSGLEHHHHHH | Design $P2_13$-1-0-MPNN-62, component B, redesigned by ProteinMPNN |
| p213-1-0-MPNN-94A | MRGHHHHHHGSWSGDDELEAVAELQELNIQLAKKLLEAVARLQE LNIDLVRKTSELTDEKTIREEIRKVKEESKRIVEEAEEEIRRAK QISKAIEEMAKLREQAKNAKTAKEGRDAARKMIEIAKKLKEMGD EERAREVLEEAAEIIEQQFKRAKEEAKNAKTAKEALEAAREMVE MVRLMIEIAKLLKEIGDEEEARKVLKEAEKRIEEVKELLEEIAK NAETDEEAREAAELMRELIRLLREIAELLDEI | Design $P2_13$-1-0-MPNN-94, component A, redesigned by ProteinMPNN |
| p213-1-0-MPNN-94A | MSEEAERVRKKAEEFEKETARLELEITLDELEKALQELREMLRK LKESLEELKKNPSEDALVRNNELIVEVLRVIVEVLSIIAKVLKL NAKLAGSWSGLEHHHHHH | Design $P2_13$-1-0-MPNN-94, component B, redesigned by ProteinMPNN |
| p213-12-7A_S1 | MRGHHHHHHGSWSGSKELEAVARLQRLNIELARKLLEAVARLQE LNIDLVRKTSELTDEKTIREEIRKVKEESKRIVLRAKAQILAAK AQSAIIAAEAALTAGDPETAREAVREALELVEQLRKLAKKAGDK EALEAAAELARQVAKVAREVGSPETALEALKVAAEAAKESGNKE EAEKILREATELIKRATEELEKRAKNARTSEEAKRAAEKLKKLI EMLKEIAKLLEEAGNEDEAEKVKEEAKE | Design $P2_13$-12-7 with shielded crystal contact, 13 aa shield,, component A |
| p213-12-7A_S2 | MRGHHHHHHGSWSGSKELEAVARLQRLNIELARKLLEAVARLQE LNIDLVRKTSELTDEKTIREEIRKVKEESKRIVLRAKAQILAAK AQSAIIAAEAALTAGDPETAREAVREALELVEQLRKLAKKAGDK EALEAAAELARQVAKVAREVGSPETALEALKVAAEAAKESGNKE EAEKILREATELIKRATEELEKRAKNARTSEEAKRAAEKLKKLI EMLKEIAKLLEEAGNEDEAEKVKEEAKELERR | Design $P2_13$-12-7 with shielded crystal contact, 17 aa shield,, component A |
| p213-12-7A_S3 | MRGHHHHHHGSWSGSKELEAVARLQRLNIELARKLLEAVARLQE LNIDLVRKTSELTDEKTIREEIRKVKEESKRIVLRAKAQILAAK AQSAIIAAEAALTAGDPETAREAVREALELVEQLRKLAKKAGDK EALEAAAELARQVAKVAREVGSPETALEALKVAAEAAKESGNKE EAEKILREATELIKRATEELEKRAKNARTSEEAKRAAEKLKKLI EMLKEIAKLLEEAGNEDEAEKVKEEAKELERRVTK | Design $P2_13$-12-7 with shielded crystal contact, 20 aa shield,, component A |

**Supplementary Table S2. Rosetta metrics of validated crystal contact modules, compared with previously designed de novo nanocage interfaces.** SC: shape complementarity; SASA: solvent-accessible surface area; CMS: contact molecular surface. Computational design models were used for all calculations.

| Interfaces | SC | SASA/Å$^2$ | CMS/Å$^2$ | ddG/kcal/mol |
|---|---|---|---|---|
| *P*$2_1$3-cc1 | 0.719 | 1979.014 | 468.985 | -70.781 |
| *P*$2_1$3-cc6 | 0.766 | 1593.892 | 418.532 | -55.976 |
| *P*$2_1$3-cc14 | 0.75 | 1698.306 | 415.383 | -56.23 |
| *P*$2_1$3-cc12 | 0.643 | 1903.22 | 442.005 | -59.51 |
| I$2_1$3-cc0 | 0.466 | 1885.495 | 368.343 | -49.66 |
| I$2_1$3-cc1 | 0.751 | 1115.71 | 246.068 | -43.044 |
| C3_2L6HC3-12 | 0.667 | 2113.897 | 973.44 | -113.238 |
| C3_2L6HC3-13 | 0.66 | 2142.626 | 993.547 | -102.556 |
| C3_2L6HC3-6 | 0.689 | 2149.931 | 962.823 | -109.765 |
| C3_5L6HC3-1 | 0.62 | 2083.014 | 806.043 | -102.999 |
| C3_ph192 | 0.657 | 2148.267 | 879.472 | -82.369 |
| T33-15 | 0.713 | 1355.685 | 216.105 | -54.51 |
| T32-15 | 0.536 | 1538.608 | 218.214 | -50.677 |
| O43-2 | 0.655 | 1308.395 | 218.662 | -69.685 |
| T33-ECY54 | 0.453 | 1162.437 | 235.631 | -50.809 |
| T33-ECY55 | 0.676 | 1062.419 | 175.44 | -55.302 |
| T33-ECY59 | 0.671 | 1086.636 | 179.141 | -59.631 |
| T33-ECY66 | 0.645 | 927.653 | 122.051 | -44.382 |
| T33-ECY67 | 0.594 | 1532.665 | 271.135 | -77.667 |
| O43-UWN38 | 0.679 | 987.774 | 145.462 | -41.749 |
| O43-UWN453 | 0.661 | 1185.968 | 216.101 | -55.634 |
| O43-ZL1 | 0.717 | 1193.553 | 243.974 | -57.413 |
| O43-ZL7 | 0.692 | 1286.934 | 274.764 | -71.355 |
| O32-ZL4 | 0.606 | 1358.55 | 239.474 | -62.689 |

**Supplementary Table S3. Crystallization conditions of validated designs**

Crystallization buffers and sample volumes are described in the Methods section. “Mixing” indicates that the two crystallization building blocks were combined at equal monomer-equivalent concentrations in 150 mM NaCl and 25 mM Tris-HCl, pH 8.0. “Hanging drop” refers to conventional hanging-drop vapor-diffusion crystallization. “Upon elution” indicates that proteins crystallized readily during purification after IMAC elution in a buffer containing 300 mM NaCl, 25 mM Tris-HCl, pH 8.0, and 300 mM imidazole.

| Design name | Method | Protein concentration | NaCl concentration | Crystallization time | Crystal size |
|---|---|---|---|---|---|
| $P2_13$-1-0 | Mixing | 25 µM | 150 mM | 2-3 days | 20-50 µm |
| $P2_13$-1-0 | Hanging drop | 12.5 µM | 0.5 M | Overnight | ~50 µm |
| $P2_13$-1-9 | Hanging drop | 25 µM | 0.5 M | 2-3 days | 20-50 µm |
| $P2_13$-1-11 | Hanging drop | 25 µM | 0.5 M | 2-3 days | 20-50 µm |
| $P2_13$-1-21 | Hanging drop | 25 µM | 0.5 M | Overnight | ~10 µm |
| $P2_13$-1-26 | Hanging drop | 25 µM | 1 M | Overnight | 50-100 µm |
| $P2_13$-6-0 | Hanging drop | 25 µM | 3 M | Overnight | 50-100 µm |
| $P2_13$-6-1 | Hanging drop | 25 µM | 2 M | Overnight | 20-50 µm |
| $P2_13$-6-2 | Hanging drop | 25 µM | 2 M | Overnight | 20-50 µm |
| $P2_13$-6-8 | Hanging drop | 25 µM | 2 M | Overnight | 20~50 µm |
| $P2_13$-6-10 | Hanging drop | 25 µM | 3 M | Overnight | 50-100 µm |
| $P2_13$-6-11 | Hanging drop | 25 µM | 2.5 M | Overnight | 20~50 µm |
| $P2_13$-6-12 | Hanging drop | 25 µM | 2.5 M | Overnight | ~10 µm |
| $P2_13$-12-7 | Hanging drop | 25 µM | 1 M | Overnight | 20-50 µm |
| $P2_13$-14-0 | Hanging drop | 25 µM | 2.5 M | Overnight | ~50 µm |
| $P2_13$-14-1 | Hanging drop | 25 µM | 2.5 M | Overnight | 20-50 µm |
| $P2_13$-14-2 | Hanging drop | 25 µM | 2 M | Overnight | ~50 µm |
| $P2_13$-14-3 | Hanging drop | 25 µM | 2 M | Overnight | ~50 µm |

| | | | | | |
|---|---|---|---|---|---|
| $P2_13$-14-9 | Hanging drop | 25 µM | 3 M | Overnight | 20-50 µm |
| $P2_13$-14-10 | Hanging drop | 25 µM | 3 M | Overnight | ~50 µm |
| $P2_13$-14-17 | Hanging drop | 25 µM | 3 M | Overnight | 50-100 µm |
| $P2_13$-14-20 | Hanging drop | 25 µM | 3 M | Overnight | 50-100 µm |
| $P2_13$-6-v1 | Hanging drop | 25 µM | 2 M | Overnight | 50-100 µm |
| $P2_13$-6-v2 | Hanging drop | 25 µM | 2.5 M | Overnight | 50-100 µm |
| $P2_13$-6-v5 | Hanging drop | 25 µM | 2 M | Overnight | 20-50 µm |
| $P2_13$-6-v14 | Hanging drop | 25 µM | 1 M | Overnight | 20-50 µm |
| $P2_13$-6-v26 | Hanging drop | 25 µM | 2 M | Overnight | 10-20 µm |
| $P2_13$-6-v14e3 | Hanging drop | 25 µM | 1.5 M | Overnight | ~10 µm |
| $P2_13$-6-v14e10 | Hanging drop | 25 µM | 2 M | Overnight | ~10 µm |
| $I2_13$-0 | Hanging drop | 25 µM | 0.5 M | 2-3 days | 10-20 µm |
| $I2_13$-0 | Hanging drop | 40 µM | 0.5 M (with 10% glycerol) | A week | 100-200 µm |
| $I2_13$-1-1 | Hanging drop | 80 µM | 0.5 M | Overnight | 20-50 µm |
| $I2_13$-1-11 | Hanging drop | 25 µM | 0.5 M | Overnight | 20-50 µm |
| $I2_13$-1-22 | Upon elution | —— | 0.3 M | In minutes | ~10 µm |

### Supplementary Table S4. Crystal structure data collection and refinement statistics

| | p213-14B (PDB ID: pdb_000037TD) |
|---|---|
| **Data collection** | |
| Space group | *C 2* |
| Cell dimensions | |
| *a*, *b*, *c* (Å) | 85.62, 49.30, 76.99 |
| α, β, γ (°) | 90, 90.04, 90 |
| Resolution (Å) | 42.81 - 2.7 (3.09 - 2.7) |
| $R_{merge}$ | 0.31 (0.62) |
| $I / \sigma I$ | 7.06 (2.44) |
| Completeness (%) | 97.21 (97.92) |
| Redundancy | 3.5 (3.5) |
| $CC_{1/2}$ | 0.925 (0.681) |
| **Refinement** | |
| Resolution (Å) | 42.81 - 2.7 (3.09 - 2.7) |
| No. reflections | 8760 (2913) |
| $R_{work}$ / $R_{free}$ | 0.2771 (0.3130) /0.3139 (0.3855) |
| No. atoms | |
| Protein | 2061 |
| *B*-factors | |
| Protein | 68 |
| R.m.s. deviations | |
| Bond lengths (Å) | 0.003 |
| Bond angles (°) | 0.440 |

*Single Crystal used for each data collection. *Values in parentheses are for highest-resolution shell.
